# Circadian PERIOD proteins regulate TC-DSB repair through anchoring to the nuclear envelope

**DOI:** 10.1101/2023.05.11.540338

**Authors:** Benjamin Le Bozec, Laure Guitton-Sert, Sarah Collins, Anne-Laure Finoux, Charlotte Payrault, Jessica Frison, Romane Carette, Vincent Rocher, Emmanuelle Guillou, Coline Arnould, Aude Guénolé, Marion Aguirrebengoa, Ikrame Lazar, Aline Marnef, Philippe Frit, Patrick Calsou, Thomas Mangeat, Nadine Puget, Gaëlle Legube

## Abstract

Repair of DNA Double-Strand Breaks (DSBs) produced in transcriptionally active chromatin occurs through a poorly characterized pathway called Transcription-Coupled DSB repair (TC-DSBR). Here, using a screening approach scoring multiple outputs in human cells, we identified proteins from the PERIOD complex, a key module ensuring circadian oscillations, as novel TC-DSBR players. We show that the core PER complex protein PER2 is recruited at TC-DSBs and that it contributes to the targeting of TC-DSBs at the nuclear envelope (NE). At the NE, SUN1 and the Nuclear Pore Complex (NPC) act as docking sites for TC-DSBs and TC-DSB anchoring fosters RAD51 assembly. Impaired DSB localization at the NE results in elevated DSB clustering and translocation rate. In agreement, the circadian clock regulates TC-DSB anchoring to the NE, RAD51 assembly, and DSB clustering. Our study shows that DSB localization to the NPC is a conserved molecular pathway that also occurs in human cells and provides a direct link between the circadian rhythm and the response to DSBs occurring in active genes. This opens new therapeutic strategies for chemotherapies based on drugs that are inducing DSBs in active loci such as topoisomerase poisons.

---

DNA Double-Strand breaks (DSBs) are among the most toxic lesions that occur on the genome, given their potential to elicit large chromosomal rearrangements. While global genome DSB repair (GG-DSBR) mechanisms have been well characterized, a transcription-coupled arm of DSB repair (TC-DSBR) has only recently been identified^1^. Yet, physiological DSBs mostly appear in transcriptionally active loci (TC-DSBs)^2–5^ and topoisomerase II inhibitors, classically used for cancer treatment, mainly induce TC-DSBs^2,4^ emphasizing the need to better understand TC-DSBR mechanisms. TC-DSBR entails the ATM- and DNA-PK-dependent transcriptional repression of damaged genes which is necessary for the initiation of resection^6,7^. TC-DSBs are preferentially repaired by Homologous Recombination (HR) in G2^8,9^, while in G1, they are refractory to repair and cluster in large subnuclear structures^10^ that we recently identified as being a novel, DSB-induced, chromatin compartment (the D-compartment)^11^ whose formation enhances translocation frequency and checkpoint activation^11,12^. Additionally, when occurring on human ribosomal DNA, TC-DSBs are physically relocated at the nucleolar periphery where they contact invaginations of the Nuclear Envelope (NE)^13^. Whether localization to the NE also occurs when DSBs arise within RNAPII-transcribed loci is yet unknown, although NE targeting of other types of persistent DSBs and collapsed replication forks have been reported in *Drosophila* and *S. Cerevisiae*^14–19^. Altogether the TC-DSBR pathway is still poorly understood.

## Multi-output screen to identify new TC-DSBR factors

In search of novel TC-DSBR players, we performed a siRNA library screen scoring multiple outputs in the well-characterized human DIvA cell line (Fig. 1a). In this cell line, annotated DSBs are induced throughout the genome in a temporally controlled manner, thanks to a restriction enzyme fused to the ligand binding domain of the estrogen receptor (AsiSI-ER)^20^. BLESS experiments (enabling mapping and quantification of DSBs throughout the genome using streptavidin-capture and sequencing of fragments adjacent to DSBs ligated to biotinylated linkers^21^) allowed to identify 174 sites robustly cleaved *in vivo* out of the 1211 AsiSI sites annotated on the human genome (mainly due to the fact that AsiSI activity is inhibited by DNA methylation)^20,22^. A large fraction of these DSBs occur in RNAPII-enriched (transcribed) loci hereafter called TC-DSBs^20,22^. The siRNA library comprised 130 candidate proteins previously identified to be associated with the SMC1 subunit of the cohesin complex, which was previously shown to be recruited at DSBs post-DSB induction^13^. The screen was designed to quantify the effect of candidate protein depletion on 1) γH2AX foci intensity using high-content imaging, 2) cell survival using a cell proliferation assay, 3) chromosome rearrangement frequency using quantitative PCR and 4) DSB clustering using high content microscopy (Fig. 1a, Extended Data Fig. 1a). A positive control siRNA against SETX, an R-loop resolving factor previously reported to be involved in TC-DSBR^23,24^, triggered an expected increase in γH2AX foci intensity, translocation frequency and DSB clustering as well as decreased cell proliferation (Fig. 1b red label, Extended Data Fig. 1b-e). We further validated our screening dataset with the candidate protein Periphilin (PPHLN1, Fig. 1b green label, Extended Data Fig. 1b-e), an H3K9me3 reader of the HUSH complex that we previously reported to be involved in the repair of DSB occurring on ribosomal DNA^13^. In agreement with the screen results (Fig. 1b), depletion of PPHLN1 triggered an increase in γH2AX foci intensity (Extended Data Fig. 1f-g). We further used a degradable version of AsiSI-ER (AID-tagged DIvA cell line^8^) to measure clonogenic potential and repair post-DSB induction. PPHLN1 depletion increased two distinct translocation events and cell death, in agreement with our screen results (Extended Data Fig. 1h-i).

**Figure 1:**
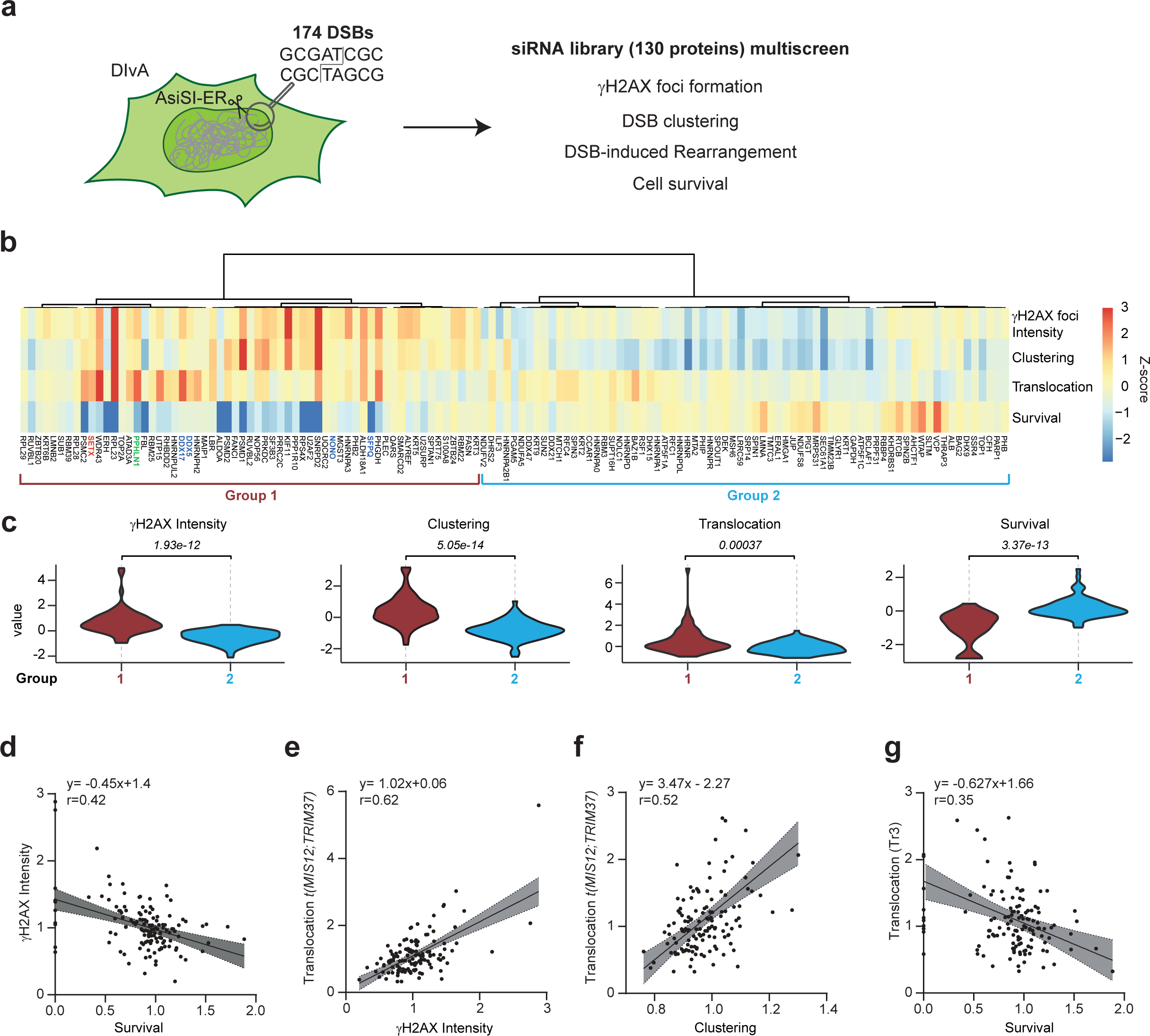
A multi-output screen for the identification of new TC-DSBR factors. **a.** Experimental scheme: DIvA cells allow to temporally induce, following treatment with 4-hydroxytamoxifen (4-OHT), DSBs at annotated positions on the human genome, among which >50% fall in active loci (TC-DSBs). TC-DSB potential interacting partners were identified through a proteomic approach^13^. 130 proteins were further subjected to a multi-output screen in AID-DIvA cells using a siRNA library, to score DSB-induced γH2AX foci intensity (using quantitative high throughput imaging), DSB clustering (γH2AX foci mean area), chromosome rearrangement (illegitimate rejoining between two TC-DSBs *t(MIS12;TRIM37)* analyzed by qPCR following DSB induction and repair) and cell survival (quantified eight days after DSB induction, using a colorimetric assay based on mitochondrial dehydrogenase activity in viable cells). **b.** Heatmap of the z-score for the 4 outputs measured (γH2AX foci intensity, Clustering, Translocation and Survival) in the screen of the 130 genes of interest ordered by hierarchical clustering (Pearson correlation distance/ward.D method) after normalization of the data to a control siRNA (see Extended Data Fig. 1b-e). A positive control (SETX) was included in the experiment (red). PPHLN1 and PER complex siRNAs are highlighted in green and blue respectively. **c.** Violin plots of scored outputs for the two groups of genes defined by the hierarchical clustering. *P*, student t-test. **d-g**. Scatter plots showing the correlations for each siRNA between gH2AX intensity and cell survival (**d**); gH2AX intensity and translocations (**e**); translocations and DSB clustering (**f**) and cell survival and translocations (**g**). The 95% confidence intervals and the coefficient of correlation (r, Pearson) are indicated.

Importantly, two-parameter correlations established that γH2AX intensity was inversely correlated with cell survival but positively correlated with translocation frequency (Fig. 1d-e), consistent with a decreased repair capacity respectively impairing cell proliferation and promoting genome rearrangements. Moreover, increased DSB clustering positively correlated with translocation frequency (Fig. 1f), and increased translocation frequency inversely correlated with cell survival (Fig. 1g) in agreement with previous work^11,12^. Altogether, results from our siRNA screen establish that TC-DSBR deficiency severely affects cell survival and further confirm the relationship between DSB clustering and genome rearrangements as recently reported^12,25^.

## The circadian clock PERIOD proteins contribute to TC-DSB repair

Hierarchical clustering identified two groups displaying statistically significant differences in clustering, translocation, γH2AX and cell survival (Fig. 1b-c). Interestingly, among the candidate proteins whose depletion gave similar phenotypes to SETX (*i.e*., increased γH2AX staining, decreased cell survival, increased chromosomal rearrangement and increased DSB clustering, Group 1, Fig. 1b-c, Extended Data Fig. 1b-e), we found NONO, SFPQ and DDX5 which all belong to the same complex, the mammalian PERIOD (PER) complex involved in the regulation of the circadian rhythm^26–29^. Similar outputs were also seen for another candidate, the DDX5-paralog DDX17, both DDX5 and DDX17 being human orthologs of the *Neurospora* PRD1 protein involved in circadian rythmicity^30^ and existing as a heterodimer^31^. The circadian clock is molecularly controlled by the BMAL1/CLOCK transcriptional activator complex, which regulates the transcription of thousands of genes. Among those, the PER-CRY transcriptional repressor can downregulate BMAL1/CLOCK activity, creating a negative transcriptional feedback loop that ensures circadian oscillations^32^. In order to directly investigate whether the core clock components of the PER complex play a role in TC-DSBR, we depleted PER1 and PER2 proteins (Extended Data Fig. 2a). As found for DDX17, NONO, DDX5 and SETX (Fig. 1b; Extended Data Fig. 1b), PER1 and PER2 knockdown increased γH2AX foci intensity following DSB induction in DIvA cells (Fig. 2a) with no major changes in cell cycle distribution (Extended Data Fig. 2b). In agreement, PER2-depleted cells also displayed elevated γH2AX levels around DSBs as detected by chromatin immunoprecipitation (ChIP) (Fig. 2b). In order to validate this finding in another, non-cancerous, cell line, we further depleted PER1 and PER2, alone or in combination, in hTERT RPE-1 cells (RPE1) stably infected with a mAID-AsiSI-ER lentiviral construct (RPE-DIvA). PER1 and/or PER2 depleted cells displayed elevated γH2AX levels post DSB induction by AsiSI (Extended Data Fig. 2c-d). Moreover, γH2AX increase following PER1/2 depletion also hold true when inducing DSB using etoposide, a Topoisomerase 2 (TOP2) inhibitor known to mainly trigger DSBs in transcribed loci and regulatory elements^2,4^ (Extended Data Fig. 2e). Of note in RPE1 cells, PER1/2 depletion also significantly increased γH2AX levels even before DSB induction using AsiSI or etoposide (Extended Data Fig. 2c-e), suggesting that PER1/2 contribute to endogenous DSB repair.

**Figure 2.**
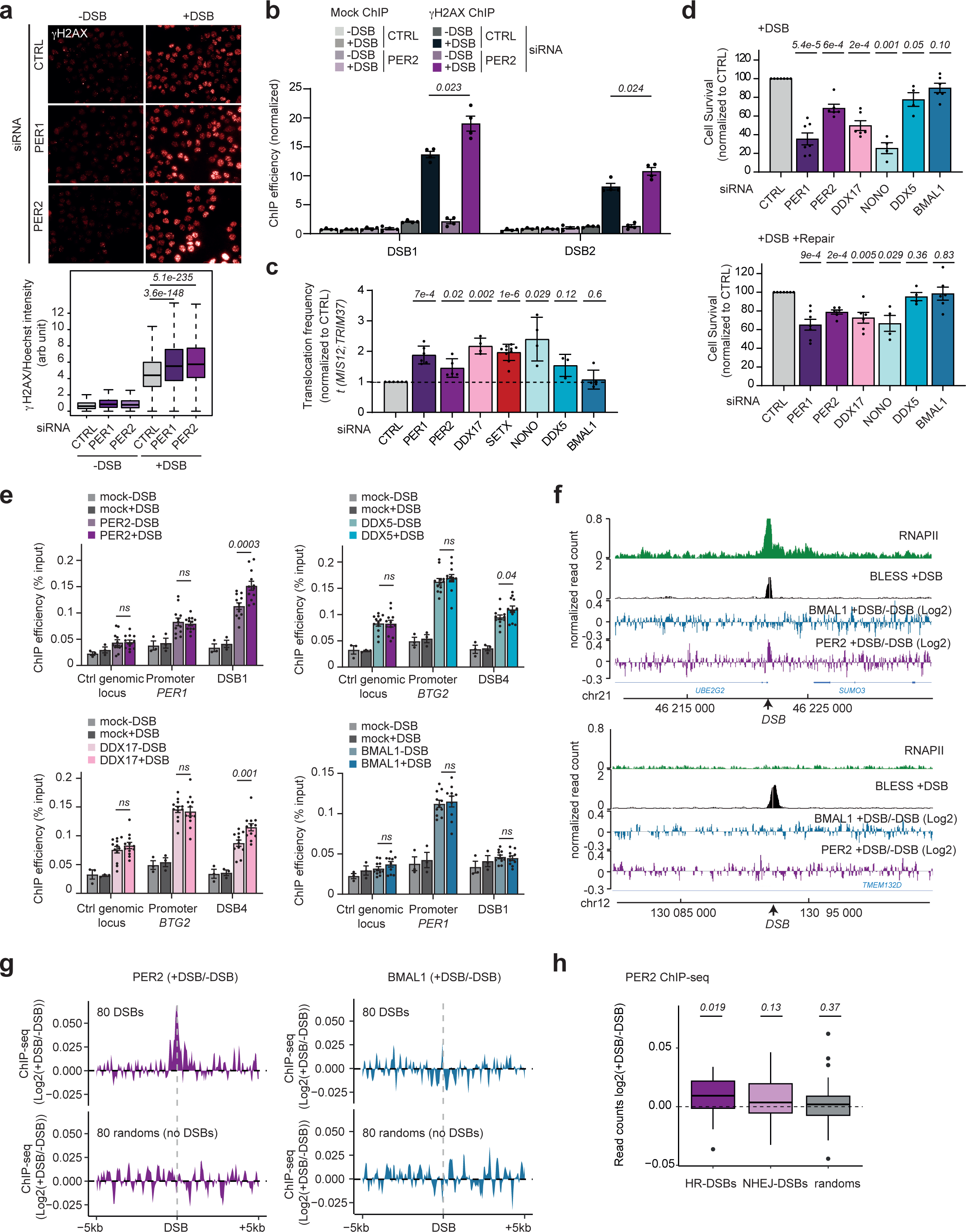
PER proteins are recruited at TC-DSBs and regulate γH2AX foci formation, translocation, and cell survival upon DSB induction **a.** γH2AX staining performed in control (CTRL), PER1 or PER2-siRNA transfected DIvA cells, before (-DSB) and after (+DSB) DSB induction. Quantification (normalized against Hoechst intensity) is shown on the bottom panel (>7000 nuclei, from a representative experiment). Center line: median; box limits: 1st and 3rd quartiles; whiskers: maximum and minimum without outliers. *P*, unpaired t-test. **b.** γH2AX ChIP performed in DIvA cells transfected with siRNA control (CTRL) or PER2 before and after DSB induction and analyzed by qPCR at two DSBs (DSB1-2). Data are normalized to a control location devoid of a DSB. Mean and SEM are shown for n=4 biological replicates. *P,* paired t-test (two-sided). **c.** *t*(*MIS12:TRIM37)* rejoining frequencies after DSB induction measured by qPCR in AID-DIvA cells transfected with siRNA CTRL, PER1 PER2, DDX17, SETX, NONO, DDX5 and BMAL1. Mean and SD (n≥4 biological replicates) are shown. *P*, paired t-test (two-sided). **d.** Clonogenic assays in AID-DIvA cells transfected with siRNA CTRL, PER1, PER2, DDX17, NONO, DDX5 or BMAL1. Upper and lower panels show the mean and SEM after 4-OHT treatment (+DSB) and after IAA treatment (+DSB+Repair) respectively (n≥4 biological replicates). *P*, paired t-test (two-sided). **e.** PER2, DDX5, DDX17 and BMAL1 ChIP efficiency (expressed as % of input immunoprecipitated) before (−DSB) and after (+DSB) DSB induction, at a control genomic locus devoid of DSB, a promoter (*BTG2* or *PER1* as indicated), and DSB1 or DSB4. Mean and SEM are shown for n≥10 replicates. *P*, paired t-test (two-sided). **f.** Genomic tracks showing RNA Polymerase II before DSB induction (RNAPII, green), BLESS after DSB induction (black), as well as BMAL1 (blue) and PER2 (violet) (log2(+DSB/−DSB)) at a TC-DSB (upper panel; chr21:460221789) and a DSB induced in a silent locus (lower panel; chr12:130091880). DSBs are indicated by arrows. **g.** Average PER2 (left) and BMAL1 (right) ChIP-seq profiles on a ±5 kb window centered on the eighty best-induced AsiSI-DSBs (top) or eighty random sites (bottom, no DSB). Data are presented in log2(+DSB/-DSB). **h.** Box plots representing Log2 (+DSB/-DSB) PER2 ChIP-seq count on a ±2kb at HR-DSB, NHEJ-DSB (n=30 in each category, among the 80 best-induced DSBs) or random sites (n=30). Center line: median; box limits: 1st and 3rd quartiles; whiskers: maximum and minimum without outliers. *P*, Wilcoxon test.

As found for DDX17, NONO, SETX and DDX5 (Fig. 1b; Fig. 2c-d; Extended Data Fig. 1c-d; Extended Data Fig. 2g-i), PER1 and PER2 depletion increased translocation frequency (Fig. 2c) and impaired cell survival following induction of TC-DSBs in DIvA cells (Fig. 2d; Extended Data Fig. 2f). Concomitant depletion of PER1 and PER2 triggered exacerbated translocation frequency and cell death following DSB induction, as compared to PER1 or PER2 depletion alone (Extended Data Fig. 2h-i). PER1/2 depletion also impaired cell survival upon treatment with the TOP2 inhibitors etoposide and doxorubicin in U2OS and RPE1 (Extended Data Fig. 2j-k).

In contrast and surprisingly, depletion of CRY2, known to function in the PER complex, did not result in an elevated translocation rate nor increased cell death (Extended Data Fig. 2h-i). Similarly, depletion of neither BMAL1 nor CLOCK transcriptional activators, alone or in combination, altered the translocation frequency or cell survival post DSB induction (Fig. 2c-d, Extended Data Fig. 2h-i). Altogether, these data show that the PERIOD proteins PER1 and PER2, as well as some other members of the PER complex (NONO, DDX17, SETX and DDX5), contribute to the response to DSBs induced in transcribed loci.

In order to determine whether the function of the PER complex is direct, we assessed the recruitment of PER complex proteins at DSBs using ChIP. PER2, DDX17 and DDX5 showed significant recruitment at a TC-DSB induced in DIvA cells (already observed for DDX5^33^), while this was not the case for BMAL1 (Fig. 2e). We further performed ChIP-seq against PER2 and BMAL1 in damaged and undamaged DIvA cells. Both proteins displayed the expected pattern on the genome (see examples Extended Data Fig. 3a). Indeed, in absence of DSB induction, BMAL1 was enriched on genomic loci previously identified by BMAL1 ChIP-seq^34^ (Extended Data Fig. 3b) and PER2 accumulated at promoters (Extended Data Fig. 3c) as expected for a transcriptional repressor and from previous studies^35^. Moreover, BMAL1-bound genes displayed significant enrichment for the “rhythmic process” GO term (Extended Data Fig. 3d). Post-DSB induction, we interestingly observed that PER2 accumulated at TC-DSBs (see an example Fig. 2f top panel, purple track) but not at a silent locus (Fig. 2f bottom panel), despite equivalent cleavage (BLESS tracks). In contrast, BMAL1 showed no accumulation at DSBs (Fig. 2f, blue tracks). On average, PER2 was significantly targeted on ∼2kb around DSBs, while BMAL1 was not (Fig. 2g, Extended Data Fig. 3e). In agreement with our above data showing that PER1/2 depletion triggers increased γH2AX even before DSB induction (Extended Data Fig. 2c-e), we also observed a significant enrichment of PER2 at previously determined endogenous DSB hotspots^11^ (Extended Data Fig. 3f). We previously identified a subset of equivalently cleaved DSBs in DIvA cells with two distinct repair behaviors: the Non-Homologous End-Joining (NHEJ)-prone DSBs and the HR-prone DSBs with the latter having a tendency to locate within transcriptionally active loci (*i.e*. TC-DSBs)^8,22^. Importantly, PER2 preferentially accumulates at HR-prone TC-DSBs compared to NHEJ-prone DSBs (*i.e.* occurring in non-transcribed regions) (Fig. 2h). Altogether these data show that the PER complex is recruited at HR-repaired, TC-DSBs, suggesting a direct role of the PER complex in TC-DSB repair.

## TC-DSBs are targeted to the nuclear envelope via a PER2-dependent mechanism

Of interest, the nuclear envelope (NE) contributes to the regulation of the circadian clock^36^ and PER proteins have been involved in the physical targeting of clock-regulated genes to the NE in *Drosophila*^37^. On another hand, persistent DSBs^14,17,38–40^, sub-telomeric DSBs^41,42^, dysfunctional telomeres^43,44^ and arrested replication forks^18,19^ have been previously found to be relocated at the NE in yeast as were stressed replication forks in mammals^45^ and heterochromatic (HC) DSBs in *Drosophila* in order to complete HR^15,46^. While in mammalian cells, physical relocation of DSBs to the NE has not yet been documented, we previously observed that ribosomal DNA DSBs can contact NE invaginations inside the nucleoplasm^13^. Hence, we set out to determine whether TC-DSBs could be targeted to the NE. As a first approach, we performed fixed and live super-resolution imaging using Random Illumination Microscopy (RIM)^47^. We observed that γH2AX foci can establish close contacts with LaminB1 filaments (Extended data Fig. 4a). Live imaging after 1 hour of DSB induction in DIvA cells expressing 53BP1-GFP and mCherry-LaminB1, showed that such relocalization of DSBs to the nuclear lamina were fast (<60s) and transient (Fig. 3a and Supplementary Videos 1-4). Contacts between repair foci and the nuclear lamina were further confirmed by Proximity Ligation Assay (PLA) performed between LaminB1 and 53BP1 (Fig. 3b) or between γH2AX and LaminB1 (Extended Data Fig. 4b). DSBs induced by etoposide in U2OS cells also displayed interactions with LaminB1 (Extended data Fig. 4c). Moreover, we could recapitulate DSB/LaminB1 interactions in RPE1 primary cells subjected to AsiSI or etoposide induced DSBs (Extended data Fig. 4d-e). In agreement, ChIP against LaminB1 (Extended Data Fig. 4f) also revealed increased LaminB1 occupancy at TC-DSBs in contrast to a DSB induced in a silent locus and repaired by NHEJ, in DIvA cells post-DSB induction (Fig. 3c). Taken together, these data suggest that TC-DSBs can be physically relocated at the NE in human cells.

**Figure 3:**
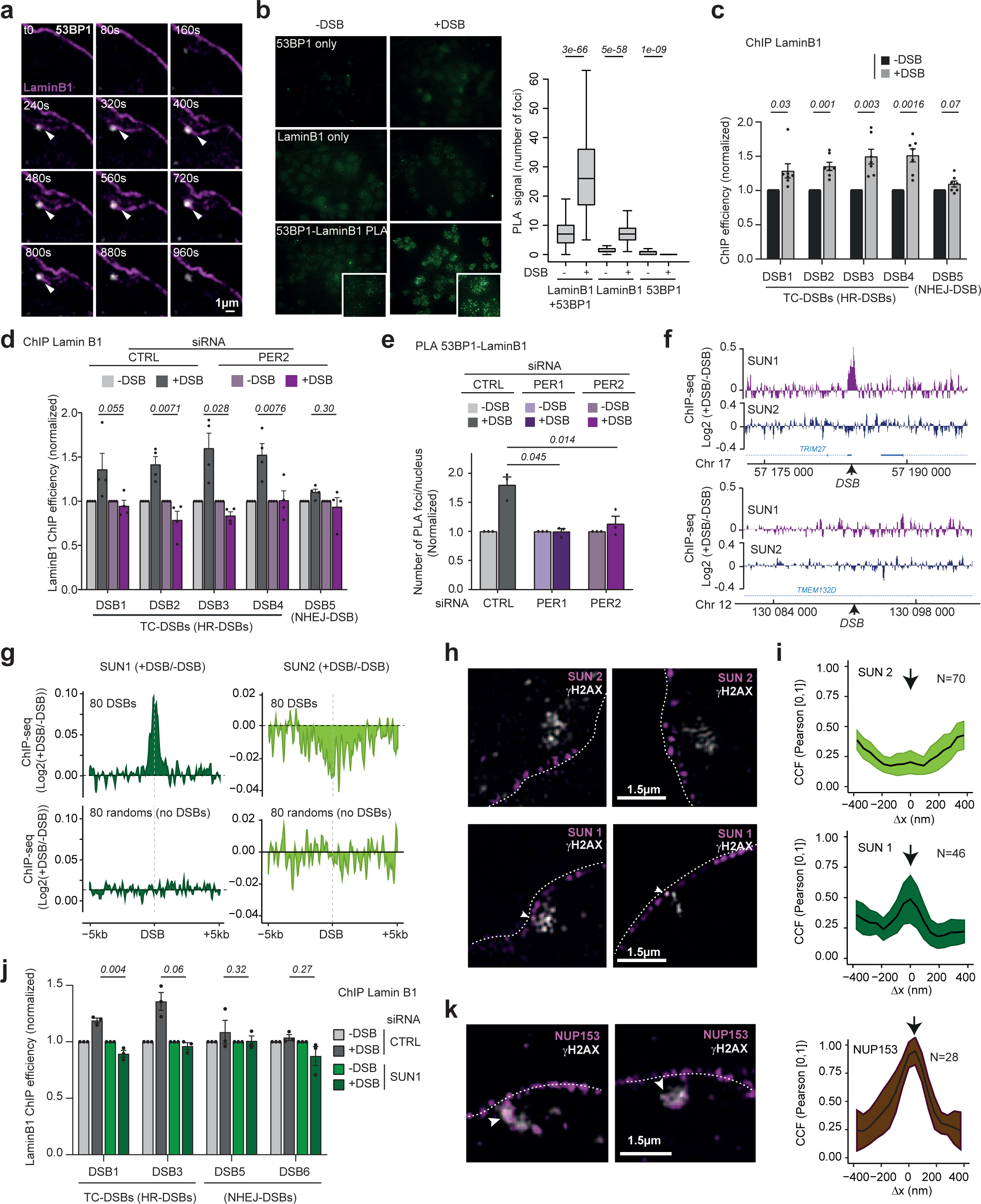
TC-DSBs are targeted to the nuclear envelope via a PER2-dependent mechanism **a.** Super-resolution live imaging performed using Random Illumination microscopy (RIM) in DIvA cells expressing 53BP1-GFP and mCherry-LaminB1, starting >1h after DSB induction. Arrows show the establishment of a contact between a focus and nuclear lamina. Images were acquired every 40s. **b.** Proximity Ligation Assay (PLA) performed using either 53BP1 antibody (top panels), Lamin B1 antibody (middle panels) or both 53BP1 and LaminB1 antibodies (bottom panels) before (-DSB) and after DSB (+DSB) induction in DIvA cells. PLA quantification is shown on the right panel. An average of 190 cells were analyzed per condition. Center line: median; box limits: 1st and 3rd quartiles; whiskers: maximum and minimum without outliers. *P*, Wilcoxon test. **c.** LaminB1 ChIP-qPCR before (−DSB) and after (+DSB) DSB induction, at TC-DSBs (HR-DSBs, DSB1-4) and one DSB induced in a silent locus and repaired by NHEJ (DSB5). Data are normalized to a control location devoid of a DSB. Mean and SEM are shown for n=7 biological replicates. *P*, paired t-test (two-sided). **d.** LaminB1 ChIP-qPCR before (−DSB) and after (+DSB) DSB induction in control and PER2 siRNA-depleted DIvA cells, at TC-DSBs (DSB1-4, HR-DSBs) and a DSB in a silent locus (DSB5, NHEJ-DSBs). Data are normalized to a control location devoid of DSB. Mean and SEM are shown for n=4 biological replicates. *P*, paired t-test (two-sided). **e.** Number of PLA foci (53BP1-LaminB1) in CTRL, PER1 or PER2-siRNA transfected DIvA cells as indicated, before (−DSB) and after (+DSB) DSB induction. Data were normalized to the untreated samples. Mean and SEM (n=3 biological replicates) are shown. *P*, paired t-test (two-sided). **f.** Genomic tracks of SUN1(purple) and SUN2 (dark blue) ChIP-seq (log2(+DSB/-DSB)) at a TC-DSB (upper panel; chr17:57184296) and a DSB induced in a silent locus (lower panel; chr12:130091880). **g.** Average profiles of SUN1 (left) and SUN2 (right) on a ±5 kb window centered on the eighty best-induced AsiSI-DSBs (top) or eighty random sites (bottom, no DSB). Data are presented as log2(+DSB/-DSB). **h.** Magnifications of SUN2 (top panels) and SUN1 (bottom panels) staining in nuclear envelope (dotted line) together with γH2AX foci in DIvA cells after DSB induction. Arrows indicate sites of colocalization. **i.** Cross Correlation Function (CCF) plots between γH2AX foci and NE-embedded SUN2 (top panel, N=70 γH2AX foci) or SUN1 (bottom panel, N=46 γH2AX foci). CCF of bicolor 3D RIM images was determined by plotting the value of Pearson’s correlation coefficient P[0.1] against Δx for each voxel of the image (see methods). The arrows show a non-random exclusion between γH2AX and SUN2 (dip at Δx=0) or a non-random overlap between the signals of SUN1and γH2AX (peak at Δx=0). **j.** LaminB1 ChIP-qPCR before (−DSB) and after (+DSB) DSB induction in control and SUN1 siRNA-depleted DIvA cells, at two TC-DSBs (DSB1 and DSB3, HR-DSBs) and two DSBs in silent loci (DSB5 and DSB6, NHEJ-DSBs). Data are normalized to a control location devoid of DSB. Mean and SEM are shown for n=3 biological replicates. *P*, paired t-test (two-sided). **k.** Left panel: Magnification of NUP153 staining in nuclear envelope (dotted line) together with γH2AX foci in DIvA cells after DSB induction. Arrows indicate sites of colocalization. Right panel: CCF plot as above between γH2AX foci and NPC protein NUP153 (N=28 γH2AX foci) showing a non-random overlap between both signals (peak at Δx=0).

**Figure 4.**
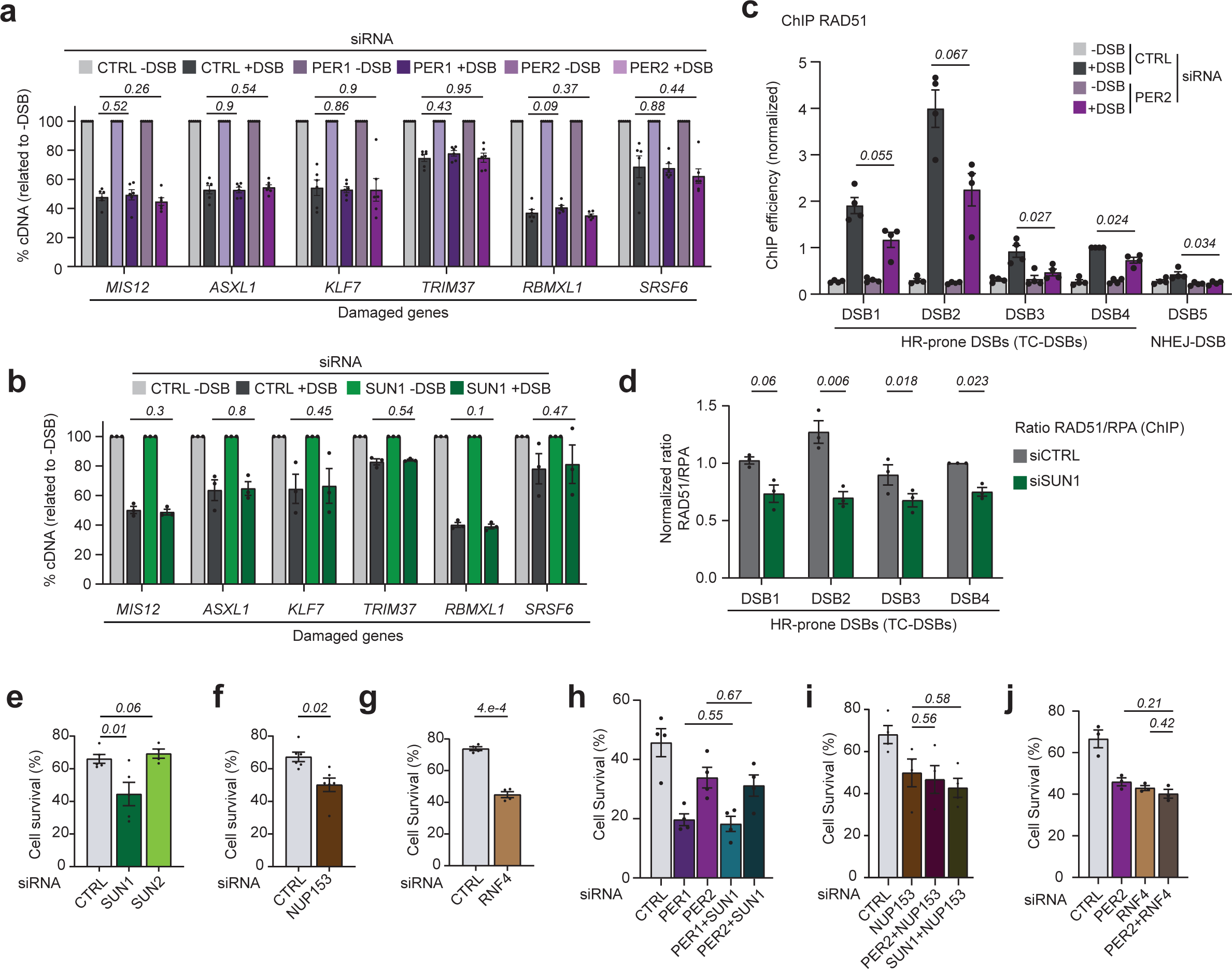
PER2 and SUN1-dependent anchoring at the Nuclear Pore Complex promotes Homologous Recombination repair **a.** RT-qPCR performed before (−DSB) and after (+DSB) DSB induction in control (CTRL), PER1 or PER2 siRNA-depleted DIvA cells for six genes carrying a TC-DSB (relative to untreated sample (-DSB)). Mean and SEM are shown for n=6 biological replicates. *P*, paired t-test (two-sided). **b.** RT-qPCR performed before (−DSB) and after (+DSB) DSB induction in control (CTRL) or SUN1 siRNA-depleted DIvA cells for six genes carrying a TC-DSB (relative to untreated sample (-DSB)). Mean and SEM are shown for n=3 biological replicates. *P*, paired t-test (two-sided). **c.** RAD51 ChIP-qPCR in DIvA cells transfected with control or PER2 siRNA before and after DSB induction, at four TC-DSBs (DSB1-4) and one DSB in a silent locus (DSB5). Data are normalized to a control location devoid of a DSB, and further expressed relative to data obtained at DSB4. Mean and SEM are shown for n=3 biological replicates. *P*, paired t-test (two-sided). **d.** Ratios of RAD51 over RPA ChIP-qPCR results in DIvA cells transfected with control (CTRL) or SUN1 siRNA after DSB induction, at four TC-DSBs (DSB1-4). Data are normalized to a control location devoid of a DSB, and further expressed relative to data obtained at DSB4. Mean and SEM are shown for n=3 biological replicates. *P*, paired t-test (two-sided). **e-j**. Clonogenic assays in AID-DIvA cells transfected with the indicated siRNAs. Mean and SEM of n biologically independent experiments after OHT and IAA treatment (DSB induction and repair) are shown. *P*, paired t-test (two-sided). **e**. Control (CTRL), SUN1 or SUN2 siRNAs (n=5). **f.** CTRL or NUP153 siRNAs (n=6). **g.** CTRL or RNF4 siRNAs (n=6). **h**. CTRL, PER1, PER2, and double PER1+SUN1 and PER2+SUN1 siRNAs (n=4). **i.** CTRL, NUP153, and double PER2+NUP153 and SUN1+NUP153 siRNAs (n=3). **j.** CTRL, PER2, RNF4 and double PER2+RNF4 siRNAs (n=3).

In order to evaluate the function of PER2 in targeting TC-DSB at the NE, we performed LaminB1 ChIP in DIvA cells upon PER2 siRNA. Of interest PER2 depletion abolished LaminB1 recruitment at TC-DSBs (Fig. 3d). Additionally, PER1 or PER2-depleted cells also displayed decreased 53BP1/LMNB1 PLA signal post-DSB induction when compared to PER1/2-proficient cells (siRNA CTRL) (Fig. 3e, Extended Data Fig. 4g). Similarly, PER1/2 depletion also decreased γH2AX/LMNB1 PLA signal in response to etoposide (Extended Data Fig. 4h). In order to investigate a potential cell cycle dependency for PER2-mediated DSB anchoring to the NE, we then monitored γH2AX/LaminB1 PLA signal combined with EdU incorporation by quantitative high throughput microscopy in DIvA cells (Extended Data Fig. 4i). PER2 depletion decreased PLA foci in G1, S and G2 cells (Extended Data Fig. 4j). Collectively, these data suggest that PER2 contributes to TC-DSB targeting at the NE.

## Tethering of TC-DSBs to the NE depends on SUN1 and NUP153

Interestingly, we and others previously reported a function for SUN domain containing-proteins, which are components of the Inner Nuclear Membrane (INM), in DSB mobility in mammals^10,48^. Moreover, SUN protein orthologs in yeast and *Drosophila* (respectively Mps3, and Koi/Spag4) were also found to act as DSB anchoring points in the NE^15–17^. We thus investigated the potential involvement of the two main SUN proteins in mammals, SUN1 and SUN2, in anchoring TC-DSBs to the NE. Interestingly, ChIP against SUN proteins (Extended Data Fig. 5a) revealed that, upon damage, SUN1 displayed enrichment at TC-DSBs compared to NHEJ-DSBs (Extended Data Fig. 5b, top panel). Surprisingly, SUN2 rather showed decreased occupancy post-DSB induction (Extended Data Fig. 5b, bottom panel), suggesting that TC-DSBs are specifically interacting with SUN1 and not SUN2. Of note SUN1 recruitment at TC-DSB was independent of the activity of the three main DNA Damage Response (DDR) kinases, namely ATM, ATR and DNAPK (Extended Data Fig. 5c). To further investigate SUN protein recruitment at DSBs, we performed ChIP-seq against SUN1 and SUN2 before and after DSB induction in DIvA cells. Visual inspection and peak calling on SUN1 and SUN2 ChIP-seq datasets revealed that in undamaged conditions, both proteins are enriched on genes (Extended Data Fig. 5d-e), accumulated at promoters (Extended Data Fig. 5f) and largely overlapped on the genome (Extended Data Fig. 5g). Of interest, post-DSB induction, SUN1 (purple track), was recruited on approximately 1kb around the break, while SUN2 rather displayed eviction (blue track) (see an example Fig. 3f top panel, and average profiles at all DSBs Fig. 3g). Both SUN1 recruitment and SUN2 eviction at DSBs were statistically significant when compared to random, undamaged genomic positions (Extended Data Fig. 5h). Notably, the profile of SUN1 at DSBs strongly resembled to that of PER2 (Extended Data Fig. 5i), and as PER2, SUN1 also showed enrichment at endogenous DSB hotspots (Extended Data Fig. 5j). Moreover, as observed for PER2, SUN1 was specifically recruited at a TC-DSB (Fig. 3f, top panel) but not at a DSB induced in a silent locus (Fig. 3f, bottom panel). On average, SUN1 displayed enhanced accumulation at DSBs with high RNAPII levels compared to those with low RNAPII levels (Extended data Fig. 5k). In agreement, as for PER2, HR-prone DSBs showed higher level of SUN1 as compared to NHEJ-prone DSBs (Extended Data Fig. 5l left panel) while SUN2 depletion was exacerbated at HR-prone DSBs compared to NHEJ-prone DSBs (Extended Data Fig. 5l right panel).

**Figure 5.**
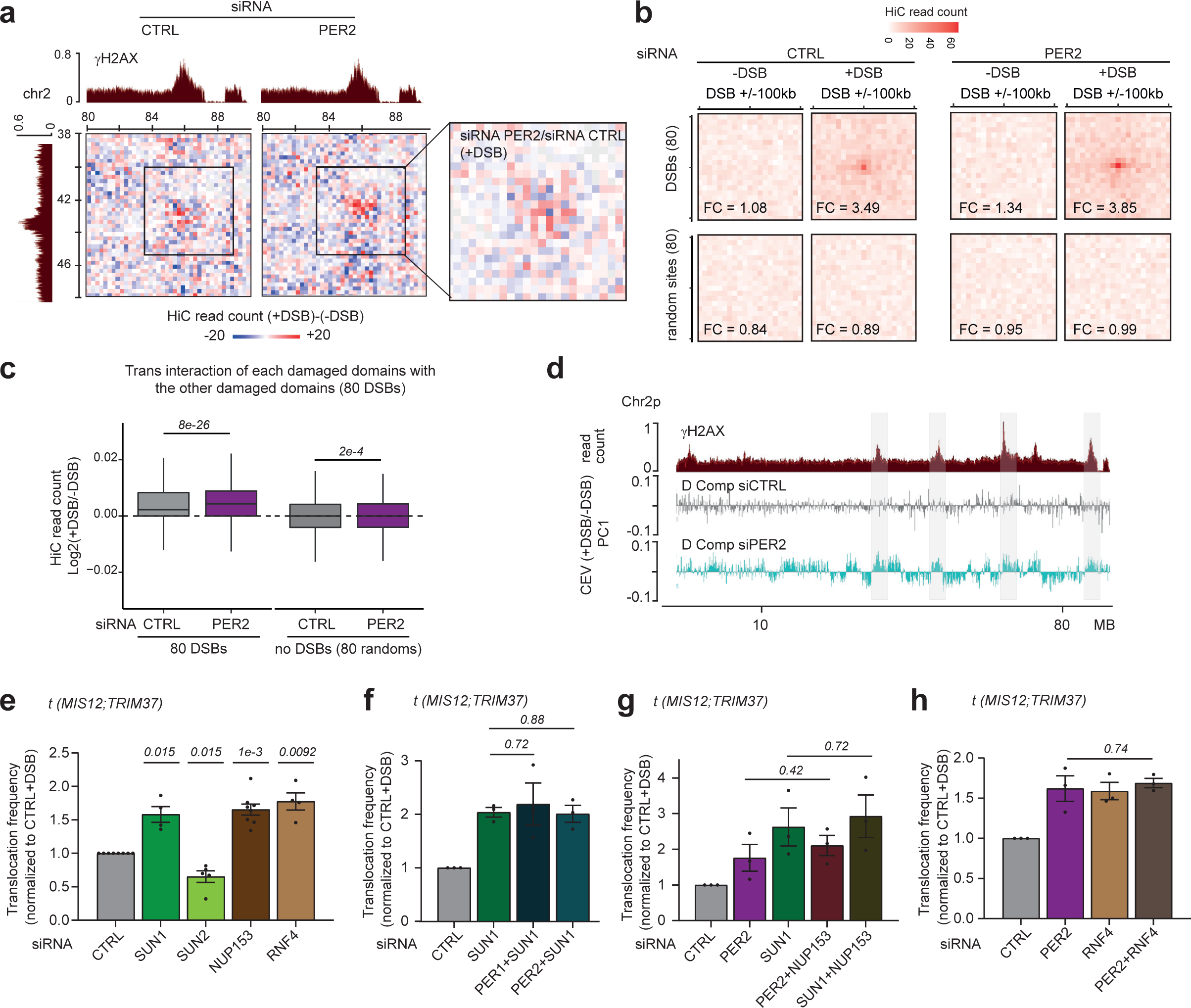
PER proteins, SUN1, NUP153 and RNF4-mediated TC-DSB anchoring to the nuclear envelope counteract DSB clustering and D-compartment formation **a.** Differential Hi-C contact matrix [(+DSB)-(-DSB)] on a region located on chromosome 2 at 25kb resolution in DIvA cells transfected with a control siRNA or a siRNA directed against PER2 as indicated. γH2AX ChIP-seq track (+DSB) are also shown (dark red). **b.** Interchromosomal DSB interaction shown as aggregate peak analysis (APA) plotted on a 200kb window (10kb resolution) before (-DSB) and after DSB (+DSB) induction in control and PER2 siRNA-transfected DIvA cells for the eighty best-induced AsiSI-DSBs (left) or eighty random sites (right, no DSB). FC: Fold change calculated between the central pixel and a square of 3×3 pixels on the bottom left corner of the matrix. APAs show an increased signal at the center (DSB-DSB interaction) after DSBs induction only for DSB sites (FC= 3.49, compared to 0.89 for random sites)). The signal increases after PER2 depletion (FC= 3.85). **c.** Box plot showing the differential Hi-C read counts (as (log2 +DSB/-DSB)) between the 80 best induced DSBs (+/-500kb) (left) or 80 random sites (right) in control (CTRL, grey) and PER2 (purple) siRNA-transfected DIvA cells. Center line: median; box limits: 1st and 3rd quartiles; whiskers: maximum and minimum without outliers. *P*, non-parametric Wilcoxon test. **d.** Genomic tracks of γH2AX ChIP-seq after DSB induction (dark red) and of Chromosomal Eigen vectors (CEV) obtained from PCA analyses performed on differential +DSB/-DSB Hi-C matrices. CEV in siCTRL (grey) and siPER2 (blue). The D-compartment is represented as positive values. siPER2 triggered the appearance of the D-compartment on the chromosome 2 normally devoid of D compartmentalization post-DSB induction in DIvA cells (areas highlighted in light grey). **e-h**. *t(MIS12:TRIM37)* rejoining frequency after DSB induction and repair measured by qPCR in AID-DIvA cells transfected with the indicated siRNAs. Mean and SEM of n biological replicates are shown. *P*, paired t-test (two-sided). **e**. Control (CTRL), SUN1, SUN2, NUP153 and RNF4 siRNAs (n≥4). **f**. CTRL, SUN1 and double PER1+SUN1 and PER2+SUN1 siRNAs (n=3). **g**. CTRL, PER2, SUN1 and double PER2+NUP153 and SUN1+NUP153 siRNAs (n=3). **h**. CTRL, PER2, RNF4 and double PER2+RNF4 siRNAs (n=3).

We then performed 3D-super resolution imaging by RIM to detect SUN1 and SUN2. Both proteins localized to the NE and displayed punctuated patterns (Extended Data Fig. 6a). Co-staining with γH2AX enabled the detection of a significant colocalization (ICQ) with SUN1, but not with SUN2 (Extended Data Fig. 6b). Van Steensel Cross-correlation function (CCF) analysis showed a non-random overlap between γH2AX foci and SUN1 while non-random exclusion was observed between γH2AX foci and SUN2 (see examples Fig. 3h and quantification Fig. 3i). In agreement with a function of SUN1 in tethering TC-DSB to the NE, depletion of SUN1 (Extended Data Fig. 6c) abolished Lamin B1 enrichment at TC-DSBs (Fig. 3j), and also decreased PLA signal between LaminB1 and γH2AX following etoposide treatment (Extended Data Fig. 6d). Altogether, these data suggest that TC-DSBs are physically targeted to the NE through an interaction with SUN1 and not SUN2.

Interestingly, SUN1 has been reported to interact with the Nuclear Pore Complex (NPC), as opposed to SUN2^49,50^. In agreement, in DIvA cells, we found that SUN1 exhibits significant colocalization with NUP153, a protein part of the NPC basket, as compared to SUN2 (Extended Data Fig. 6e). Given that the NPC has also been identified as a docking site for persistent DSBs and collapsed replication forks in yeast^14,18,19,38,51^ and heterochromatic DSBs in drosophila^15^ we thus postulated that the specificity of SUN1 as opposed to SUN2 to anchor DSBs at the NE may be related to its association to the NPC. Indeed, NUP153 accumulates at TC-DSBs but not at a DSB induced in an inactive locus (Extended Data Fig. 6f). 3D-super resolution RIM and CCF analysis further indicated that, as SUN1, NUP153 displays significant colocalization with γH2AX foci (Fig. 3k, examples on the left panel and quantification on the right panel). Of importance, depletion of NUP153 (Extended Data Fig. 6c) triggered a decreased association of TC-DSB with the NE, as measured by PLA between γH2AX and LaminB1 (Extended Data Fig. 6g), further confirming a function of NUP153 in TC-DSB anchoring. Overall, our data suggest that TC-DSBs are tethered to the NE through a SUN1 and NPC-dependent mechanism.

## PER2 and SUN1-dependent DSB anchoring at the NPC promotes HR

Given that PER2, SUN1 and NUP153 are recruited at HR-prone DSBs (Fig. 2h, Extended Data Fig. 5l, Extended Data Fig. 6f) and that previous work showed that targeting DSBs and collapsed replication forks to the NPC promotes HR in yeast and drosophila^14,15,17–19,38,40,46^, we further investigated whether this pathway contributes to HR-repair at TC-DSBs in human cells. One of the first steps of TC-DSB repair is the transcriptional repression of nearby genes (including the damaged gene itself) which is essential for the proper execution of resection and HR repair^1,52^. Given that the PER complex is a transcriptional repressor, we envisaged that it could promote HR repair at TC-DSBs via a role in transcriptional repression. However, RT-qPCR analyses revealed that both PER2 and PER1 are dispensable for transcriptional repression following DSB induction (Fig. 4a). Similarly, SUN1 depletion did not modify the ability of TC-DSB to trigger transcriptional repression (Fig. 4b) showing that the PER2/SUN1-dependent TC-DSB targeting pathway to the NE is not required for the transcriptional inactivation of genes *in cis* to DSBs. We further investigated the effect of SUN1 and PER1 or PER2 depletion on end-resection, by quantification of single-stranded DNA at DSBs^53^ (Extended Data Fig. 6h). siRNA depletion of SUN1 did not alter resection (Extended Data Fig. 6i) while both PER1 and PER2 triggered a mild decrease of resection (Extended Data Fig. 6j), although a lot less pronounced than upon CtIP depletion. Altogether, these data suggest that the contribution of NE anchoring in HR repair takes place downstream of transcriptional repression and resection.

In drosophila, the targeting of HC breaks to the NE favors RAD51 loading^15,46^. We thus further assayed in DIvA cells, whether PER2-dependent TC-DSB targeting at the NE regulates RAD51 recruitment. Indeed, RAD51 loading at TC-DSBs was significantly reduced in PER2-depleted cells (Fig. 4c). Of interest, in yeast, the eviction of the single strand DNA binding protein RPA, required to ensure Rad51 nucleofilament formation, is promoted by its ubiquitination by the Sumo-Targeted Ubiquitin Ligase (STUbL) Slx5/8^19^, which is also required to relocate collapsed forks and DSBs to the NE^15,18,19,38,40,46^. We thus further tested RPA and RAD51 recruitment at TC-DSBs upon depletion of SUN1 and RNF4, the Slx5/8 ortholog in human cells which has also been previously reported to be involved in HR repair^54,55^. SUN1 and RNF4 depletion (Extended Data Fig. 6c, Extended Data Fig. 6k) altered the balance between RAD51 and RPA at TC-DSB (Fig. 4d, Extended Data Fig. 6l).

In agreement with our above findings, depletion of SUN1, NUP153 and RNF4 all impaired cell survival post TC-DSB induction, while SUN2 did not, suggesting deficient repair upon impairment of the TC-DSB NE-anchoring pathway (Fig. 4e-g). SUN1 and NUP153 depletion also impaired cell survival following etoposide in RPE1 cells (Extended Data. Fig. 6m). Importantly co-depletion of both SUN1 and PER1 or PER2 did not exacerbate the impaired survival observed upon depletion of PER1 or PER2 proteins independently (Fig. 4h). Similarly, co-depletion of NUP153 and PER2 or SUN1 did not trigger a significant decrease of cell survival as compared to NUP153 depletion alone (Fig. 4i), nor did co-depletion of RNF4 and PER2 (Fig. 4j), indicating they belong to the same pathway.

**Figure 6.**
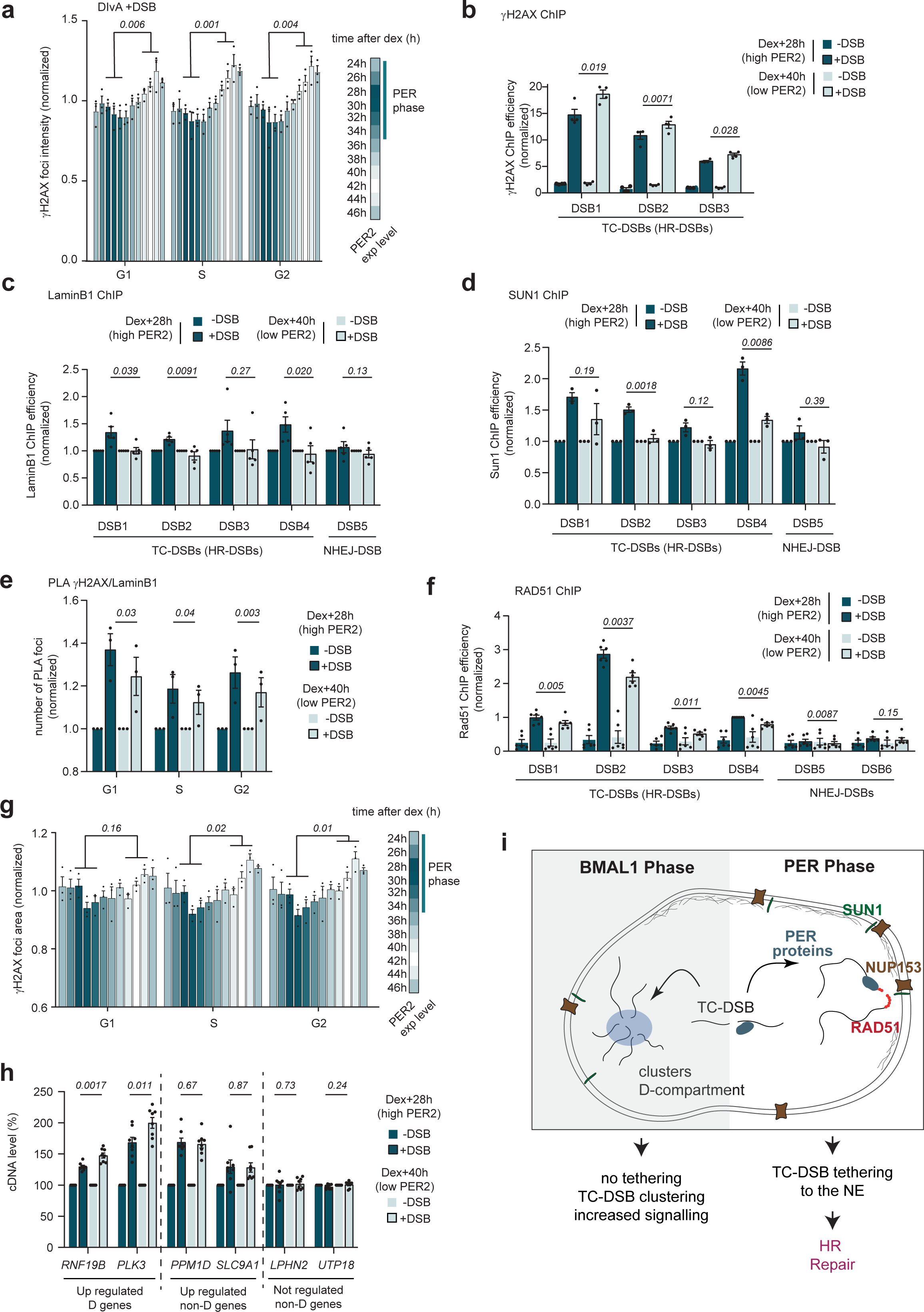
The Circadian rhythm regulates TC-DSB targeting to nuclear envelope and RAD51 loading **a.** γH2AX foci intensity, measured by high content microscopy, in DIvA cells after DSB induction (+DSB 2h), in G1, S and G2 phases at different time points after dexamethasone (Dex) treatment (synchronization of the circadian rhythm). Bars at the different time points are colored according to the PER2 expression level (from highest in dark cyan, to lowest in white). Data were normalized to the average. Mean and SEM (n = 3 biological replicates, with >3000 nuclei acquired per experiment) are shown. *P*, paired t-test (two-sided) was computed using high PER time-points (28h, 30h, 32h) compared to low PER time-points (40h, 42h, 44h). **b.** γH2AX ChIP-qPCR performed in DIvA cells before (-DSB) and after DSB (+DSB) induction at 28h and 40h after dexamethasone treatment (Dex) at three TC-DSBs (DSB1-3). Data were normalized to a control location devoid of DSB. Mean and SEM for n=4 biological replicates. *P,* paired t-test (two-sided). **c.** LaminB1 ChIP-qPCR performed in DIvA cells before (-DSB) and after DSB induction (+DSB) at 28h and 40h after dexamethasone (Dex) treatment at four TC-DSBs (DSB1-4) and DSB5 (DSB in silent locus). Data were normalized to a control location devoid of DSB and expressed relative to the undamaged condition. Mean and SEM for n=5 biological replicates. *P*, paired t-test (two-sided). **d.** SUN1 ChIP-qPCR performed in DIvA cells before (-DSB) and after DSB induction (+DSB) at 28h and 40h after dexamethasone (Dex) treatment at four TC-DSBs (DSB1-4) and DSB5. Data are normalized to a control location devoid of DSB and further expressed related to data obtained in the undamaged condition. Mean and SEM for n=3 biological replicates. *P*, paired t-test (two-sided). **e.** Number of PLA foci (γH2AX-LaminB1) per nucleus in DIvA cells before (-DSB) and after DSB induction (+DSB) at 28h and 40h after dexamethasone (Dex) treatment. Data were normalized to the untreated samples. Mean and SEM (n=3 biological replicates) are shown. *P*, paired t-test (two-sided). **f.** RAD51 ChIP-qPCR performed in DIvA cells before (-DSB) and after DSB (+DSB) induction at 28h and 40h after dexamethasone (Dex) treatment at four TC-DSBs (DSB1-4) and two NHEJ-DSBs (DSB5-6). Data were normalized to a control location devoid of DSB and expressed relative to DSB4-28h. Mean and SEM for n=6 biological replicates. *P*, paired t-test (two-sided). **g.** γH2AX foci mean area, measured by high content microscopy, in DIvA cells after DSB induction (+DSB 2h), in G1, S and G2 phase at different time points after dexamethasone (Dex) treatment. Data were normalized to the average. Mean and SEM (n=3 biological replicates) are shown. *P*, paired t-test (two-sided) was computed using high PER time points (28h, 30h, 32h) compared to low PER time points (40h, 42h, 44h).. **h.** DSB responsive genes expression was measured by RT-qPCR before (−DSB) and after (+DSB) DSB induction in DIvA cells at 28h and 40h after dexamethasone treatment. cDNA level of two D-compartment DSB responsive gene (*RNF19B* and *PLK3*), two non-D compartment DSB responsive genes (*PPM1D* and *SLC9A1*) and two control genes not regulated post DSB induction (*LPHN2*, *UTP18*) are shown. Mean and SEM for n=4 biological replicates. *P,* paired t-test (two-sided). **i.** Model. During the day (PER phase), PERIOD proteins are recruited at DSBs occurring in transcribed loci (TC-DSBs). This triggers the targeting of the TC-DSBs to the NE and their tethering to SUN1 and the Nuclear Pore Complex, further ensuring RAD51 loading and HR repair. During the night (BMAL1 phase), in absence of PER1/2, impaired TC-DSB anchoring to the NE leads to impaired RAD51 loading, and enhanced DSB clustering and D-compartment formation.

Taken together, our data suggest that TC-DSB anchoring to the NE mediated by a PER2, SUN1, NUP153 and RNF4-dependent pathway, fosters RAD51 assembly, thereby ensuring efficient repair and cell survival post DSB induction.

## PER2-mediated DSB anchoring to the NE counteracts DSB clustering and D-compartment formation

In mammalian nuclei, principal component analysis (PCA) of Hi-C experiments revealed that chromatin compartmentalizes to form the so-called “A” and “B” compartments, with the B-compartment corresponding to microscopically visible heterochromatin foci, while the A-compartment correspond to euchromatin^56^. Of interest, both experimental data and computational modeling of chromosome behaviour showed that targeting at the nuclear lamina prevents heterochromatin to coalesce into internal larger nuclear bodies^57^. The current model postulates that upon detachment of chromatin from the NE, phase separation allows multiple HC foci to cluster thus increasing B-compartmentalization. Importantly, we recently reported that upon DSB induction, another chromatin compartment forms, so called the D-compartment (for DSB-induced), through polymer-polymer phase separation^11^. This D-compartment that arises via DSB clustering gathers γH2AX-modified topologically associated domains (TADs) as well as additional undamaged loci, including a subclass of DSB-activated genes^11^.

Of interest, our siRNA screen indicated that the depletion of NONO, SETX and DDX17 increased clustering (Fig. 1b, Extended data Fig. 1f) and we found that PER2 contributes in DSBs targeting to the NE (Fig. 3). Hence, we set out to investigate whether PER2 depletion could also increase DSB clustering and D-compartment formation as a result of a lack of damaged TADs anchoring at the NE. To address this point, we performed Hi-C experiments before and after DSB induction in PER2-proficient and deficient cells. PER2 depletion did not drastically affect chromosome organization in undamaged cells (Extended Data Fig. 7a), nor the overall distribution of A and B compartments (Extended Data Fig. 7b). However, PER2 depletion triggered increased compartmentalization as visualized by increased A-A and B-B interaction, as well as decreased A-B interaction (Extended Data Fig. 7c). This agrees with the previously reported role of the PER complex in chromatin anchoring at the NE which would predict an increased compartmentalization upon NE detachment in PER-depleted cells. Upon DSB induction, we could recapitulate our previous finding showing DSB clustering (Extended Data Fig. 7d, arrows). Importantly, PER2 depletion increased DSB clustering as shown on individual events (Fig. 5a, Extended Data Fig. 7d), average aggregate peak analysis (APA) plots of inter-chromosomal or intra-chromosomal DSB-DSB contacts centered on all 80 best-induced DSBs (Fig. 5b, Extended Data Fig. 7e) and box plots quantifying the interactions between DSBs (Fig. 5c). PER2 depletion also triggered enhanced clustering of endogenous DSB hotspots (Extended Data Fig. 7f) and of etoposide-induced DSBs as measured using quantitative high throughput microscopy (Extended data Fig. 7g).

We further computed Chromosomal Eigen Vectors (CEV) using the first PC of Principal Component Analysis (PCA) on differential Hi-C matrices to identify the D-compartment as previously described^11^. As expected, we could recapitulate D-compartment formation upon damage in DIvA cells transfected with a control siRNA (Extended Data Fig. 7h). PER2 deficiency did not alter D-compartment detection on chromosomes previously found to display D-compartment (Extended Data Fig. 7h). However, we could observe D-compartment formation on additional chromosomes upon PER2 depletion (Fig. 5d). Collectively our data suggest that PER proteins prevent DSB clustering and D-compartment formation by fostering the targeting of TC-DSBs to the nuclear lamina.

D-compartment formation fosters the DNA damage response, but it also comes at the expense of increased translocation rate^12,25,58^. In agreement with the above data involving PER2 in counteracting D-compartment formation, increased translocation frequency was observed upon depletion of the PER complex proteins (Fig. 2c). Similarly, and in line with their involvement in TC-DSB tethering at NE (Extended data Fig. 6d, Fig. 3g, Extended data Fig. 6g), SUN1 and NUP153 depletion increased translocation (Fig. 5e) suggesting enhanced DSB clustering in SUN1 and NUP153 deficient cells. Of note, this was not the case when depleting SUN2, in agreement with the absence of SUN2 recruitment at DSBs. Moreover, in agreement with the previously reported role of Slx5/8 in the relocation of persistent DSBs and arrested forks in yeast^14,19,40,51^, as well as of HC breaks in drosophila^15^, RNF4 depletion also increased translocation frequency (Fig. 5e), suggesting enhanced DSB clustering in absence of the STUbL

RNF4. Importantly co-depletion of both SUN1 and PER1 or PER2 did not further increased translocation compared to SUN1 depletion alone (Fig. 5f). Similarly, co-depletion of NUP153 and PER2 or SUN1 did not trigger a significant increase in translocation frequency as compared to NUP153 depletion alone (Fig. 5g), nor did co-depletion of RNF4 and PER2 (Fig. 5h). Taken together, our data suggest that TC-DSB anchoring to the NE mediated by a PER2, SUN1, NUP153 and RNF4-dependent pathway, counteracts DSB clustering and translocations in human cells.

## The circadian clock regulates the response to TC-DSBs and their anchoring to the NE

Given that the PER complex is a core component of the circadian clock, we set out to investigate whether the circadian clock could also regulate the response to TC-DSBs. As expected, dexamethasone treatment synchronized the circadian clock in both U2OS and DIvA cells, as assessed by the cycling expression of PER2, BMAL1 and CRY2 (Extended Data Fig. 8a-b). We further monitored γH2AX foci formation in different cell cycle phases, at different time points after dexamethasone-mediated circadian clock synchronization by combining γH2AX staining and EdU labeling, using quantitative high throughput microscopy in DIvA cells. In line with the results obtained with the depletion of PER2 by siRNA, our data indicate that γH2AX foci intensity was increased post-DSB induction during the low-PER2 circadian phase (Fig. 6a, white bars; Extended Data Fig. 8c) as compared to the high-PER2 phase (Fig. 6a blue bars; Extended Data Fig. 8c). This was independent of the cell cycle phase (Fig. 6a). Moreover, increased accumulation of γH2AX at TC-DSBs during the low-PER2 phase, was further confirmed by ChIP (Fig. 6b). Of importance the same holds true in S and G2 phases when DSBs were induced using etoposide (Extended Data Fig. 8d). To further reinforce the notion that the low-PER2 phase of the circadian clock impacts TC-DSB repair, we performed circadian clock synchronization and analyzed several outputs that were affected by the depletion of PER2. First, the circadian clock did not alter transcriptional repression in *cis* to DSBs (Extended Data Fig. 8e) as expected from PER2 depletion data (Fig. 4a). Second, as found using PER2 depletion (Fig. 3d), the low-PER2 circadian phase reduced the recruitment of LaminB1 (Fig. 6c) and SUN1 (Fig. 6d) to TC-DSBs when monitored by ChIP-qPCR, suggesting that DSB anchoring at the NE is defective when the PER1/2 proteins are expressed at low levels. Indeed, γH2AX/LaminB1 PLA signal decreased during the low PER2 phase (Fig. 6e). Third, given that our data indicated a role of PER2 in RAD51 loading at TC-DSBs (Fig.4c), we assessed RAD51 recruitment using ChIP. As expected from PER2 depletion data, RAD51 recruitment at TC-DSBs decreased during the low PER phase (Fig. 6f). In agreement, RAD51 foci formation monitored using high throughput microscopy at different time points after dexamethasone treatment revealed an impaired RAD51 foci formation during the low-PER2 phase (Extended Data Fig. 8f). Fourth, quantitative high throughput microscopy further showed an increase in γH2AX foci area during the low-PER2 phase, in agreement with enhanced DSB clustering (Fig. 6g).

We previously reported that, in addition to γH2AX domains, the D-compartment also physically attracts additional loci including some, but not all, DNA damage responsive (DDR) genes, in order to potentiate their activation post-DSB induction^25^. Notably, we found that *RNF19B* and *PLK3*, two upregulated DDR genes found within the D-compartment, were significantly more activated post-DSB during the low-PER2 phase when compared to the high-PER2 phase (Fig. 6h). This agrees with an increased D-compartment formation in absence of PER2 (Fig. 5a-d, Extended Data Fig. 7h). Of importance, this was not the case for other DDR genes which are not targeted to the D-compartment (*PPM1D*, *SLC9A1*)^25^, or for other non-induced genes (*LPHN2*, *UTP18*). Altogether these data indicate that the circadian rhythm regulates the repair and signaling of TC-DSBs by controlling their PER2-dependent anchoring to the NE and, therefore, the formation of the D-compartment, further fine-tuning the response to DSBs.

## Discussion

In this study, we uncovered that the repair of DSBs occurring in transcriptionally active chromatin in human requires their anchoring to the NE, a process that is under the direct control of PERIOD proteins and is therefore sensitive to circadian oscillations (Fig. 6i). Notably, in a recent preprint, the Mekhail lab reports that etoposide induces nuclear envelope tubules (dsbNET) inside the nucleus which establish contacts with DSBs to foster repair^59^. Since persistent DSBs and arrested forks in yeast as well as HC breaks in *Drosophila*, also relocate to the NE and NPC, altogether our findings suggest that targeting repair refractory DSBs to this specific nuclear microenvironment is evolutionary conserved.

Of interest, previous work uncovered that the maintenance of genomic integrity and cancer progression are tightly linked to the circadian clock^60–62^. In general, genes involved in DNA repair pathways are transcriptionally regulated by the circadian clock^61,63–68^. Here we found that in human cells, PER2 is directly targeted at DSBs. In contrast to a very recent report^69^, we observed neither BMAL1 binding at TC-DSBs, although BMAL1 distribution on the genome behaved as expected, nor a function of BMAL1/CLOCK heterodimer in promoting cell survival and counteracting translocation upon TC-DSBs induction in DIvA cells. This difference may arise from analyzing overexpressed^69^ versus endogenous (our study) BMAL1, from the DSB-inducible systems used in both studies (AsiSI versus Zeocin and I-PpoI) and/or from difference in the time points analyzed post-DSB induction.

Upon PER1 or PER2 depletion, as well as during the low-PER phase, γH2AX level was increased while RAD51 assembly was reduced suggesting a decreased repair capacity. Of interest, in the accompanying paper, the Huertas lab reports that CRY1 is also recruited at radiation-induced DSBs, where it directly represses resection and HR. Altogether our studies suggest a two-arm control of HR repair during the 24h circadian cycle, with minimum HR occurring at night in diurnal species. Indeed, large transcriptomic surveys, performed in baboon^63^ and very recently in human^70^, allowed to carefully establish the transcriptional pattern of core clock regulators during the day. PERIOD genes *Per1*, *Per2* and *Cry2* display a maximum expression level around noon, while *Cry1* expression peaks at sunset, in agreement with a PERIOD complex-independent function of CRY1. This allows us to propose the following model (Extended Data Fig. 8g): in the morning, the presence of the PERIOD PER1/PER2 proteins combined with the absence of CRY1, establishes an environment that favors HR repair of TC-DSBs by allowing resection to occur, DSB anchoring to the NE and RAD51 loading. Subsequently, the gradual increase of CRY1 during the afternoon causes minimum resection at nightfall. The absence of PERIOD complex expression early at night establishes a second layer of HR repression during the night. The gradual decrease of CRY1, combined with the re-expression of PER proteins, then allows HR to resume the next morning.

Of interest, sleep was found to regulate the repair of physiological DSBs in neurons^71^, so whether such a circadian regulation of TC-DSB repair also takes place in post-mitotic cells will deserve further investigation.

In summary, our findings have implications for disease etiology and treatment. Indeed, evidence suggests that a dysregulation of the circadian clock contributes to the progression of neurodegenerative disorders^72^ and cancer initiation and progression^60,62^, both types of diseases being tightly coupled to DSB repair mechanisms and the maintenance of genome integrity. Moreover, and importantly, topoisomerase II poison-based chemotherapies are a first-line treatment against a number of cancers and given that these poisons mainly trigger DSBs in active chromatin, our findings that circadian rhythm regulates the repair of TC-DSBs may be an important feature to take into account for chronochemotherapy.

## Data and Code availability

Code is available at https://github.com/LegubeDNAREPAIR/CircadianClock. High throughput data are available at Array Express (E-MTAB-12712) using the link https://www.ebi.ac.uk/biostudies/arrayexpress/studies/E-MTAB-12712?key=2f8af589-d077-466a-a563-725b33761cda

## Acknowledgments

We thank the genomics core facility of EMBL for high-throughput sequencing. We thank Vanessa Dougados for her help during the RIM acquisitions and the LITC TRI-GENOTOUL photonic imaging platform. We thank L. Ligat from the CRCT Cell Imaging Platform (INSERM-UMR1037) for technical assistance in high-content microscopy. We thank T. Clouaire and N. Firmin for technical help. We thank T. Clouaire, and E. Cau for their advice throughout the project and critical reading of the manuscript. Development of 3D RIM on the LITC was supported by ANR-20-CE45-0024. Funding in GL laboratory was provided by grants from the European Research Council (ERC-2014-CoG 647344; and ERC-AdG-101019963), the Agence Nationale pour la Recherche (ANR-18-CE12-0015), the Fondation ARC pour la recherche sur le cancer, the ITMO Cancer, the Ligue Nationale Contre le Cancer, comité de l’Aude and the Fondation Bettencourt-Schueller. C.A was a recipient of a FRM fellowship (FRM FDT201904007941), S.C. was a recipient of a PhD fellowship from Joint Training and Research Programme on Chromatin Dynamics & the DNA Damage Response (H2020 ITN aDDRess, grant N° 812829), J.F is a recipient of a PhD fellowship from the Horizon Europe Marie Skłodowska-Curie Actions Doctoral Networks CohesiNet (Grant Agreement No. 859853). I.L. is supported by a funding from the Fondation ARC pour la recherche sur le cancer. N.P and P.C are INSERM researchers.

## Authors contributions

B.L.B, L.G.S, J.F, R.C, E.G, C.P, C.A, I.L, A.G, A.L.F, A.M. and N.P, performed and analyzed experiments. P.F and P.C optimized the mAID-AsiSIER lentiviral construct. S.C, M.A, and V.R performed bioinformatic analyses of all high-throughput sequencing datasets. T.M performed RIM acquisition and analysis. G.L. and N.P. supervised the work. G.L. wrote the manuscript. All authors commented, edited and approved the manuscript.

## Methods

### Cell culture and treatment

U2OS, DIvA (AsiSI-ER-U20S)^20^, AID-DIvA (AID-AsiSI-ER-U20S)^8^, 53BP1-GFP DIvA^73^, hTERT RPE1 and RPE-DIvA (mAID-AsiSI-ER-RPE1) cells were grown in Dulbecco’s modified Eagle’s medium (DMEM) supplemented with 10%SVF (Invitrogen), antibiotics and either 1µg/mL puromycin (DIvA cells) or 800µg/mL G418 (AID-DIvA cells) or both puromycin and G418 (53BP1-GFP DIvA) or 10µg/ml Blasticidine (RPE-DIvA) at 37°C under a humidified atmosphere with 5%CO_2_. The cell lines were regularly checked for mycoplasma contamination. For AsiSI-dependent DSB induction, DIvA cell lines were treated with 300nM 4-hydroxytamoxifen (OHT) (Sigma, H7904) for 4h (unless otherwise indicated). For DSB induction with etoposide (Sigma, E1383), U2OS cells were treated with 0,5µM etoposide for 4h, dexamethasone-synchronized cells were treated with 1µM etoposide for 2h and hTERT RPE1 cells were treated with 1µM etoposide for 4h. For DSB induction with doxorubicin (Sigma, D1515), U2OS cells were treated with 0,1µM doxorubicin for 4h. For DDR kinase inhibition, DIvA cells were pretreated for 1h and during subsequent DSB induction with either 20µM ATMi (KU-55933 from Sigma, SML1109), 2µM ATRi (ETP-46464 from Sigma, SML1321) or 2μM DNAPKi (NU-7441 from Selleckchem, S2638). To stop inducing AsiSI-DSBs, AID-DIvA cells were washed twice in pre-warmed PBS after OHT treatment and further incubated with 500µM auxin (IAA) (Sigma, I5148) for 2h (unless otherwise indicated). For circadian rhythm synchronization, cells were incubated for 1h with 500nM dexamethasone (Dex) Sigma, D4902), then the medium was replaced (time = 0h). Dex treatment was done every 2h during 24h and the DSB induction was started for all synchronized cells at the same time, 24h after the last Dex treatment, to obtain a full period circadian rhythm (analyses are shown from 24h to 46h post-Dex). To induce DSBs, synchronized cells are treated during 2h or 4h surrounding the synchronization timepoint (*e.g.*, from 23h to 25h after Dex for a 2h treatment corresponding at the synchronized time = 24h).

### RPE-DIvA cell line generation

#### Generation of mAID-AsiSI expression vectors

*pLV-OsTIR1-2Myc-T2A-mAID-HA-ER-NLS-AsiSI-P2A-PuroR*. The lentiviral vector allowing the coexpression of (i) the AsiSI restriction enzyme amino-terminally fused to a mini-auxin inducible degron (mAID)^74^, a human influenza hemagglutinin (HA) tag, a modified estrogen receptor ligand binding domain (ER) and a nuclear localization signal (NLS) of SV40 large T antigen, (ii) the Myc-tagged rice F-box protein TIR1, and (iii) an antibiotic resistance selectable marker, was generated from the previously described pLV3 plasmid^75^ by the following sequential modifications. First, a coding sequence for the T2A self-cleaving peptide was inserted between the Kpn2I and MluI restriction sites using pre-annealed T2A-F and T2A-R oligonucleotides. The mAID cDNA was then PCR-amplified using mAID-F and mAID-R primers and the pAID1.1-N plasmid (BioROIS, Japan) as a template, and inserted between XmaI and EcoRI restriction sites of the previous plasmid. A P2A cassette encoding the self-cleaving peptide (pre-annealed P2A-F and P2A-R oligonucleotides) was further inserted between MluI and EcoRI sites. Next, a cDNA fragment coding for puromycin-resistance was amplified by PCR with primers Puro-F and Puro-R, and added by Hot-Fusion^76^ between EcoRI and NdeI restriction sites. The HA-ER-NLS-AsiSI coding sequence was then PCR-amplified from a previously described pBabe-Puro-HA-ER-NLS-AsiSI vector^20^ with primers AsiSI-F and AsiSI-R and inserted by Hot-Fusion between BamHI and MluI restriction sites. Finally, the OsTIR1-2Myc coding sequence was amplified by PCR (primers TIR1-F and TIR1-R) from a synthetic DNA construct with optimized codons and inserted by Hot-Fusion at the MssI site. *pLV-OsTIR1-2Myc-T2A-mAID-HA-ER-NLS-AsiSI-P2A-BSD*. As hTERT RPE1 are already resistant to puromycin, puromycin-resistance gene was switched to blasticidin-resistance by NEBuilder HiFi DNA Assembly strategy. Amplification of pLV-OsTIR1-2Myc-T2A-mAID-HA-ER-NLS-AsiSI-P2A-BSD was done by PCR using F1_fwd, F1_rev, F2_fwd, F2_rev, F3_fwd, F3_rev primers. Coding sequence for blasticidin resistance was amplified from pCRIS-PITCHv2_BSD_dTAG plasmid (Addgene, #9179) using primers BSD_fwd and BSD_rev. PCR were performed using PrimeSTAR Max DNA Polymerase (Takara, ref R045A). 0.1pmol of each fragment were then assembled for 1h at 50°C using NEBuilder HiFi DNA Assembly master mix from the NEBuilder HiFi DNA Assembly Cloning Kit (NEB, ref E5520S). Reaction products were amplified in provided bacteria and purified using QIAprep Spin Miniprep Kit.

The intermediate and final constructs were checked by sequencing (Eurofins Genomics, Ebersberg, Germany). All restriction and modification enzymes were purchased from Thermofisher Scientific (Illkirch, France). Oligonucleotides were from Eurofins Genomics. Production of lentiviral particles in HEK-293T cells and transduction of cells were performed as previously described^77^.

#### Generation of RPE-DIvA cell line

3.10^5^ cells were infected with lentiviral particules in DMEM, Hepes pH 7.4 10mM and polybrene 8µg/ml. Media was changed 18h after and transduced cells were selected with 12µg/ml blasticidin for 10 days. Clones were isolated and selected according to γH2AX and AsiSI staining by immunofluorescence, and survival after DSB induction by 300nM OHT treatment for 4 hours.

### Multi-output screen methodology

Each of the 130 SMARTpool siRNAs from the TC-DSBR-focused siRNA library (Extended Data Table 1) was transfected in 10^6^ AID-DIvA cells as described below (siRNA and plasmid transfection section), always including a negative (CTRL) and a positive (SETX) control siRNAs. Transfected cells were plated (i) in 96-well Cell Carrier Ultra plates (20.000 cells/well in triplicates per condition) for γH2AX foci analysis (intensity and area) with quantitative high throughput imaging as described below (High-Content microscopy and Immunofluorescence sections); (ii) in 96-well plates (1000 cells/well in quadruplicates per condition for cell survival analysis as described below (WST-1 cell survival assay section); and in 10cm diameter dish for genomic DNA extraction and translocation frequency analysis as described below (DSB-induced rearrangement /Translocation assay section). All data from these different outputs post-DSB induction were expressed normalized to the negative control siRNA (CTRL) and compared to the positive control siRNA (SETX). For the 130 genes of interest, each measured variable (“γH2AX.foci Intensity”, “Clustering”, “Translocation” and “Survival”) was centered and scaled to obtain the z-score displayed in a heatmap. Genes were ordered according hierarchical clustering computed with Pearson correlation distance and ward.D clustering method. Then genes were divided in 2 groups based on the hierarchical clustering and means of z-scores for each group were computed and displayed in independent boxplot for each variable. Mean differences between groups were tested using student t-test.

### siRNA and plasmid transfection

In U2OS and U2OS derived cell lines, siRNA transfections were performed using the 4D-Nucleofector and the SE cell line 4D-Nucleofector X kit**s** (Lonza) according to the manufacturer’s instructions. Briefly, 1-10µL of 100µM annealed siRNA was transfected in 1-20×10^6^ cells with 20-100µL SE solution in Nucleocuvettes using the U2OS program CM-104.

In RPE1 cell lines, 20nM of siRNA were transfected using Lipofectamine RNAiMAX reagent (Invitrogen 13778075) according to the manufacturer’s instructions. Subsequent cell treatments were performed ∼42-48h post-transfection. Most siRNAs used in this study were siGENOME SMARTpools (Dharmacon) which are a mixture of 4 individual siRNAs, see Extended Data Table 1. Individual siRNA sequences (Eurogentec) were used for SUN1, SUN2 or RBBP8/CtIP (Extended Data Table 1). For all experiments, the control siRNA (CTRL) was the siGENOME Non-Targeting Control siRNA Pool #2 (Dharmacon), except for Fig. 4h and Fig. 4i where the control siRNA used with SUN1 and SUN2 siRNAs was Ctrl1 (Extended Data Table 1). Plasmid transfections were performed using the 4D-Nucleofector and the SE cell line 4D-Nucleofector X S kit (Lonza) according to the manufacturer’s instructions. Briefly, 0.5µg DNA was transfected in 10^6^ cells with 20µL SE solution in Nucleocuvette strips using the U2OS program CM-104, and subsequent OHT treatments were performed 48h later. mCherry-LaminB1-10 plasmid was a gift from Michael Davidson (Addgene plasmid #55069; http://n2t.net/addgene:55069; RRID:Addgene_55069).

### Immunofluorescence

Transfected 53BP1-GFP DIvA cells were grown on glass coverslips and fixed with 4% Paraformaldehyde during 15min at room temperature. After, 2 washes with PBS, the permeabilization step was performed by treating cells with 0,5%Triton X-100 in PBS for 10min, then cells were blocked with PBS-BSA3% for 30min. Primary antibody against γH2AX (Extended Data Table 2) was diluted in PBS-BSA3% and incubated with cells overnight at 4°C. After 3 washes for 5min in PBS-BSA3%, cells were incubated with anti-mouse secondary antibody (conjugated to Alexa594 or Alexa488, Invitrogen), diluted 1:1000 in PBS-BSA3%, for 1h at room temperature. After a Hoechst 33342 (Invitrogen) staining (5µg/ml for 5min at room temperature), Citifluor (AF1-25, Clinisciences) was used for coverslip mounting.

### High content microscopy (QIBC)

U2OS, DIvA (AsiSI-ER-U20S)^20^, AID-DIvA (AID-AsiSI-ER-Transfected or Dex-synchronized DIvA and U20S cells were plated in 96-well Cell Carrier Ultra plates (Perkin Elmer). Prior to the end of OHT, etoposide or bleomycin treatments, cells are treated with 10µM EdU (Invitrogen, C10340) at 37 °C for 15min. For γH2AX staining, cells are fixed and permeabilized as described above. For RAD51 staining, buffer II pre-extraction (20mM NaCl, 0.5% NP-40, 20mM HEPES at pH 7.5, 5mM MgCl_2_, 1 mM DTT) was added to cells for 20min on ice followed by fixation in 4% Paraformaldehyde during 15min at room temperature and washed twice with PBS. Then EdU was labeled by Click-iT AlexaFluor 647 reaction (Invitrogen, C10340) according to the manufacturer’s instructions (cells protected from light from now). Cells were washed twice for 5min in PBS-BSA3% and then blocked with PBS-BSA3% for 30min at room temperature. Primary antibodies targeting γH2AX or RAD51 (Extended Data Table 2) were diluted in PBS-BSA3% and incubated with cells overnight at 4°C. After 3 washes of 3min in PBS-BSA3%, cells were incubated with anti-mouse Alexa 488 secondary antibody (Invitrogen) diluted 1:1000 in PBS-BSA3%, for 1h at room temperature. After 3 washes of 3min in PBS, nuclei were labeled with 5µg/ml Hoechst 33342 (Invitrogen) for 30min at room temperature. The cells were then washed twice with PBS and stored at 4°C. γH2AX foci were further analyzed with an Operetta CLS High-Content Imaging System (Perkin Elmer) and Harmony software (version 4.9). For quantitative image analysis, ∼20-80 fields per well were acquired with a 20X (for γH2AX circadian synchronization) or 40X objective lens to visualize >3000 cells per well in triplicate. Subsequent analyses were performed with Columbus software (version 2.8.2). Briefly, the Hoechst-stained nuclei were selected with B method, and the size and roundness of nuclei were used as parameters to eliminate false positive compounds. Then, the Find Spots D method was used (Detection sensitivity =0.2-0.4; Splitting coefficient =1; Background correction =0.5) to determine the number, mean area and intensity of repair foci in each nucleus. For cell cycle analysis, the sum of the Hoechst intensity and the mean of the EdU intensity were plotted in order to select G1, S, and G2 cells.

Determination of the number (y axis) and size of γH2AX foci (mean area on the x axis) in each nucleus has been used to infer DSB clustering (>8000 nuclei analyzed per sample) and the resulting scatter plots are divided into four quadrants on the basis of the medians^10,11^. Percentages of cell populations with low number of γH2AX foci of large size (lower right-side area of the scatter plot), called “cluster positive cells”, or with high number of γH2AX foci of small average size (upper left area of the scatter plot), called “cluster negative cells”, are determined.

### DSB-induced rearrangement/translocation assay

AID-DIvA cells were treated as indicated and DNA was extracted using the DNeasy kit (Qiagen). Illegitimate rejoining frequencies between different AsiSI sites, *t(MIS12;TRIM37)* and *t(LINC00217;LYRM2)*, were analyzed by qPCR (in 3-4 replicates) using primers referenced in Extended Data Table 1^23^. Results were normalized using two control regions (Norm1 and Norm17) both far from any AsiSI sites and γH2AX domains (primers in Extended Data Table 1). Normalized translocation frequencies were calculated using the Bio-Rad CFX Manager 3.1 software.

### WST-1 cell survival assay

Transfected AID-DIvA cells were plated in 96-well plates for each of the 3 conditions: -DSB, +DSB, +DSB+repair. 48h later, cells were treated or not with OHT for 4h and further incubated or not with IAA for 4h. Eight days after DSB induction, the medium is replaced by a mix of 100µl warmed medium + 10µl WST-1 ready-to-use solution per well and the cells were incubated for 2h at 37°C. The tetrazolium salts from Premix WST-1 Cell Proliferation Assay System (TAKARA) are cleaved by mitochondrial dehydrogenase in viable cells to produce formazan dye, which is quantified by measuring its absorbance at 450nm (Multiskan GO Microplate Spectrophotometer from Thermo Fisher Scientific). The percentage of viable cells is normalized to the undamaged cells and compared to the control condition.

### Clonogenic assays

Transfected AID-DIvA cells were seeded, in duplicate, at a clonal density in 10cm diameter dishes and, 48h later, were treated or not with 4OHT for 4h and washed twice in pre-warmed PBS and further incubated with auxin (IAA) for another 4h when indicated. Then, the medium was replaced for all cell dishes. After 8 days at 37°C under a humidified atmosphere with 5%CO_2_, cells were washed in PBS and stained with crystal violet (Sigma, V5265). Images of stained colonies were obtained using the White Sample Tray for the ChemiDoc Touch Imaging System (Bio-Rad) and colonies were counted using the Spot Detector plugin of Icy software, Version 2.0.2.0 (icy.exe program). Only colonies containing more than 50 cells were scored. The same protocol was applied to transfected U2OS and hTERT RPE1 cells treated with etoposide or doxorubicin.

### Western blots

DIvA cells were incubated in RIPA buffer (50mM Tris pH8.0, 150mM NaCl, 0.5% sodium deoxycholate, 1% NP-40, 0.1% SDS) on ice for 20 min. Samples were further incubated with 250 units of Benzonase (Sigma, E1014) per million of cells for 10 min at room temperature, then centrifuged at 13,000 rpm for 10 min at 4°C. The supernatants, containing soluble protein extracts, were then mixed with SDS loading buffer and reducing agent, and incubated 10 min at 70°C. Electrophoreses were performed using 4–12% NuPAGE Bis-Tris gels (Invitrogen) and semi-dry western blotting on nitrocellulose membranes (Bio-Rad) was done with the Trans-Blot Turbo System (Bio-Rad) according to the manufacturer’s instructions. Membranes were incubated in TBS containing 0.1% Tween 20 (Sigma, P1379) and 4% BSA during 1h at room temperature for blocking, followed by overnight incubation at 4 °C using primary antibody diluted in TBS-Tween-4% BSA. The corresponding mouse or rabbit horseradish peroxidase-coupled secondary antibodies (Sigma, A2554 and A0545) were used at 1:10,000 to reveal the proteins, using a luminol-based enhanced chemiluminescence HRP substrate (Super Signal West Dura Extended Duration Substrate, ThermoScientific). Picture acquisition of the membranes was done using the ChemiDoc Touch Imaging System (Bio-Rad) and pictures were analyzed using Image Lab software (Bio-Rad). All primary antibodies used in this study are detailed in Extended Data Table 2. Note: although different conditions and several human PER1 antibodies were tested, none of them resulted in a ∼140kDa signal corresponding to the PER1 protein on the western-blots.

### RT-qPCR

Total RNA was extracted from U2OS and DIvA cell lines before and after DSB induction using homogenization with QIAshredder and the RNeasy kit (Qiagen) following the manufacturer’s instructions. RNA was then reverse transcribed to cDNA using the AMV reverse transcriptase (Promega, M510F). qPCR experiments were performed to assess the levels of cDNA using primers referenced in Extended Data Table 1. cDNA levels were then normalized with GAPDH (Gene ID: 2597), TBP (Gene ID: 6908) and 18S (RNA18SN5, Gene ID: 100008588) cDNA levels using the CFX Manager 3.1 software (Bio-Rad).

### Resection assay

Measure of resection was performed as described in^53^ with the following modifications. Genomic DNA was extracted from transfected DIvA cells just after DSB induction using the DNeasy kit (Qiagen) without any vortexing step. 1 µg of DNA was incubated 20min at 37°C with 15 units of RNAseH1 (New England Biolabs) and first digested for 2h at 37°C with 35 units of AsiSI restriction enzyme (New England Biolabs). Then, 250ng of DNA were further digested overnight at 37°C using 10 units of BanI restriction enzyme, that cuts at 200bp from the DSB1, or of BanII restriction enzyme, that cuts at 1122bp from the DSB7 (Extended Data Table 1). Both enzymes were heat inactivated for 20 min at 80°C. Digested and undigested samples were analyzed by qPCR (10ng/well) using primers in Extended Data Table 1. ssDNA% was calculated with the following equation: Percent of ssDNA= 1/((PUISSANCE(2;(Ct Digested − Ct Non Digested) − 1) + 0.5))*100.

### Proximity Ligation Assay (PLA)

PLA experiments were performed using Duolink® In Situ Detection assay from Sigma-Aldrich according to the manufacturer’s instructions. Primary antibodies used for PLA are detailed in Extended Data Table 2.

#### High throughput γH2AX-LaminB1 PLA

siRNA-transfected or Dexamethasone-synchronized cells were plated in 96-well Cell Carrier Ultra plates (Perkin Elmer). At the end of DSB inducing treatment, cells are treated with 10µM EdU (Invitrogen, C10340) for 15min. Cells are fixed with 4% paraformaldehyde during 15min at room temperature. Permeabilization step was performed by treating cells with 0,5% Triton X-100 in PBS for 5min. Then EdU was labeled by Click-iT Alexa488 reaction (Invitrogen, C10340) according to the manufacturer’s instructions. Cells were blocked with 1 drop of blocking solution provided with the Duolink® PLA Probes (DUO92002 and DUO92004, Sigma-Aldrich) for 1h at 37°C in a humid chamber. Primary antibodies targeting γH2AX and LaminB1 were diluted in the antibody dilution buffer provided with the Duolink® PLA Probes and incubated with cells overnight at 4°C. After two 5min washes in Duolink® In Situ Wash Buffer A (DUO82049, sigma-Aldrich), cells were incubated with anti-mouse plus and anti-rabbit minus probes for 1h at 37°C, under slow rotation. After two washes in wash buffer A for 5min, cells were incubated for 30min at 37°C in the ligation reaction mix from the Duolink® In Situ Detection Reagents Red (DUO92008, Sigma-Aldrich). The cells were washed again twice in wash buffer A for 5min and incubated for 1h40 at 37°C under slow rotation for the amplification step. Finally, after two 10min washes in wash buffer B (DUO82049, sigma-Aldrich), nuclei were stained in the staining solution from the Duolink® In Situ Microplate Nuclear Stain (DUO82064, Sigma-Aldrich) for 30min at 37°C and cells were kept in the Anti-Fade solution (DUO82064, Sigma-Aldrich) until image acquisition. For quantitative image analysis, images were acquired with a 40X objective lens to visualize >3000 cells per well in duplicate. Subsequent analyses were performed with Columbus software (version 2.8.2) to determine the numbers of PLA foci in each nucleus. G1, S and G2 nuclei were selected based on EdU and DAPI staining distribution in all cells.

#### 53BP1-LaminB1 and γH2AX-LaminB1 PLA

siRNA-transfected cells were seeded on 13mm coverslips and treated for 4h with 4OHT before fixation in 4%PFA. Slides were then processed as before, except that there were no EdU staining, and that at the end of the PLA procedure, after the two 10min washes in wash buffer B, cells were then washed for 1min in 0.01% wash buffer B and mounted on slides with Duolink® In Situ Mounting Medium with DAPI (DUO82040, Sigma-Aldrich). Image acquisition was done on Leica DM6000 at 40X objective lens to visualize >175 cells per experiment in triplicate. Subsequent analyses were performed with Cell Profiler software (version 4.2.4) to determine the numbers of PLA foci in each nucleus.

### 3D super resolution using Random Illumination Microscopy (RIM)

#### 3D RIM two cameras setup

The 3D RIM home-made setup is coupled to an inverted microscope (TEi Nikon) equipped with 100x magnification, a 1.49 N.A. objective (CFI SR APO 100XH ON 1.49 NIKON) and two SCMOS cameras (ORCA-Fusion, Hamamatsu) mounted in an industrial apochromatic alignment module (abbelight SA). Fast diode lasers (Oxxius) with wavelengths centered at 488nm (LBX-488-200-CSB) and 561nm (LMX-561L-200-COL) are used for all experiments. The bandpass emission filters in front of the two respective cameras are FF01-514/30-25 for camera 1 and FF01-609/54-25 for camera 2. The binary phase modulator (QXGA fourth dimension) conjugated to the image plane combined with polarization elements are used to generate dynamic speckle on the object plane as described in^47^. The synchronization of the hardware (Z-platform, cameras, microscope, laser and SLM) is performed by an improved version of the commercial software INSCOPER.

#### Fixed 3D RIM acquisition and reconstruction

Two-color 3D RIM imaging was performed with the 3D RIM system. The acquisition of 40 planes was done sequentially with a 120nm step in the image plane. For each image plane, 400 speckles were used to increase the desired super-resolved 3D resolution (95nm,95nm,200nm). The image reconstructions were performed with the software (ALgoRIMhttps://github.com/teamRIM/tutoRIM). The Wiener filter used is 0.01, the deconvolution parameter is 0.02 and the regularization parameter is 0.02 for the first channel images. The Wiener filter used is 0.02, the deconvolution parameter is 0.03 and the regularization parameter is 0.02 for the second channel images. The misalignment between the two cameras and the residual chromatic aberrations are corrected using the SVI (Scientific Volume Imaging) software.

#### 3D+t two colors RIM super-resolution acquisition and reconstruction

3D+t RIM movies were performed on AID-DIvA cells expressing 53BP1-GFP and mcherry-LaminB1. 3D movies of 47min with 40sec intervals were made (60 times). The whole cell acquisition time was 6 seconds with 48 random patterns optimized for live cells for each plane. Image reconstructions were performed with the software (ALgoRIMhttps://github.com/teamRIM/tutoRIM). The Wiener filter used is 0.01, the deconvolution parameter is 0.151 and the regularization parameter is 0.08 for the first channel images. The Wiener filter used is 0.1, the deconvolution parameter is 0.25 and the regularization parameter is 0.08 for the second channel images. The misalignment between the two cameras and the residual chromatic aberrations are corrected using the SVI (Scientific Volume Imaging) software.

#### 3D film editing

Bleaching correction is performed after RIM reconstruction with the open-source FIJI 546 software (https://imagej.net/software/fiji/) based on an exponential FIT from the background. The FIJI 3D drift correction plugin is performed for 3D registration (https://github.com/fiji/Correct_3D_Drift). Rendering of 3D+t movies was performed with the VTK library implemented in ICY software (https://icy.bioimageanalysis.org/) from 3D CROP on 550 areas of interest. Chromatism correction was performed by Scientific Volume Imaging (SVI) software.

#### Colocalization analyses

The colocalization analyzer from the Scientific Volume Imaging (SVI) software is used on fixed 3D RIM images to quantify the interactions of γH2AX nanofoci with those of SUN1 and SUN2.

#### Quantitative immunocolocalization (ICQ)

The intensity correlation quotient was defined in^78^ by the following equation for each voxel “i”:

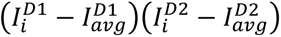

For a random or mixed interaction, this number will tend towards 0, and for a dependent correlation, it will tend towards +0.5. This parameter does not directly use the intensity of each pair of voxels and has the advantage of eliminating the bias towards particularly high or too low intensities.

#### Cross-correlation function CCF

The Van Steensel cross-correlation function (VSCF) CCF is used to quantify interactions and has been described in^79^. It is obtained by calculating the Pearson coefficient after shifting the second camera image over a distance of Δx voxels. Thresholding was carefully tailored for each image to reject the 10^th^ percentile of the lowest intensity value for both channels. The CCF was measured with the x-shift set to 940nm without rotation and the resulting three graphs were averaged and plotted with a 95 percentil. The Pearson coefficient is the classic equation below for each voxel “i”:

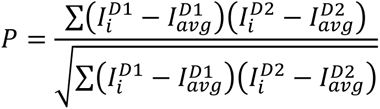

with 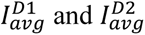 the averages of camera 1 and camera 2 of the microscope.

For better visualization of the shape of the CCF function for each condition, we normalized the CCF function between 0 and 1 with the following normalization calculation: 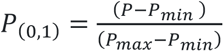, for Figure 3i and 3k or 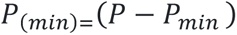 for Extended Data Figure 6e.

Negative values for Δx indicate a shift in nm of the red image to the left, positive values indicate a shift to the right. Non-random overlap results in a peak at Δx=0 and non-random exclusion results in a dip at Δx=0. Uncorrelated distributions will not show any clear peak in the CCF.

### ChIP-qPCR, ChIP-seq and data analysis

#### ChIP-qPCR

ChIP experiments were performed in DIvA cells according to the protocol described in^8,20^ with 200μg of chromatin per immunoprecipitation. The quantity of primary antibodies used is detailed in Extended Data Table 2. qPCR experiments were performed on both IP and input samples to assess the percent of input DNA immunoprecipitated, using AsiSI-induced DSBs and primers referenced in Extended Data Table 1. When indicated in the figure legends, data are normalized to a control location devoid of DSB (Ctrl genomic locus) and could be further expressed related to data obtained at DSB4 or in undamaged condition.

#### Library preparation and sequencing

Multiple ChIP experiment samples were pooled and sonicated for 15 cycles (30 sec on, 30 sec off, high setting) with a Bioruptor (Diagenode), then concentrated with a vacuum (Eppendorf). 10ng of purified DNA (average size 250–300bp) was used to prepare sequencing libraries with the Next Ultra II Library Prep Kit for Illumina (New England Biolabs) using the application note for “Low input ChIP-seq”, and subjected to 75bp single-end sequencing using Illumina NextSeq500 at the EMBL Genomics core facility (Heidelberg, Germany).

#### ChIP-seq data processing

ChIP-seq data processing was performed as described^22^. In brief, raw sequencing reads were aligned using bwa (https://bio-bwa.sourceforge.net/) to the human reference genome (hg19) and then, were sorted, deduplicated, and indexed using samtools (http://www.htslib.org/). Bigwig coverage tracks were subsequently extracted from the processed bam files using bamCoverage from deepTools (HYPERLINK https://deeptools.readthedocs.io/en/develop/), and were normalized by total read count to be used for downstream analyses.

#### DSB Average profiles and boxplots

Differential coverage tracks representing the log2 ratio between damaged and undamaged ChIP-seq samples were created using the bamCompare function of deepTools on default settings. These bigwigs were then used as input for the computeMatrix function of deepTools to calculate the coverage at DSBs using 200 bins covering either 10kb or 1kb as indicated on Figures. These matrices were then processed using a custom script in R with ggplot2.

#### Peak calling

Peak calling for ChIP-seq datasets (BMAL1, SUN1 and SUN2) was performed using *MACS2* using -q 0.1 for BMAL1 resulting in 1776 peaks, and -q 0.005 for SUN1 and SUN2 resulting in 27326 and 14571 peaks respectively. Endogenous DSBs were identified as described in ^11^. In brief, peaks were called from untreated pATM-ChIP-seq data^80^ using macs2, identifying 1,206 regions. A random set of regions was then generated from the gkmsvm R package.

#### Gene Ontology Analysis

Genes for BMAL1 GO analysis were selected if the identified BMAL1 peak was within a genes body or within its promoter region. The subsequent gene list was used to perform gene ontology analysis using the *enrichGO* function from *clusterprofiler* in R.

#### Peak annotation

The genomic locations of SUN1 and SUN2 peaks were identified by using the *ChIPseeker* package in R, specifically the *annotatePeak* function to annotate the regions and *plotAnnoPie* for subsequent visualization. To calculate the coverage of SUN1 at SUN2 regions, the *computeMatrix* function of deepTools was used using 200 bins over a 10kb window. The matrix was then processed using a custom script in R with ggplot2.

### Hi-C and data analyses

#### Hi-C libraries

Hi-C experiments were performed in DIvA cells following transfection with CTRL or PER2 siRNAs and with or without DSB induction as described in^80^. Briefly, 10^6^ cells were used per condition. Hi-C libraries were generated using the Arima-HiC+ for HiC (Arima Genomics) by following the manufacturer’s instructions. DNA was sheared using the Covaris S220 to obtain an average fragment size of 350-400pb. Sequencing libraries were prepared on beads using the Next Ultra II DNA Library Prep Kit for Illumina and Next Multiplex Oligos for Illumina (New England Biolabs) following instructions from the Arima-HiC+ for HiC (Arima Genomics). Hi-C data were processed as described in^25,80^see below.

#### Hi-C heatmaps

Hi-C reads were mapped to hg19 and processed with Juicer v2.0 using default settings (https://github.com/aidenlab/juicer). Hi-C count matrices were generated using Juicer at multiple resolutions: 100kb, 50kb, 25kb, 10kb, 5kb and 1kb. Hi-C heatmap screenshots were obtained using Juicebox (https://github.com/aidenlab/Juicebox/wiki/Download).

#### Aggregate Peak Analysis (APA)

APA plots of inter- or intra-chromosomal interactions of DSBs and control regions were created using the APA program of Juicer tools (https://github.com/aidenlab/juicer/wiki/APA) with a 10kb resolution −/+ 100kb.

#### Trans contact quantification

To determine inter-chromosomal interaction changes (in *trans*), we built the whole-genome Hi-C matrix for each experiment by merging all chromosome-chromosome interaction matrices using Juicer and R to obtain a genome matrix with 33kx33k bin interactions for 100kb resolution. Interactions between bins inside damaged TADs (240X240 for the 80 best-induced AsiSI DSBs) were extracted and counted for each condition, then log2 ratio was calculated on normalized count (cpm), and plotted as boxplots or heatmaps (https://github.com/LegubeDNAREPAIR/ATMcompD/blob/main/scriptR/Heatmap_D_trans. R).

#### A/B compartment

To identify A and B main chromosomal compartments, the extraction of the first Eigen vector of the correlation matrix (PC1) was done on the Observed/Expected matrix at 500kb resolution using juicer eigenvector command. The resulting values were then correlated with ATAC-seq signal in order to attributes positive and negative values to the A and B compartment, respectively, on each chromosome. The Observed/Expected bins were then arranged based on the PC1 values and aggregated into 21 percentiles, to visualize A-B interactions on the siPER2 and siCTRL experiments (saddle plots), and differences were computed between PER2 and control conditions for each percentile.

#### D-compartment

For D-compartment identification, we retrieved the first component (PC1) of a PCA made on the differential observed over expected Hi-C matrix 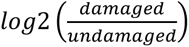at 100kb resolution. Each matrix was extracted from the .hic files using Juicer and the ratio was computed bin per bin. Pearson Correlation matrices were then computed for each chromosome, and PCA was applied to each matrix. The first component of each PCA was then extracted and correlated with the position of DSBs. A PC1 showing a positive correlation with DSBs was then called D-compartment, and PC1 showing negative correlation with DSBs were multiplied by −1. We were able to extract the D-compartment on several chromosomes for +DSB/-DSB^25^. D-compartment (first component of the PCA) was converted into a coverage file using rtracklayer R package. For analysis of DSB responsive gene expression by RT-qPCR, we used the previously determined D compartment genes as described in^25^.

### Contact for Reagent and Resource Sharing

Further requests for reagents and resources should be asked to the lab PI, Gaëlle Legube. DIvA cell lines are subjected to an MTA with the CNRS.

## References

1. Marnef, A., Cohen, S., and Legube, G. (2017). Transcription-Coupled DNA Double-Strand Break Repair: Active Genes Need Special Care. Journal of Molecular Biology 429, 1277–1288. 10.1016/j.jmb.2017.03.024.

2. Canela, A., Maman, Y., Huang, S.N., Wutz, G., Tang, W., Zagnoli-Vieira, G., Callen, E., Wong, N., Day, A., Peters, J.-M., et al. (2019). Topoisomerase II-Induced Chromosome Breakage and Translocation Is Determined by Chromosome Architecture and Transcriptional Activity. Molecular Cell 75, 252–266.e8. 10.1016/j.molcel.2019.04.030.

3. Ballarino, R., Bouwman, B.A.M., Agostini, F., Harbers, L., Diekmann, C., Wernersson, E., Bienko, M., and Crosetto, N. (2022). An atlas of endogenous DNA double-strand breaks arising during human neural cell fate determination. Sci Data 9, 400. 10.1038/s41597-022-01508-x.

4. Gothe, H.J., Bouwman, B.A.M., Gusmao, E.G., Piccinno, R., Petrosino, G., Sayols, S., Drechsel, O., Minneker, V., Josipovic, N., Mizi, A., et al. (2019). Spatial Chromosome Folding and Active Transcription Drive DNA Fragility and Formation of Oncogenic MLL Translocations. Mol Cell 75, 267–283.e12. 10.1016/j.molcel.2019.05.015.

5. Madabhushi, R., Gao, F., Pfenning, A.R., Pan, L., Yamakawa, S., Seo, J., Rueda, R., Phan, T.X., Yamakawa, H., Pao, P.-C., et al. (2015). Activity-Induced DNA Breaks Govern the Expression of Neuronal Early-Response Genes. Cell 161, 1592–1605. 10.1016/j.cell.2015.05.032.

6. Pankotai, T., Bonhomme, C., Chen, D., and Soutoglou, E. (2012). DNAPKcs-dependent arrest of RNA polymerase II transcription in the presence of DNA breaks. Nat Struct Mol Biol 19, 276–282. 10.1038/nsmb.2224.

7. Shanbhag, N.M., Rafalska-Metcalf, I.U., Balane-Bolivar, C., Janicki, S.M., and Greenberg, R.A. (2010). ATM-dependent chromatin changes silence transcription in cis to DNA double-strand breaks. Cell 141, 970–981. 10.1016/j.cell.2010.04.038.

8. Aymard, F., Bugler, B., Schmidt, C.K., Guillou, E., Caron, P., Briois, S., Iacovoni, J.S., Daburon, V., Miller, K.M., Jackson, S.P., et al. (2014). Transcriptionally active chromatin recruits homologous recombination at DNA double-strand breaks. Nat Struct Mol Biol 21, 366–374. 10.1038/nsmb.2796.

9. Ouyang, J., Yadav, T., Zhang, J.-M., Yang, H., Rheinbay, E., Guo, H., Haber, D.A., Lan, L., and Zou, L. (2021). RNA transcripts stimulate homologous recombination by forming DR-loops. Nature 594, 283–288. 10.1038/s41586-021-03538-8.

10. Aymard, F., Aguirrebengoa, M., Guillou, E., Javierre, B.M., Bugler, B., Arnould, C., Rocher, V., Iacovoni, J.S., Biernacka, A., Skrzypczak, M., et al. (2017). Genome-wide mapping of long-range contacts unveils clustering of DNA double-strand breaks at damaged active genes. Nat Struct Mol Biol 24, 353–361. 10.1038/nsmb.3387.

11. Arnould, C., Rocher, V., Saur, F., Bader, A.S., Muzzopappa, F., Collins, S., Lesage, E., Le Bozec, B., Puget, N., Clouaire, T., et al. (2023). Chromatin compartmentalization regulates the response to DNA damage. Nature 623, 183–192. 10.1038/s41586-023-06635-y.

12. Zagelbaum, J., Schooley, A., Zhao, J., Schrank, B.R., Callen, E., Zha, S., Gottesman, M.E., Nussenzweig, A., Rabadan, R., Dekker, J., et al. (2022). Multiscale reorganization of the genome following DNA damage facilitates chromosome translocations via nuclear actin polymerization. Nat Struct Mol Biol. 10.1038/s41594-022-00893-6.

13. Marnef, A., Finoux, A.-L., Arnould, C., Guillou, E., Daburon, V., Rocher, V., Mangeat, T., Mangeot, P.E., Ricci, E.P., and Legube, G. (2019). A cohesin/HUSH-and LINC-dependent pathway controls ribosomal DNA double-strand break repair. Genes Dev. 33, 1175–1190. 10.1101/gad.324012.119.

14. Nagai, S., Dubrana, K., Tsai-Pflugfelder, M., Davidson, M.B., Roberts, T.M., Brown, G.W., Varela, E., Hediger, F., Gasser, S.M., and Krogan, N.J. (2008). Functional Targeting of DNA Damage to a Nuclear Pore-Associated SUMO-Dependent Ubiquitin Ligase. Science 322, 597–602. 10.1126/science.1162790.

15. Ryu, T., Spatola, B., Delabaere, L., Bowlin, K., Hopp, H., Kunitake, R., Karpen, G.H., and Chiolo, I. (2015). Heterochromatic breaks move to the nuclear periphery to continue recombinational repair. Nat Cell Biol 17, 1401–1411. 10.1038/ncb3258.

16. Oza, P., Jaspersen, S.L., Miele, A., Dekker, J., and Peterson, C.L. (2009). Mechanisms that regulate localization of a DNA double-strand break to the nuclear periphery. Genes Dev. 23, 912–927. 10.1101/gad.1782209.

17. Horigome, C., Oma, Y., Konishi, T., Schmid, R., Marcomini, I., Hauer, M.H., Dion, V., Harata, M., and Gasser, S.M. (2014). SWR1 and INO80 Chromatin Remodelers Contribute to DNA Double-Strand Break Perinuclear Anchorage Site Choice. Molecular Cell 55, 626–639. 10.1016/j.molcel.2014.06.027.

18. Kramarz, K., Schirmeisen, K., Boucherit, V., Ait Saada, A., Lovo, C., Palancade, B., Freudenreich, C., and Lambert, S.A.E. (2020). The nuclear pore primes recombination-dependent DNA synthesis at arrested forks by promoting SUMO removal. Nat Commun 11, 5643. 10.1038/s41467-020-19516-z.

19. Whalen, J.M., Dhingra, N., Wei, L., Zhao, X., and Freudenreich, C.H. (2020). Relocation of Collapsed Forks to the Nuclear Pore Complex Depends on Sumoylation of DNA Repair Proteins and Permits Rad51 Association. Cell Rep 31, 107635. 10.1016/j.celrep.2020.107635.

20. Iacovoni, J.S., Caron, P., Lassadi, I., Nicolas, E., Massip, L., Trouche, D., and Legube, G. (2010). High-resolution profiling of γH2AX around DNA double strand breaks in the mammalian genome. EMBO J 29, 1446–1457. 10.1038/emboj.2010.38.

21. Crosetto, N., Mitra, A., Silva, M.J., Bienko, M., Dojer, N., Wang, Q., Karaca, E., Chiarle, R., Skrzypczak, M., Ginalski, K., et al. (2013). Nucleotide-resolution DNA double-strand break mapping by next-generation sequencing. Nat Methods 10, 361–365. 10.1038/nmeth.2408.

22. Clouaire, T., Rocher, V., Lashgari, A., Arnould, C., Aguirrebengoa, M., Biernacka, A., Skrzypczak, M., Aymard, F., Fongang, B., Dojer, N., et al. (2018). Comprehensive Mapping of Histone Modifications at DNA Double-Strand Breaks Deciphers Repair Pathway Chromatin Signatures. Molecular Cell 72, 250–262.e6. 10.1016/j.molcel.2018.08.020.

23. Cohen, S., Puget, N., Lin, Y.-L., Clouaire, T., Aguirrebengoa, M., Rocher, V., Pasero, P., Canitrot, Y., and Legube, G. (2018). Senataxin resolves RNA:DNA hybrids forming at DNA double-strand breaks to prevent translocations. Nat Commun 9, 533. 10.1038/s41467-018-02894-w.

24. Cohen, S., Guenolé, A., Lazar, I., Marnef, A., Clouaire, T., Vernekar, D.V., Puget, N., Rocher, V., Arnould, C., Aguirrebengoa, M., et al. (2022). A POLD3/BLM dependent pathway handles DSBs in transcribed chromatin upon excessive RNA:DNA hybrid accumulation. Nat Commun 13, 2012. 10.1038/s41467-022-29629-2.

25. Arnould, C., Rocher, V., Bader, A.S., Lesage, E., Puget, N., Clouaire, T., Mourad, R., Noordermeer, D., Bushell, M., and Legube, G. (2021). ATM-dependent formation of a novel chromatin compartment regulates the Response to DNA Double Strand Breaks and the biogenesis of translocations (Molecular Biology) 10.1101/2021.11.07.467654.

26. Brown, S.A., Ripperger, J., Kadener, S., Fleury-Olela, F., Vilbois, F., Rosbash, M., and Schibler, U. (2005). PERIOD1-Associated Proteins Modulate the Negative Limb of the Mammalian Circadian Oscillator. Science 308, 693–696. 10.1126/science.1107373.

27. Duong, H.A., Robles, M.S., Knutti, D., and Weitz, C.J. (2011). A Molecular Mechanism for Circadian Clock Negative Feedback. Science 332, 1436–1439. 10.1126/science.1196766.

28. Padmanabhan, K., Robles, M.S., Westerling, T., and Weitz, C.J. (2012). Feedback Regulation of Transcriptional Termination by the Mammalian Circadian Clock PERIOD Complex. Science 337, 599–602. 10.1126/science.1221592.

29. Kim, J.Y., Kwak, P.B., and Weitz, C.J. (2014). Specificity in Circadian Clock Feedback from Targeted Reconstitution of the NuRD Corepressor. Molecular Cell 56, 738–748. 10.1016/j.molcel.2014.10.017.

30. Emerson, J.M., Bartholomai, B.M., Ringelberg, C.S., Baker, S.E., Loros, J.J., and Dunlap, J.C. (2015). *period* −1 encodes an ATP-dependent RNA helicase that influences nutritional compensation of the *Neurospora* circadian clock. Proc. Natl. Acad. Sci. U.S.A. 112, 15707–15712. 10.1073/pnas.1521918112.

31. Ogilvie, V.C. (2003). The highly related DEAD box RNA helicases p68 and p72 exist as heterodimers in cells. Nucleic Acids Research 31, 1470–1480. 10.1093/nar/gkg236.

32. Takahashi, J.S. (2017). Transcriptional architecture of the mammalian circadian clock. Nat Rev Genet 18, 164–179. 10.1038/nrg.2016.150.

33. Sessa, G., Gómez-González, B., Silva, S., Pérez-Calero, C., Beaurepere, R., Barroso, S., Martineau, S., Martin, C., Ehlén, Å., Martínez, J.S., et al. (2021). BRCA2 promotes DNA-RNA hybrid resolution by DDX5 helicase at DNA breaks to facilitate their repair‡. EMBO J 40, e106018. 10.15252/embj.2020106018.

34. Qu, M., Qu, H., Jia, Z., and Kay, S.A. (2021). HNF4A defines tissue-specific circadian rhythms by beaconing BMAL1::CLOCK chromatin binding and shaping the rhythmic chromatin landscape. Nat Commun 12, 6350. 10.1038/s41467-021-26567-3.

35. Koike, N., Yoo, S.-H., Huang, H.-C., Kumar, V., Lee, C., Kim, T.-K., and Takahashi, J.S. (2012). Transcriptional Architecture and Chromatin Landscape of the Core Circadian Clock in Mammals. Science 338, 349–354. 10.1126/science.1226339.

36. Tartour, K., and Padmanabhan, K. (2022). The Clock Takes Shape—24 h Dynamics in Genome Topology. Front. Cell Dev. Biol. 9, 799971. 10.3389/fcell.2021.799971.

37. Xiao, Y., Yuan, Y., Jimenez, M., Soni, N., and Yadlapalli, S. (2021). Clock proteins regulate spatiotemporal organization of clock genes to control circadian rhythms. Proc. Natl. Acad. Sci. U.S.A. 118, e2019756118. 10.1073/pnas.2019756118.

38. Kalocsay, M., Hiller, N.J., and Jentsch, S. (2009). Chromosome-wide Rad51 Spreading and SUMO-H2A.Z-Dependent Chromosome Fixation in Response to a Persistent DNA Double-Strand Break. Molecular Cell 33, 335–343. 10.1016/j.molcel.2009.01.016.

39. Horigome, C., Unozawa, E., Ooki, T., and Kobayashi, T. (2019). Ribosomal RNA gene repeats associate with the nuclear pore complex for maintenance after DNA damage. PLoS Genet 15, e1008103. 10.1371/journal.pgen.1008103.

40. Horigome, C., Bustard, D.E., Marcomini, I., Delgoshaie, N., Tsai-Pflugfelder, M., Cobb, J.A., and Gasser, S.M. (2016). PolySUMOylation by Siz2 and Mms21 triggers relocation of DNA breaks to nuclear pores through the Slx5/Slx8 STUbL. Genes Dev. 30, 931–945. 10.1101/gad.277665.116.

41. Oshidari, R., Strecker, J., Chung, D.K.C., Abraham, K.J., Chan, J.N.Y., Damaren, C.J., and Mekhail, K. (2018). Nuclear microtubule filaments mediate non-linear directional motion of chromatin and promote DNA repair. Nat Commun 9, 2567. 10.1038/s41467-018-05009-7.

42. Chung, D.K.C., Chan, J.N.Y., Strecker, J., Zhang, W., Ebrahimi-Ardebili, S., Lu, T., Abraham, K.J., Durocher, D., and Mekhail, K. (2015). Perinuclear tethers license telomeric DSBs for a broad kinesin-and NPC-dependent DNA repair process. Nat Commun 6, 7742. 10.1038/ncomms8742.

43. Khadaroo, B., Teixeira, M.T., Luciano, P., Eckert-Boulet, N., Germann, S.M., Simon, M.N., Gallina, I., Abdallah, P., Gilson, E., Géli, V., et al. (2009). The DNA damage response at eroded telomeres and tethering to the nuclear pore complex. Nat Cell Biol 11, 980–987. 10.1038/ncb1910.

44. Pinzaru, A.M., Kareh, M., Lamm, N., Lazzerini-Denchi, E., Cesare, A.J., and Sfeir, A. (2020). Replication stress conferred by POT1 dysfunction promotes telomere relocalization to the nuclear pore. Genes Dev 34, 1619–1636. 10.1101/gad.337287.120.

45. Lamm, N., Read, M.N., Nobis, M., Van Ly, D., Page, S.G., Masamsetti, V.P., Timpson, P., Biro, M., and Cesare, A.J. (2020). Nuclear F-actin counteracts nuclear deformation and promotes fork repair during replication stress. Nat Cell Biol 22, 1460–1470. 10.1038/s41556-020-00605-6.

46. Caridi, C.P., D’Agostino, C., Ryu, T., Zapotoczny, G., Delabaere, L., Li, X., Khodaverdian, V.Y., Amaral, N., Lin, E., Rau, A.R., et al. (2018). Nuclear F-actin and myosins drive relocalization of heterochromatic breaks. Nature 559, 54–60. 10.1038/s41586-018-0242-8.

47. Mangeat, T., Labouesse, S., Allain, M., Negash, A., Martin, E., Guénolé, A., Poincloux, R., Estibal, C., Bouissou, A., Cantaloube, S., et al. (2021). Super-resolved live-cell imaging using random illumination microscopy. Cell Reports Methods 1, 100009. 10.1016/j.crmeth.2021.100009.

48. Lottersberger, F., Karssemeijer, R.A., Dimitrova, N., and de Lange, T. (2015). 53BP1 and the LINC Complex Promote Microtubule-Dependent DSB Mobility and DNA Repair. Cell 163, 880–893. 10.1016/j.cell.2015.09.057.

49. Liu, Q., Pante, N., Misteli, T., Elsagga, M., Crisp, M., Hodzic, D., Burke, B., and Roux, K.J. (2007). Functional association of Sun1 with nuclear pore complexes. Journal of Cell Biology 178, 785–798. 10.1083/jcb.200704108.

50. Li, P., and Noegel, A.A. (2015). Inner nuclear envelope protein SUN1 plays a prominent role in mammalian mRNA export. Nucleic Acids Res, gkv1058. 10.1093/nar/gkv1058.

51. Su, X.A., Dion, V., Gasser, S.M., and Freudenreich, C.H. (2015). Regulation of recombination at yeast nuclear pores controls repair and triplet repeat stability. Genes Dev 29, 1006–1017. 10.1101/gad.256404.114.

52. Caron, P., van der Linden, J., and van Attikum, H. (2019). Bon voyage: A transcriptional journey around DNA breaks. DNA Repair 82, 102686. 10.1016/j.dnarep.2019.102686.

53. Zhou, Y., Caron, P., Legube, G., and Paull, T.T. (2014). Quantitation of DNA double-strand break resection intermediates in human cells. Nucleic Acids Research 42, e19–e19. 10.1093/nar/gkt1309.

54. Galanty, Y., Belotserkovskaya, R., Coates, J., and Jackson, S.P. (2012). RNF4, a SUMO-targeted ubiquitin E3 ligase, promotes DNA double-strand break repair. Genes Dev 26, 1179–1195. 10.1101/gad.188284.112.

55. Han, M.M., Hirakawa, M., Yamauchi, M., and Matsuda, N. (2022). Roles of the SUMO-related enzymes, PIAS1, PIAS4, and RNF4, in DNA double-strand break repair by homologous recombination. Biochem Biophys Res Commun 591, 95–101. 10.1016/j.bbrc.2021.12.099.

56. Lieberman-Aiden, E., van Berkum, N.L., Williams, L., Imakaev, M., Ragoczy, T., Telling, A., Amit, I., Lajoie, B.R., Sabo, P.J., Dorschner, M.O., et al. (2009). Comprehensive Mapping of Long-Range Interactions Reveals Folding Principles of the Human Genome. Science 326, 289–293. 10.1126/science.1181369.

57. Falk, M., Feodorova, Y., Naumova, N., Imakaev, M., Lajoie, B.R., Leonhardt, H., Joffe, B., Dekker, J., Fudenberg, G., Solovei, I., et al. (2019). Heterochromatin drives compartmentalization of inverted and conventional nuclei. Nature 570, 395–399. 10.1038/s41586-019-1275-3.

58. Roukos, V., Voss, T.C., Schmidt, C.K., Lee, S., Wangsa, D., and Misteli, T. (2013). Spatial Dynamics of Chromosome Translocations in Living Cells. Science 341, 660–664. 10.1126/science.1237150.

59. Shokrollahi, M., Stanic, M., Hundal, A., Chan, J. N.Y., Urman, D., Hakem, A., Garcia, R.E., Hao, J., Maass, P.G., Dickson, B.C., et al. (2023). DNA double-strand break-capturing nuclear envelope tubules drive DNA repair. bioRxiv, 2023.05.07.539750. 10.1101/2023.05.07.539750.

60. Sancar, A., and Van Gelder, R.N. (2021). Clocks, cancer, and chronochemotherapy. Science 371, eabb0738. 10.1126/science.abb0738.

61. Kondratov, R.V., and Antoch, M.P. (2007). Circadian proteins in the regulation of cell cycle and genotoxic stress responses. Trends in Cell Biology 17, 311–317. 10.1016/j.tcb.2007.07.001.

62. Masri, S., and Sassone-Corsi, P. (2018). The emerging link between cancer, metabolism, and circadian rhythms. Nat Med 24, 1795–1803. 10.1038/s41591-018-0271-8.

63. Mure, L.S., Le, H.D., Benegiamo, G., Chang, M.W., Rios, L., Jillani, N., Ngotho, M., Kariuki, T., Dkhissi-Benyahya, O., Cooper, H.M., et al. (2018). Diurnal transcriptome atlas of a primate across major neural and peripheral tissues. Science 359, eaao0318. 10.1126/science.aao0318.

64. Alexandrou, A.T., Duan, Y., Xu, S., Tepper, C., Fan, M., Tang, J., Berg, J., Basheer, W., Valicenti, T., Wilson, P.F., et al. (2022). PERIOD 2 regulates low-dose radioprotection via PER2/pGSK3β/β-catenin/Per2 loop. iScience 25, 105546. 10.1016/j.isci.2022.105546.

65. Papp, S.J., Huber, A.-L., Jordan, S.D., Kriebs, A., Nguyen, M., Moresco, J.J., Yates, J.R., and Lamia, K.A. (2015). DNA damage shifts circadian clock time via Hausp-dependent Cry1 stabilization. eLife 4, e04883. 10.7554/eLife.04883.

66. Shafi, A.A., McNair, C.M., McCann, J.J., Alshalalfa, M., Shostak, A., Severson, T.M., Zhu, Y., Bergman, A., Gordon, N., Mandigo, A.C., et al. (2021). The circadian cryptochrome, CRY1, is a pro-tumorigenic factor that rhythmically modulates DNA repair. Nat Commun 12, 401. 10.1038/s41467-020-20513-5.

67. Anabtawi, N., Cvammen, W., and Kemp, M.G. (2021). Pharmacological inhibition of cryptochrome and REV-ERB promotes DNA repair and cell cycle arrest in cisplatin-treated human cells. Sci Rep 11, 17997. 10.1038/s41598-021-97603-x.

68. Gaddameedhi, S., Selby, C.P., Kaufmann, W.K., Smart, R.C., and Sancar, A. (2011). Control of skin cancer by the circadian rhythm. Proc. Natl. Acad. Sci. U.S.A. 108, 18790– 18795. 10.1073/pnas.1115249108.

69. Zhang, C., Chen, L., Sun, L., Jin, H., Ren, K., Liu, S., Qian, Y., Li, S., Li, F., Zhu, C., et al. (2023). BMAL1 collaborates with CLOCK to directly promote DNA double-strand break repair and tumor chemoresistance. Oncogene. 10.1038/s41388-023-02603-y.

70. Talamanca, L., Gobet, C., and Naef, F. (2023). Sex-dimorphic and age-dependent organization of 24-hour gene expression rhythms in humans. Science 379, 478–483. 10.1126/science.add0846.

71. Zada, D., Sela, Y., Matosevich, N., Monsonego, A., Lerer-Goldshtein, T., Nir, Y., and Appelbaum, L. (2021). Parp1 promotes sleep, which enhances DNA repair in neurons. Molecular Cell 81, 4979–4993.e7. 10.1016/j.molcel.2021.10.026.

72. Nassan, M., and Videnovic, A. (2022). Circadian rhythms in neurodegenerative disorders. Nat Rev Neurol 18, 7–24. 10.1038/s41582-021-00577-7.

73. Caron, P., Choudjaye, J., Clouaire, T., Bugler, B., Daburon, V., Aguirrebengoa, M., Mangeat, T., Iacovoni, J.S., Álvarez-Quilón, A., Cortés-Ledesma, F., et al. (2015). Non-redundant Functions of ATM and DNA-PKcs in Response to DNA Double-Strand Breaks. Cell Reports 13, 1598–1609. 10.1016/j.celrep.2015.10.024.

74. Natsume, T., Kiyomitsu, T., Saga, Y., and Kanemaki, M.T. (2016). Rapid Protein Depletion in Human Cells by Auxin-Inducible Degron Tagging with Short Homology Donors. Cell Rep 15, 210–218. 10.1016/j.celrep.2016.03.001.

75. Nemoz, C., Ropars, V., Frit, P., Gontier, A., Drevet, P., Yu, J., Guerois, R., Pitois, A., Comte, A., Delteil, C., et al. (2018). XLF and APLF bind Ku80 at two remote sites to ensure DNA repair by non-homologous end joining. Nat Struct Mol Biol 25, 971–980. 10.1038/s41594-018-0133-6.

76. Fu, C., Donovan, W.P., Shikapwashya-Hasser, O., Ye, X., and Cole, R.H. (2014). Hot Fusion: an efficient method to clone multiple DNA fragments as well as inverted repeats without ligase. PLoS One 9, e115318. 10.1371/journal.pone.0115318.

77. Cheng, Q., Barboule, N., Frit, P., Gomez, D., Bombarde, O., Couderc, B., Ren, G.-S., Salles, B., and Calsou, P. (2011). Ku counteracts mobilization of PARP1 and MRN in chromatin damaged with DNA double-strand breaks. Nucleic Acids Res 39, 9605–9619. 10.1093/nar/gkr656.

78. Li, Q. (2004). A Syntaxin 1, G o, and N-Type Calcium Channel Complex at a Presynaptic Nerve Terminal: Analysis by Quantitative Immunocolocalization. Journal of Neuroscience 24, 4070–4081. 10.1523/JNEUROSCI.0346-04.2004.

79. van Steensel, B., van Binnendijk, E.P., Hornsby, C.D., van der Voort, H.T., Krozowski, Z.S., de Kloet, E.R., and van Driel, R. (1996). Partial colocalization of glucocorticoid and mineralocorticoid receptors in discrete compartments in nuclei of rat hippocampus neurons. Journal of Cell Science 109, 787–792. 10.1242/jcs.109.4.787.

80. Arnould, C., Rocher, V., Finoux, A.-L., Clouaire, T., Li, K., Zhou, F., Caron, P., Mangeot, Philippe.E., Ricci, E.P., Mourad, R., et al. (2021). Loop extrusion as a mechanism for formation of DNA damage repair foci. Nature 590, 660–665. 10.1038/s41586-021-03193-z.

