## Extended Figures and Tables for "Circadian PERIOD proteins regulate TC-DSB repair through anchoring to the nuclear envelope"

1. *MCD, Centre de Biologie Intégrative (CBI), CNRS, Université de Toulouse, UT3, 31062 Toulouse, France*
2. *BigA, NGS data analysis facility, Centre de Biologie Intégrative (CBI), CNRS, Université de Toulouse, UT3, 31062 Toulouse, France*
3. *Institut de Pharmacologie et de Biologie Structurale (IPBS), Université de Toulouse, CNRS, Université Toulouse III - Paul Sabatier (UT3), Toulouse, France*
4. *Equipe Labellisée Ligue Contre le Cancer 2018, Toulouse, France*
5. *LITC Core Facility, Centre de Biologie Intégrative (CBI), CNRS, Université de Toulouse, UT3, 31062 Toulouse, France*
6. *Lead author*

**Running title:** Circadian regulation of TC-DSB repair

#### Extended Data Figures 1-8

#### Extended Data Figure Legends 1-8

#### Extended Data Tables 1-2

Le Bozec et al, Extended data Figure 1

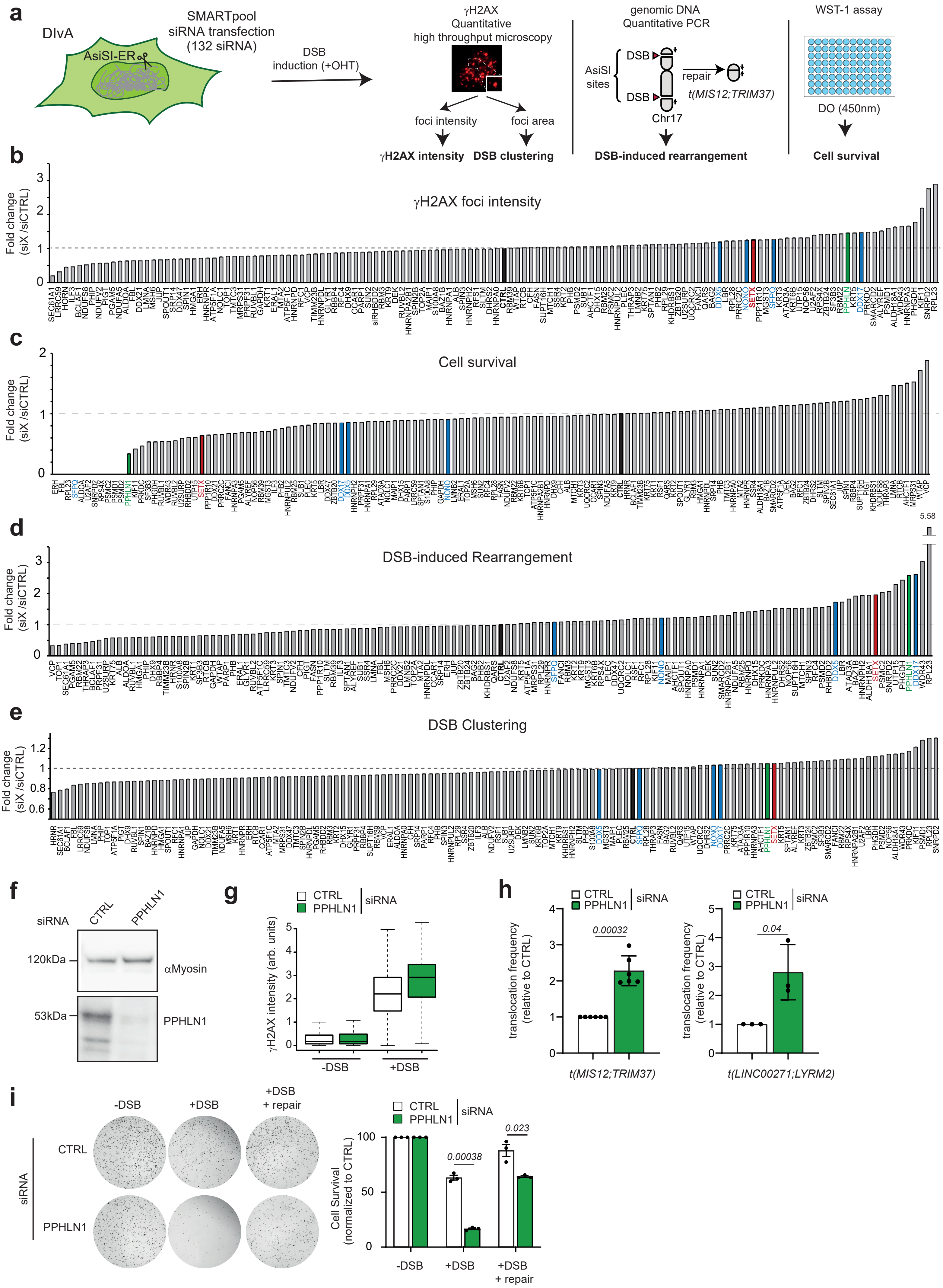

### Le Bozec et al, Extended data Figure 2

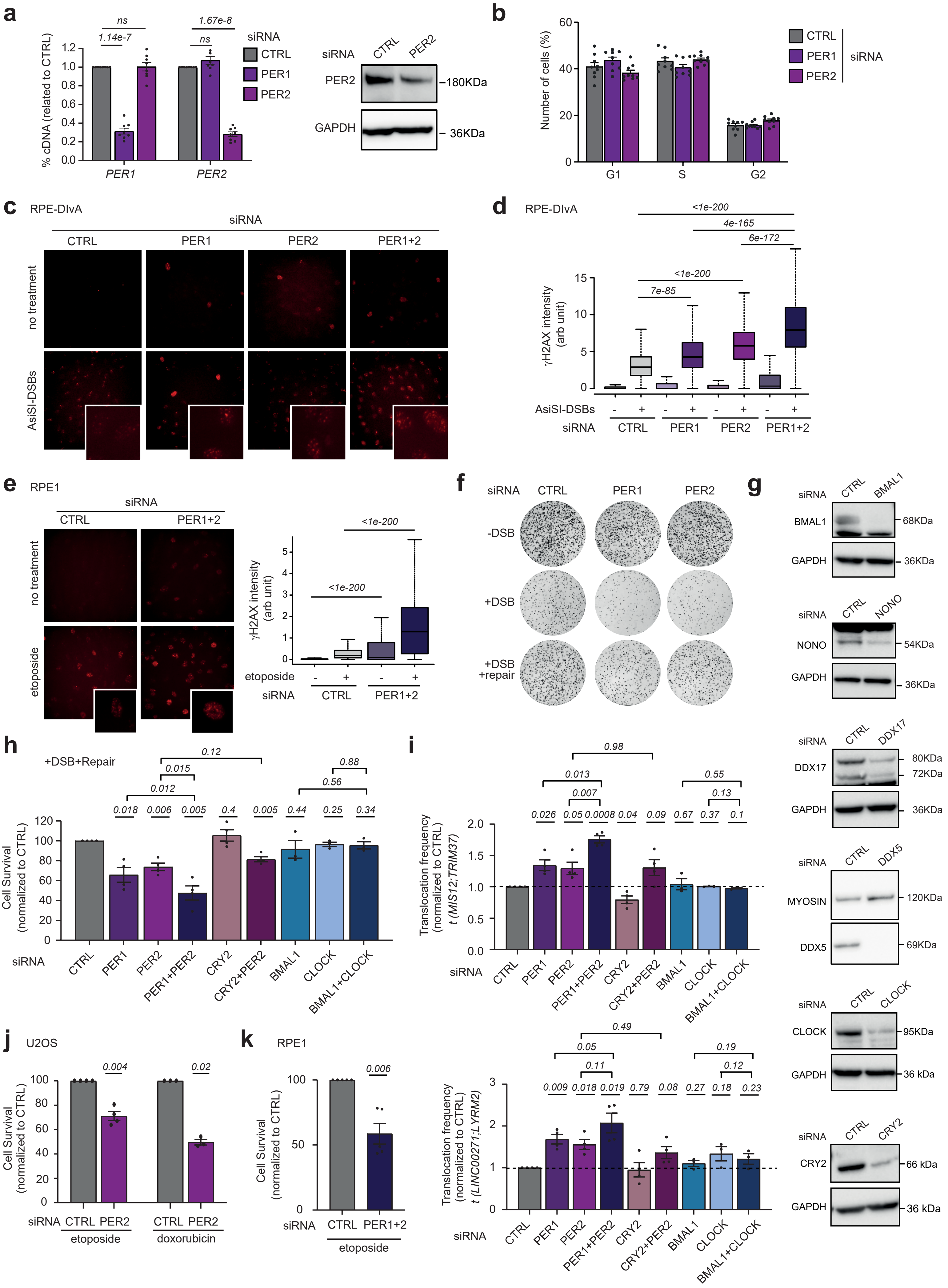

Le Bozec et al, Extended data Figure 3

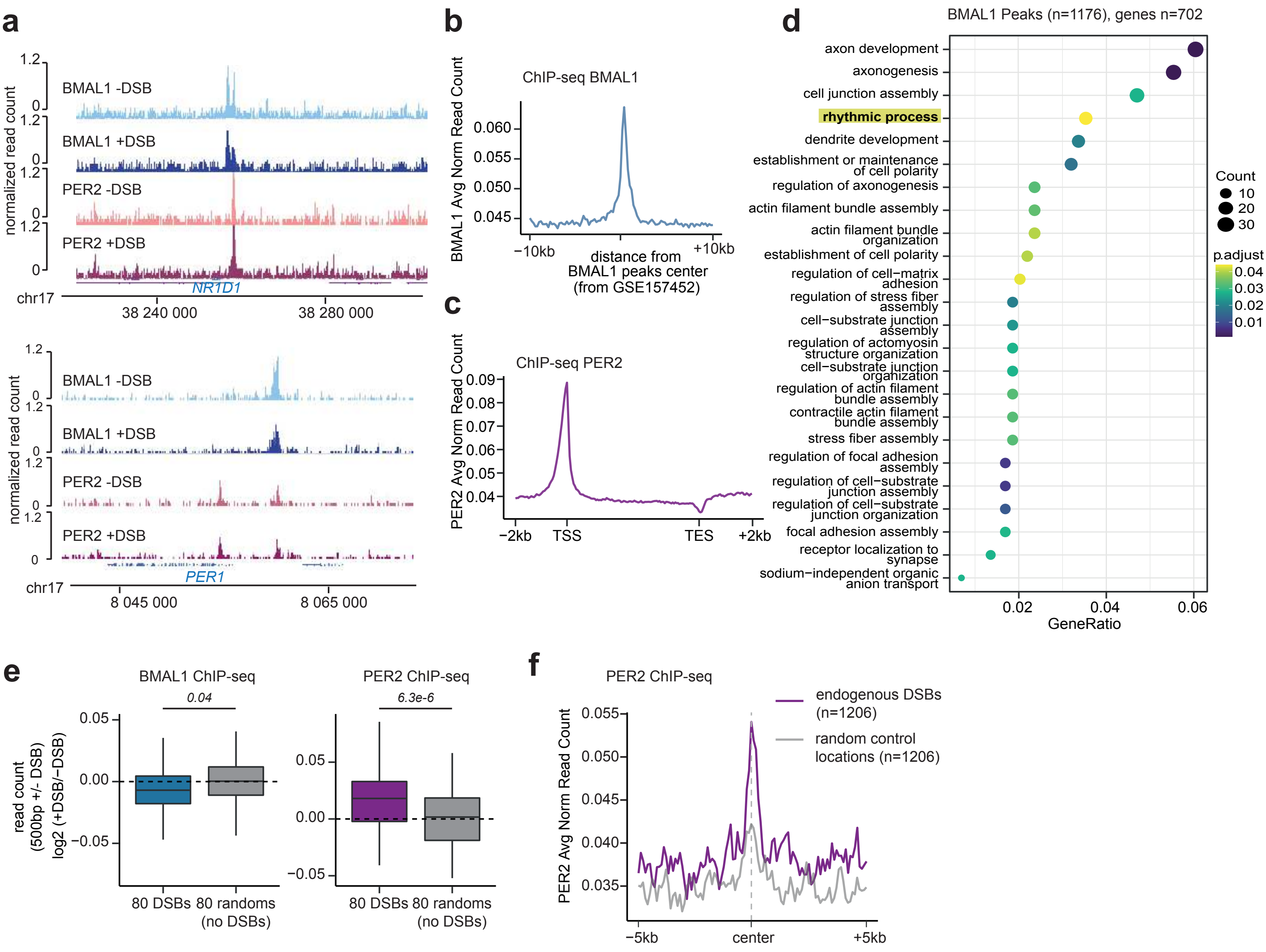

### Le Bozec et al, Extended data Figure 4

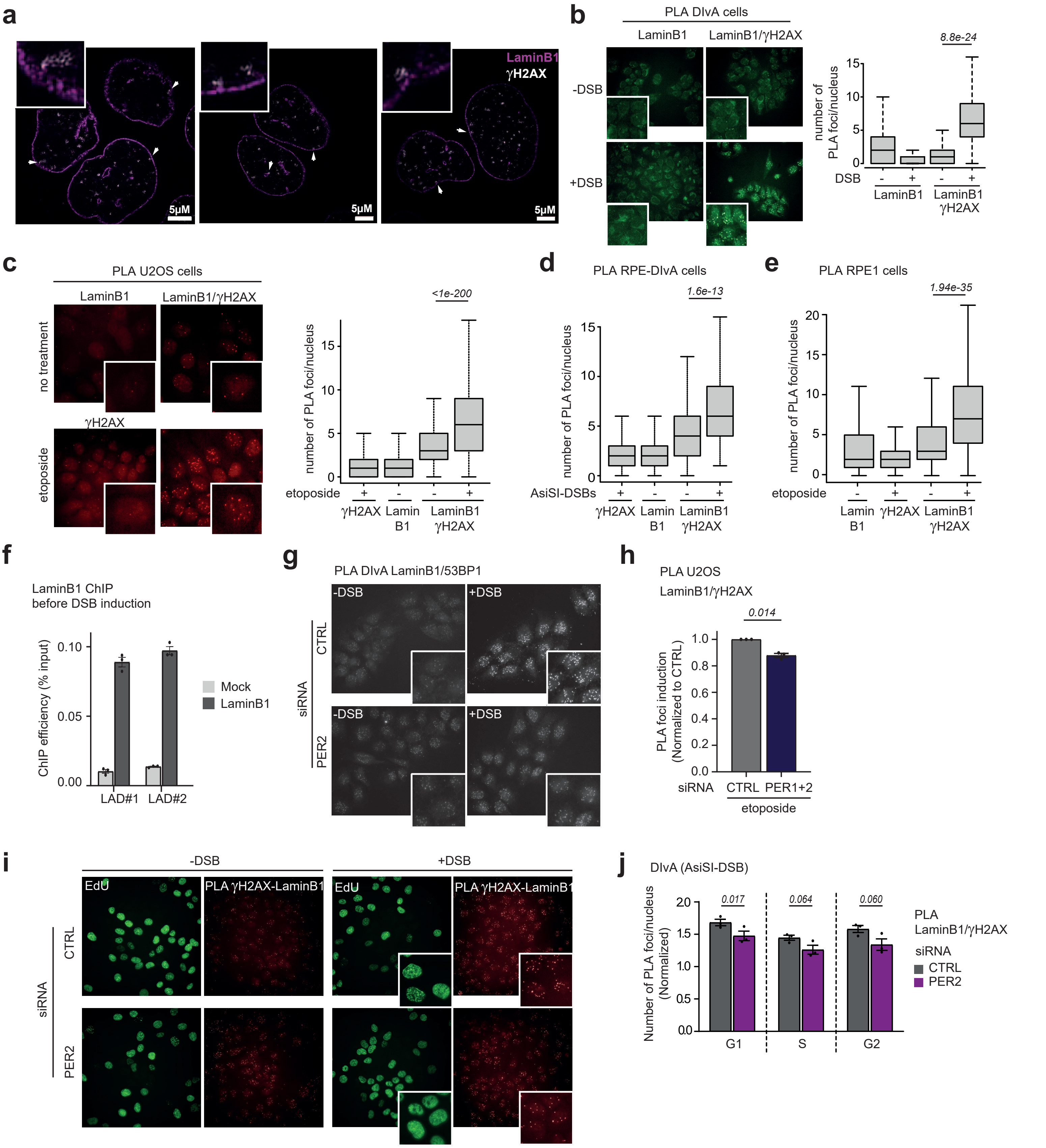

Le Bozec et al, Extended data Figure 5

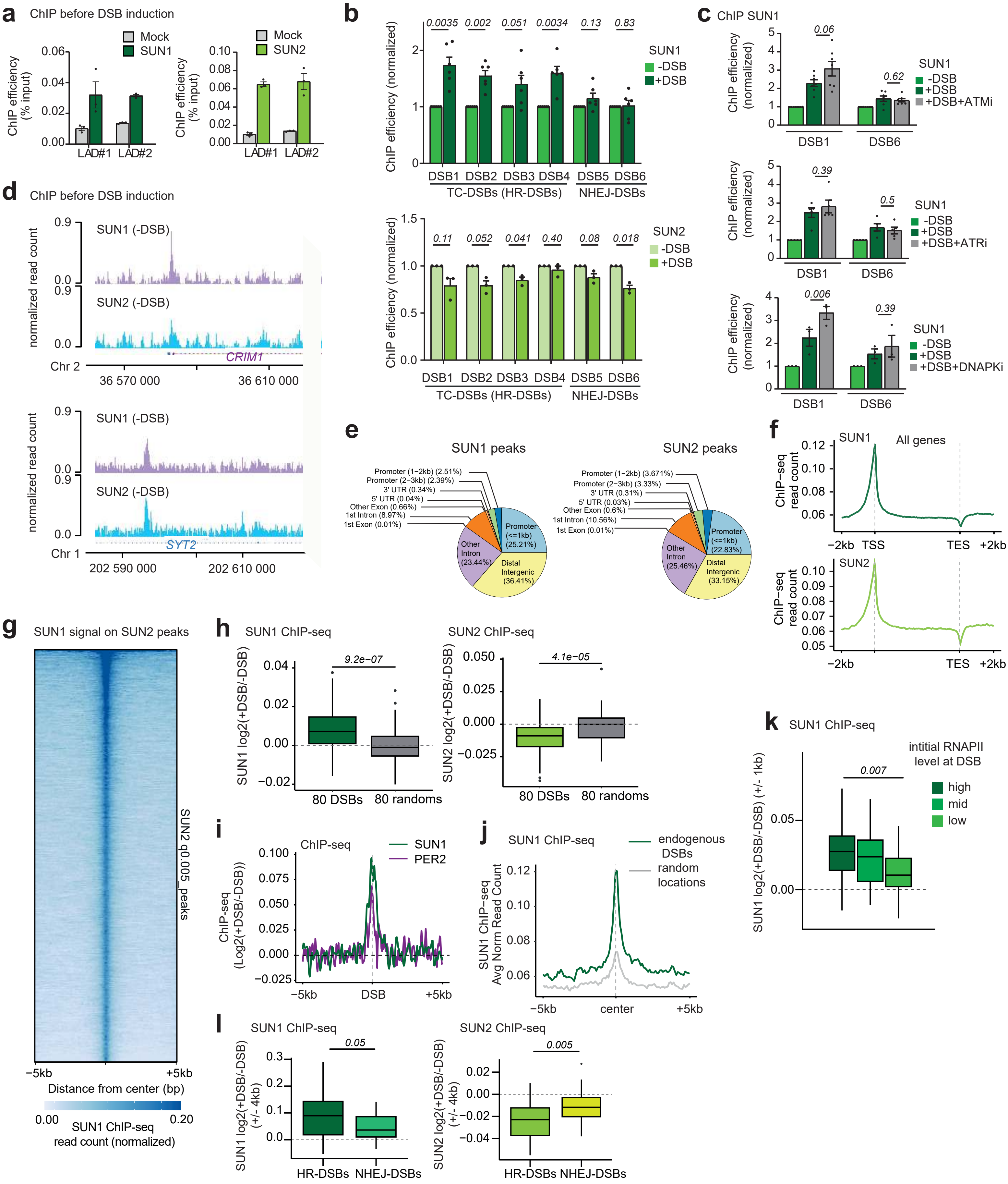

Le Bozec et al, Extended data Figure 6

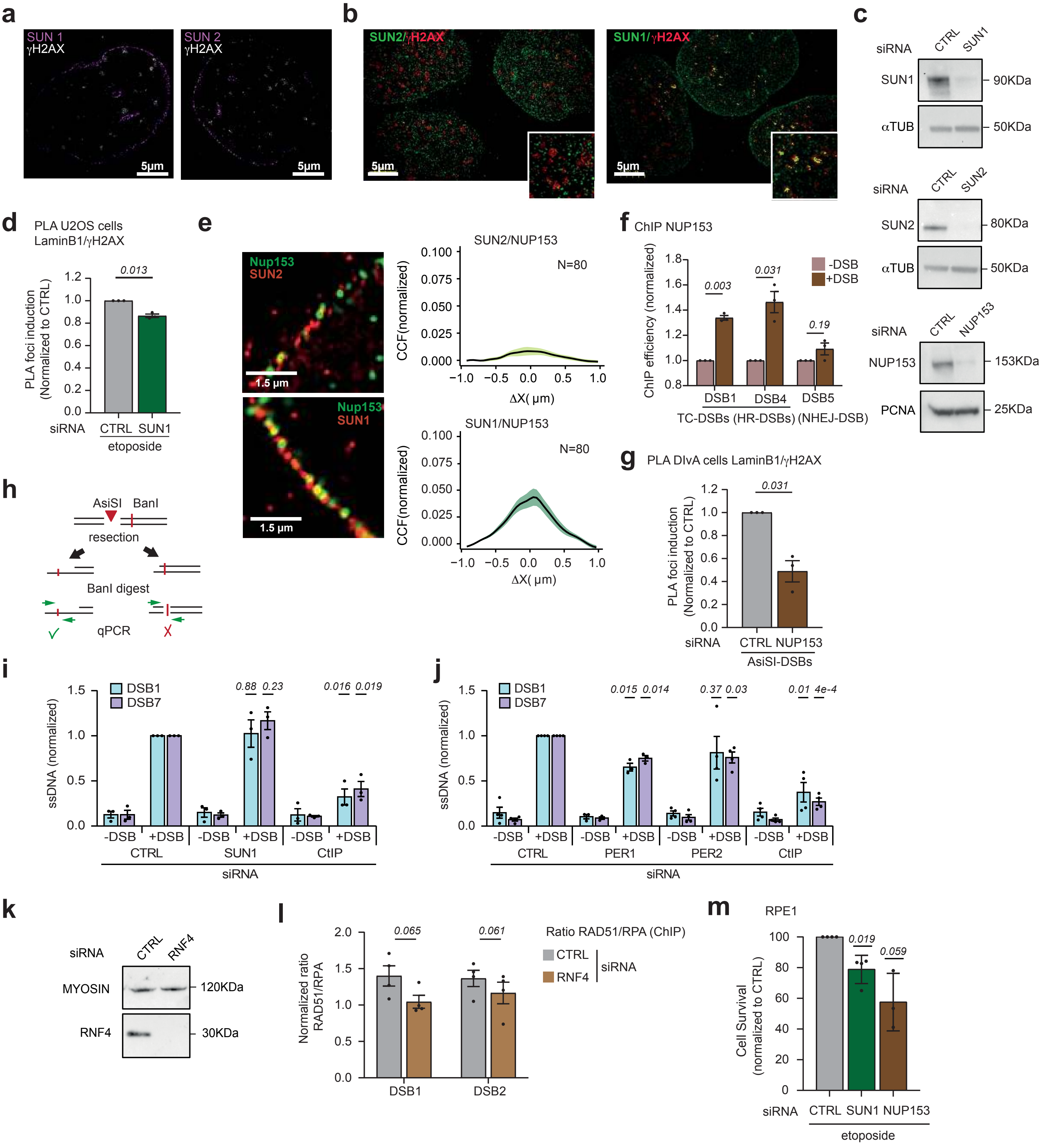

### Le Bozec et al, Extended data Figure 7

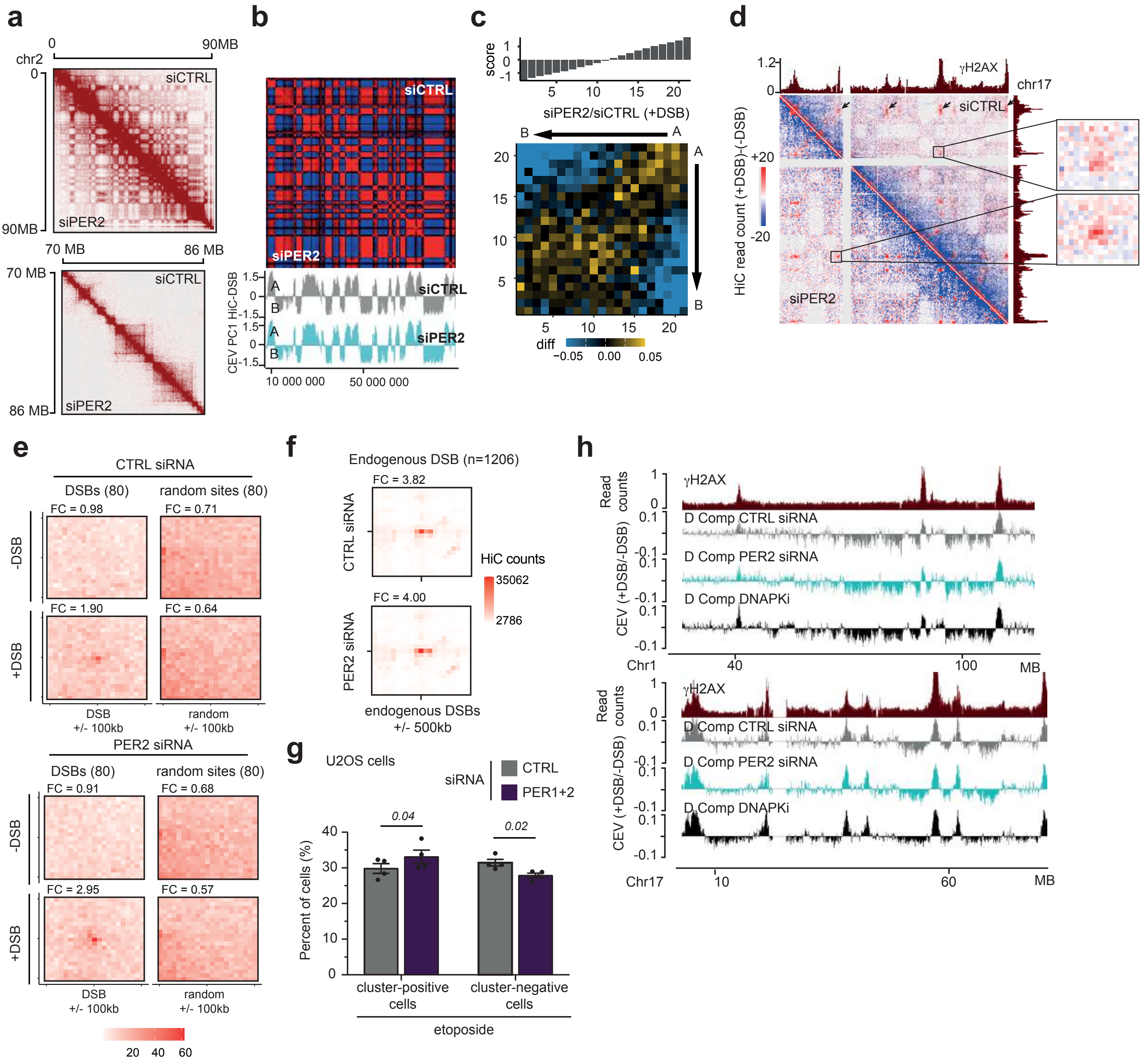

Le Bozec et al, Extended data Figure 8

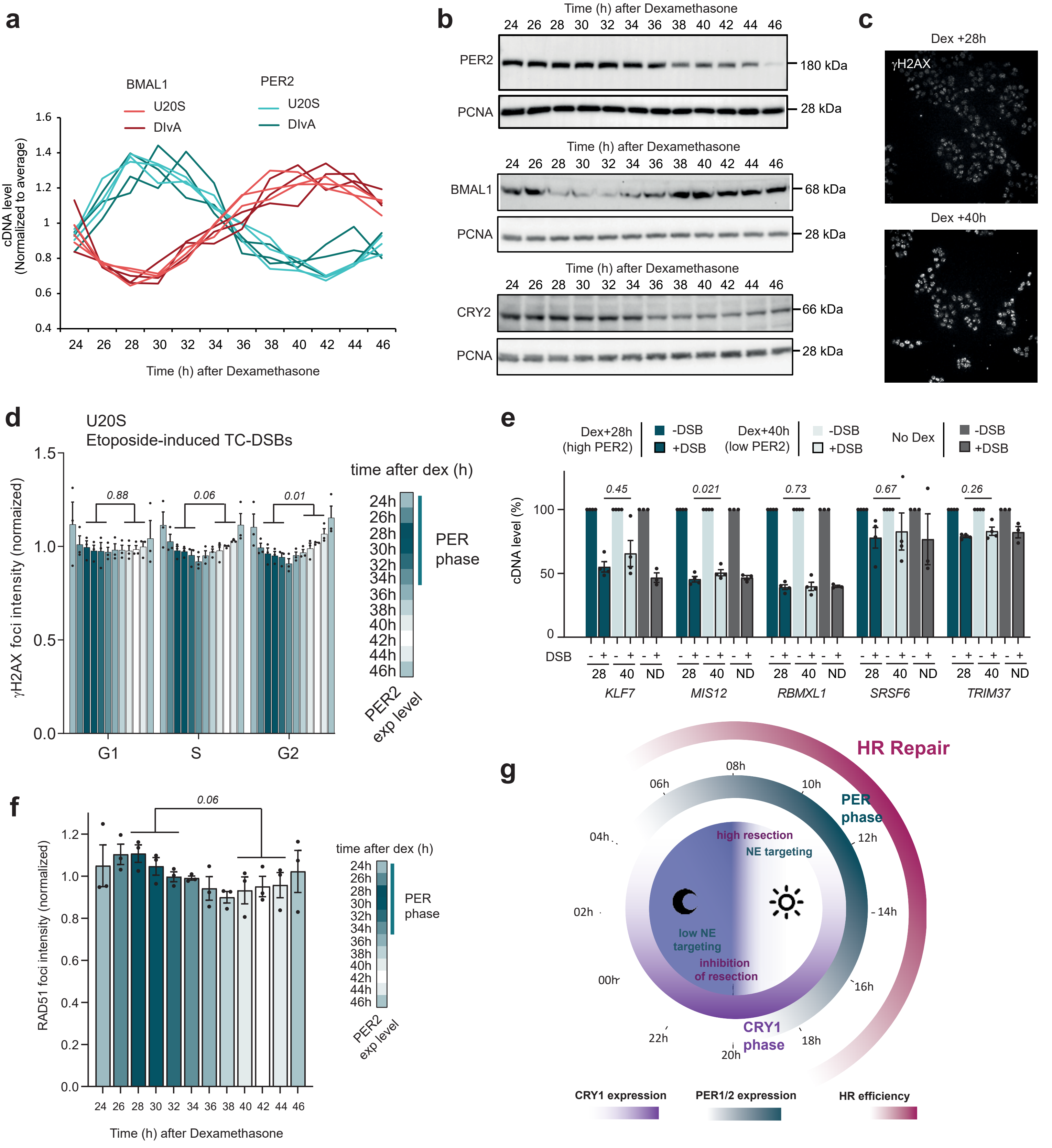

### Extended Data Figure Legends

#### Extended Data Figure 1: Multiscreen validation.

**a**. Schematic steps for multi-output screening of a TC-DSBR-focused siRNA library. Following siRNA transfection of AID-DIvA cells, four different outputs were analyzed before and after DSB induction (+OHT). The γH2AX signal was quantified with high-throughput microscopy and both the γH2AX foci intensity and the γH2AX foci mean area were measured in >5000 nuclei per condition to obtain respectively γH2AX intensity and DSB clustering data. Genomic DNA was extracted after DSB induction and repair (+OHT+IAA), and quantitative PCR using primers (small arrows) located on only one side of each of two distant AsiSI sites on chromosome 17 was used to detect their illegitimate rejoining and obtain the frequency of the *t(MIS12;TRIM37)* DSB-induced rearrangement. Eight days after DSB induction, the cells were incubated with tetrazolium salts (WST-1 cell viability assay) and formazan dye production formed by metabolically active cell was quantified by measuring its absorbance (450nm) to assess the cell survival.

**b-e**. Results from the multi-output screen in AID-DIvA cells using 132 SMARTpool siRNAs. Data are expressed normalized to the control siRNA (CTRL, black bars). A positive control (SETX) was included in the experiment (red bars). PPHLN1 and PER complex siRNAs are highlighted in green and blue respectively. **b.** γH2AX foci intensity was scored in siRNA transfected cells, using quantitative high throughput imaging. **c.** Cell survival of siRNA transfected cells was quantified eight days after DSB induction, using a colorimetric assay based on mitochondrial dehydrogenase activity in viable cells. **d.** *t(MIS12;TRIM37)* rearrangement frequency was analyzed by qPCR following DSB induction and repair. **e.** γH2AX foci mean area (DSB clustering) was scored in siRNA transfected cells, using quantitative high throughput imaging.

**f.** Western blots after control (CTRL) or PPHLN1 siRNA knockdown in AID-DIvA cells (n=2).

**g**. γH2AX foci intensity performed in control (CTRL) and PPHLN1 siRNA transfected DIvA cells as indicated, before (−DSB) and after (+DSB) DSB induction. A representative experiment (>7000 nuclei) is shown. Center line: median; box limits: 1st and 3rd quartiles; whiskers: maximum and minimum without outliers. *P*, unpaired t-test.

**h**. *t(MIS12;TRIM37)* (left panel) and *t(LINC00271:LYRM2)* (right panel) rejoining frequencies after OHT and IAA treatment analyzed by quantitative PCR in AID-DIvA cells transfected with control (CTRL) and PPHLN1 siRNA. Mean and SEM (n≥3 biological replicates) are shown. *P*, paired t-test (two-sided).

**i.** Clonogenic assays in AID-DIvA cells transfected with control (CTRL) and PPHLN1 siRNA, before (−DSB) and after (+DSB) DSB induction and after IAA treatment (+DSB+repair) as indicated. Left panel shows a representative experiment and right panel shows the mean and SEM of cell survival (n=3 biological replicates). *P*, paired t-test (two-sided).

**Extended Data Figure 2: PER protein knockdown decreases cell survival and increases translocations upon DSB induction.**

**a**. Left panel: cDNA levels analyzed by RT-qPCR in AID-DIvA cells transfected with a control siRNA (CTRL) and PER1 or PER2 siRNAs. Mean and SEM are shown for n=8 biological replicates. *P*, paired t-test (two-sided). Right panel: Western Blot after control (CTRL) or PER2 siRNA knockdown in DivA cells (n=2).

**b**. Cell cycle analysis of DivA cells after transfection with a control siRNA (CTRL) and PER1 or PER2 siRNAs. Mean and SEM are shown for n=9 biological replicates.

**c**. Examples of γH2AX staining performed in control (CTRL), PER1, PER2 or PER1+PER2 (PER1+2) siRNA-transfected RPE-DIvA cells, before (no treatment) and after AsiSI-induced DSBs.

**d.** Quantification of γH2AX intensity in RPE-DIvA cells is shown (>1000 nuclei, representative experiment, n=4 biological replicates for PER1+2). Center line: median; box limits: 1st and 3rd quartiles; whiskers: maximum and minimum without outliers. *P*, Wilcoxon-Mann-Whitney test.

**e**. Left panel: examples of γH2AX staining performed in control (CTRL) or PER1+PER2 (PER1+2) siRNAs-transfected RPE1 cells before (no treatment) and after etoposide treatment. Right panel: Quantification of γH2AX intensity before (-) and after (+) etoposide-induced DSBs is shown (>6500 nuclei, representative of n=4 biological replicates). Center line: median; box limits: 1st and 3rd quartiles; whiskers: maximum and minimum without outliers. *P*, Welch test.

**f.** Clonogenic assays in AID-DIvA cells transfected with control (CTRL), PER1 or PER2 siRNA, before (−DSB) and after (+DSB) DSB induction and after IAA treatment (+DSB+repair) as indicated. A representative experiment is shown.

**g**. Western Blots after control (CTRL) or BMAL1, NONO, DDX17, DDX5, CLOCK and CRY2 siRNA knockdown in AID-DIvA cells (n=2).

**h.** Clonogenic assays in AID-DIvA cells transfected with control (CTRL), PER1, PER2, PER1+PER2, CRY2, CRY2+PER2, BMAL1, CLOCK or BMAL1+CLOCK siRNAs. Mean and SEM after DSB induction and repair (n≥3). *P*, paired t-test (two-sided).

**i.** *t*(*MIS12:TRIM37)* (upper panel) and *t(LINC00271:LYRM2)* (lower panel) rejoining frequencies after DSB induction and repair measured by qPCR in AID-DIvA cells transfected with control (CTRL), PER1, PER2, PER1+PER2, CRY2, CRY2+PER2, BMAL1, CLOCK or BMAL1+CLOCK siRNAs. Mean and SEM (n≥3) are shown. *P*, paired t-test (two-sided).

**j.** Clonogenic assays in U2OS cells transfected with control (CTRL) or PER2 siRNA. Mean and SEM after etoposide or doxorubicin treatment (n≥3). *P*, paired t-test (two-sided).

**k.** Clonogenic assays in RPE1 cells transfected with control (CTRL) or PER1+PER2 (PER1+2) siRNAs. Mean and SEM after etoposide treatment (n=5). *P*, paired t-test (two-sided).

**Extended Data Figure 3: PER2 binds to induced DSBs, unlike BMAL1.**

**a**. Genomic tracks showing ChIP-seq against BMAL1 (blue) and PER2 (pink-violet) at *NR1D1* (upper panel) and *PER1* (bottom panel) promoters before (−DSB) and after (+DSB) DSB induction.

**b**. Average profile of BMAL1 ChIP-seq signal in absence of DSB induction, centered on BMAL1 peaks on a ±10kb window.

**c**. Average profile of PER2 ChIP-seq signal on a ±2kb window across all protein coding genes on the genome (hg19) in absence of DSB induction. TSS: transcription start site; TES: transcription end site.

**d**. BMAL1 ChIP-seq peak data in absence of DSB induction, analyzed for enrichment of Gene Ontology terms. Exact *P*-value (Fisher test) was adjusted (Benjamini-Hochberg correction).

**e**. Box plots representing BMAL1 (left panel) and PER2 (right panel) enrichment (log2(+DSB/−DSB)) from ChIP-seq signal in a 1kb window centered on the eighty best-induced AsiSI-DSBs (80 DSBs) or eighty random sites (80 randoms (no DSBs)). Center line: median; box limits: 1st and 3rd quartiles; whiskers: maximum and minimum without outliers. *P*, paired Wilcoxon test.

**f**. Average PER2 ChIP-seq profile on a ±5 kb window centered on n=1206 endogenous DSBs (violet) and n=1206 random positions (light grey).

**Extended Data Figure 4: TC-DSBs interact with LaminB1.**

**a**. Super-resolution imaging, performed using Random Illumination Microscopy (RIM), showing γH2AX foci (white) and LaminB1 (purple) after DSB induction in DIvA cells. Arrows show close contact between DSBs and nuclear laminB1.

**b**. Left panels: PLA performed using either LaminB1 antibody alone (left) or both LaminB1 and γH2AX antibodies (right) before (−DSB) and after DSB (+DSB) induction in DIvA cells. Right panel: Box plots showing PLA foci per nucleus quantification (center line: median; box limits: 1$^{st}$ and 3$^{rd}$ quartiles; whiskers: maximum and minimum without outliers). An average of 230 cells were analyzed per condition (n=3). *P*, Wilcoxon test.

**c**. Left panels: PLA performed using either LaminB1 or γH2AX antibody alone (left) or both LaminB1 and γH2AX antibodies (right) before (no treatment) and after DSB induction (etoposide) in U2OS cells. Right panel: Box plots showing PLA foci per nucleus quantification before (-) and after (+) etoposide-induced DSBs (>3000 nuclei, representative of n=3 biological replicates). Center line: median; box limits: 1$^{st}$ and 3$^{rd}$ quartiles; whiskers: maximum and minimum without outliers. *P*, Welch test.

**d**. PLA foci per nucleus quantification performed using either LaminB1 or γH2AX antibody alone or both LaminB1 and γH2AX antibodies before (-) and after (+) AsiSI-induced DSBs in RPE-DIvA cells (>200 nuclei, representative of n=3 biological replicates). Center line: median; box limits: 1st and 3rd quartiles; whiskers: maximum and minimum without outliers. *P*, Wilcoxon-Mann-Whitney test.

**e**. PLA foci per nucleus quantification performed using either LaminB1 or γH2AX antibody alone or both LaminB1 and γH2AX antibodies before (-) and after (+) etoposide-induced DSBs in RPE1 cells (>400 nuclei, representative of n=3 biological replicates). Center line: median; box limits: 1st and 3rd quartiles; whiskers: maximum and minimum without outliers. *P*, Wilcoxon-Mann-Whitney test.

**f**. LaminB1 ChIP efficiency (% of input immunoprecipitated) before DSB induction in DIvA cells, at two Lamina-Associated Domains (LADs). ChIP without antibody (mock) and mean and SEM for n=3 biological replicates are shown.

**g**. PLA performed between LaminB1 and 53BP1 in control (CTRL) or PER2 siRNA-transfected DIvA cells as indicated, before (−DSB) and after (+DSB) AsiSI-DSB induction. A representative experiment is shown (n=3).

**h**. PLA performed between LaminB1 and γH2AX antibodies in control (CTRL) or PER1+PER2 (PER1+2) siRNA-transfected U2OS cells after etoposide-induced DSBs. Quantification of PLA foci per nucleus following etoposide is normalized to untreated and to CTRL. Mean and SEM (n=3) are shown. *P*, paired t-test (two-sided).

**i**. PLA performed between LaminB1 and γH2AX (red foci) in control (CTRL) or PER2 siRNA-transfected DIvA cells as indicated, before (−DSB) and after (+DSB) AsiSI-DSB induction. The EdU incorporation (green nuclei) and Hoechst staining (not shown) allow to determine G1, S, and G2 cells. A representative experiment is shown (n=3).

**j**. Number of PLA foci (γH2AX-LaminB1) per nucleus following AsiSI-DSB induction in DIvA cells transfected with control (CTRL) or PER2 siRNA, in G1, S, and G2 cells (>7000 nuclei were acquired in each condition for each biological replicate). Data were normalized to untreated cells (-DSB). Mean and SEM (n=3 biological replicates) are shown. *P*, Wilcoxon test.

**Extended Data Figure 5: TC-DSBs interact with SUN1, but not SUN2.**

**a**. SUN1 (left panel) and SUN2 (right panel) ChIP efficiency (% of input immunoprecipitated) before DSB induction in DIvA cells, at two LADs. ChIP without antibody (mock) and mean and SEM are shown for n=3 biological replicates.

**b**. SUN1 (upper panel) and SUN2 (bottom panel) ChIP efficiency before (−DSB) and after (+DSB) DSB induction, at four TC-DSBs (HR-DSBs; DSB1-4) and two NHEJ-DSBs (DSB5-6). Data are normalized to a control location devoid of DSB and further expressed related to data obtained in undamaged condition. Mean and SEM are shown for n≥3 biological replicates. *P*, paired t-test (two-sided).

**c**. SUN1 ChIP efficiency before (−DSB) and after DSB induction without (+DSB) or with ATM inhibition (+DSB+ATMi; upper panel), or ATR inhibition (+DSB+ATRi; middle panel), or DNA-PK inhibition (+DSB+DNAPKi; lower panel), at one TC-DSB (HR-DSB; DSB1) and one DSB in a silent locus (NHEJ-DSB; DSB6). Data are normalized to a control location devoid of DSB and further expressed related to data obtained in undamaged condition. Mean and SEM are shown for n=7 (ATMi), n=5 (ATRi) and n=3 (DNAPKi) biological replicates. *P*, paired t-test (two-sided).

**d**. Genomic tracks showing normalized read count of ChIP-seq against SUN1 (purple) and SUN2 (blue) before (−DSB) DSB induction at *CRIM1* (upper panel) and *SYT2* (bottom panel) loci.

**e**. Peak calling on SUN1 (left panel) and SUN2 (right panel) ChIP-seq data before DSB induction. The percentages on the pie charts show the genomic locations of SUN1 and SUN2 peaks.

**f**. Average profiles of SUN1 (top panel) and SUN2 (bottom panel) ChIP-seq signal before DSB induction on a ±2kb window across all protein coding genes on the genome (hg19). TSS: transcription start site; TES: transcription end site.

**g**. Heatmap representing the SUN1 ChIP-seq signal (normalized read count) on a 10kb window around SUN2 peaks, ordered based on the SUN2 ChIP-seq signal level, in DIvA cells before DSB induction.

**h**. Box plots representing SUN1 (left panel) and SUN2 (right panel) ChIP-seq enrichment (log2(+DSB/−DSB)) in a 1kb window centered on the eighty best-induced AsiSI-DSBs or eighty random sites (no DSBs). Center line: median; box limits: 1st and 3rd quartiles; whiskers: maximum and minimum. *P*, paired Wilcoxon test.

**i**. Average SUN1 (green) and PER2 (violet) ChIP-seq profiles on a ±5 kb window centered on the eighty best-induced AsiSI-DSBs. Data are presented in log2 (+DSB/-DSB).

**j**. Average SUN1 ChIP-seq profile on a ±5 kb window centered on n=1206 endogenous DSBs (green) and n=1206 random positions (light grey).

**k**. Box plot representing Log2 (+DSB/-DSB) SUN1 ChIP-seq on ±1 kb window around DSBs categorized based on their RNAPII level prior to DSB induction. Center line: median; box limits: 1st and 3rd quartiles; whiskers: maximum and minimum without outliers. *P*, Wilcoxon test.

**l**. Box plots representing Log2 (+DSB/-DSB) SUN1 (left panel) or SUN2 (right panel) ChIP-seq count on a ±4kb at HR-DSBs and NHEJ-DSBs (30 in each category). Center line: median; box limits: 1st and 3rd quartiles; whiskers: maximum and minimum without outliers. *P*, Wilcoxon test.

**Extended Data Figure 6: Tethering of TC-DSBs to the NE depends on SUN1 and NUP153 and DSB-anchoring at the NPC promotes HR.**

**a**. SUN1 (left) or SUN2 (right) staining (in purple) performed in DIvA cells after DSB induction (γH2AX foci in white) and acquired using RIM Super-resolution microscope.

**b**. γH2AX staining (in red) combined with the staining (in green) of SUN2 (left panel) or SUN1 (right panel) performed in DIvA cells after DSB induction and acquired using 3D RIM Super-resolution microscope. The areas represented in yellow (ICQ > 0.45) correspond to a high non-random colocalization (see methods).

**c**. Western Blots after control (CTRL) or SUN1 (top panel), SUN2 (middle panel) and NUP153 (bottom panel) siRNA knockdown in DIvA cells (n=2).

**d**. Quantification of LaminB1 and γH2AX PLA nuclear foci induction in control (CTRL) or SUN1-siRNA transfected U2OS cells following etoposide-induced DSBs, normalized to untreated and to CTRL. Mean and SEM are shown (n=3). *P*, paired t-test (two-sided).

**e.** Left panels: Magnifications of SUN2 (top panel) and SUN1 (bottom panel) staining (in red) together with NUP153 foci (in green). Right panels: Cross Correlation Function (CCF) plots between NE-embedded SUN2 (top panel) or SUN1 (bottom panel) and NUP153 foci (N=80). CCF of bicolor 3D RIM images was determined by plotting the value of Pearson's coefficient against Δx for each voxel of the image. The plots show a strong non-random overlap between the signals of SUN1 and NUP153 (peak at Δx=0).

**f.** NUP153 ChIP efficiency (% of input immunoprecipitated) before (−DSB) and after (+DSB) DSB induction in DIvA cells, at two TC-DSBs (HR-DSBs, DSB1,4) and one DSB in a silent locus (NHEJ-DSB, DSB5). Data are normalized to a control location devoid of DSB and further expressed related to data obtained in undamaged condition. Mean and SEM are shown for n=3 biological replicates. *P*, paired t-test (two-sided).

**g.** Quantification of LaminB1 and γH2AX nuclear PLA foci induction in control (CTRL) or NUP153-siRNA transfected DIvA cells following AsiSI-induced DSBs (+DSB), normalized to untreated and to CTRL. Mean and SEM (n=3 biological replicates) are shown. *P*, paired t-test (two-sided).

**h.** Resection assay schematic representation.

**i.** Quantification of single-strand DNA before (−DSB) and after (+DSB) AsiSI-DSB induction in control (CTRL), SUN1 or CtIP siRNA-depleted DIvA cells at two TC-DSBs (HR-DSBs, DSB1,7), normalized to CTRL. Mean and SEM are shown for n=3 biological replicates. *P*, paired t-test (two-sided).

**j.** Quantification of single-strand DNA before (−DSB) and after (+DSB) AsiSI-DSB induction in control (CTRL), PER1, PER2 or CtIP siRNA-depleted DIvA cells at two TC-DSBs (HR-DSBs, DSB1,7), normalized to CTRL. Mean and SEM are shown for n=3 (PER1) and n=4 (PER2, CtIP) biological replicates. *P*, paired t-test (two-sided).

**k.** Western Blots after control (CTRL) or RNF4 siRNA knockdown in DIvA cells (n=3).

**l.** Ratios of RAD51 over RPA ChIP-qPCR results in DIvA cells transfected with control (CTRL) or RNF4 siRNA after DSB induction, at two TC-DSBs (DSB1-2). Data are normalized to a control location devoid of a DSB. Mean and SEM are shown for n=4 biological replicates. *P*, paired t-test (two-sided).

**m.** Clonogenic assays in RPE1 cells transfected with CTRL, SUN1 or NUP153 siRNAs. Data were normalized to the untreated samples, and further expressed relative to CTRL. Mean and SEM after etoposide treatment (n=4 biological replicates). *P*, paired t-test (two-sided).

**Extended Data Figure 7: PER2 decreases A/B compartmentalization and counteracts DSB clustering and D-compartment formation.**

**a**. HiC contact matrix obtained on chromosome 2 in undamaged DIvA cells transfected with a control siRNA (siCTRL) or a siRNA directed against PER2 as indicated. Top panel, 250kb resolution, bottom panel, 25kb resolution.

**b**. Pearson correlation matrix (top panel) and Chromosomal Eigen vectors (CEV) showing A/B compartments (bottom panel), in undamaged DIvA cells transfected with a control siRNA (siCTRL) or a siRNA directed against PER2 as indicated.

**c**. Differential saddle plot between siPER2 and siCTRL showing A-B interactions after DSB induction (100kb bin).

**d**. Differential Hi-C contact matrix (+DSB)-(-DSB) on chromosome 17 (250kb resolution) in DIvA cells transfected with a control (CTRL) siRNA or a siRNA directed against PER2 as indicated. γH2AX ChIP-seq track (+DSB) are also shown (dark red). Examples of DSB clustering are shown (arrows). The magnification shows an individual DSB-DSB clustering event increasing after PER2 depletion.

**e**. Intrachromosomal DSB interactions shown as aggregate peak analysis (APA) plotted on a 200kb window (10kb resolution) before (-DSB) and after DSB (+DSB) induction in control (CTRL, upper panel) and PER2 (bottom panel) siRNA-transfected DIvA cells, for the eighty best-induced AsiSI-DSBs (left) or eighty random sites (right). FC: Fold change calculated between the central pixel and a square of 3x3 pixels in the bottom left corner of the matrix. APAs show an increased signal at the center (DSB-DSB interaction) after DSB induction only for DSB sites (FC=1.90, compared to 0.64 for random sites). The signal increases after PER2 depletion (FC=2.95, compared to 1.90 for the control siRNA).

**f**. Endogenous DSB interchromosomal interactions shown as aggregate peak analysis (APA) plotted on a 1Mb window (50kb resolution) in control (CTRL) and PER2 siRNA-transfected DIvA cells for 1,206 regions. FC: Fold change calculated between the central pixel and a square of 3x3 pixels on the bottom left corner of the matrix. APAs show an increased signal at the center (DSB-DSB interaction) after PER2 depletion (FC= 4.00, compared to 3.82 for the control siRNA).

**g**. Etoposide-induced DSB clustering in U2OS cells transfected with control (CTRL) and PER1+PER2 (PER1+2) siRNAs. Quantification of the cluster-positive cells *versus* cluster negative cells after DSB induction (>8000 cells counted for each condition), based on γH2AX staining (number and area of foci in each nucleus) using high-content microscopy (see methods). Percentages of cells in each category are indicated. Mean and SEM are shown for n=4 biological replicates. *P,* paired t-test (two-sided).

**h**. Genomic tracks of the γH2AX ChIP-seq after DSB induction (dark red), and of Chromosomal Eigen vectors (CEV) obtained by the PC1 of PCA analyses performed on differential (+DSB/-DSB) Hi-C matrices. CEV obtained for the Hi-C performed in control (CTRL, grey) and PER2 (cyan) siRNA-transfected DIvA cells, as well as upon DNA-PK inhibition (DNAPKi, black) are shown. D-compartment (D-Comp) are positive value, while negative values are non-D. Both siRNAs show D-compartment formation as already described in transfected DIvA cells (chromosomes 1 and 17 are shown).

**Extended Data Figure 8: Synchronization of circadian rhythm in human cells shows a rhythmic response to DSBs.**

**a**. Measure of PER2 and BMAL1 cDNA levels in circadian rhythm synchronized DIvA and U2OS cells after dexamethasone treatment. RT-qPCR results were normalized to GAPDH, TBP and 18S and to the average. n=3 biological replicates per cell line are shown.

**b**. Western Blots of PER2, BMAL1 and CRY2 in circadian rhythm synchronized DIvA cells after dexamethasone treatment (n=2).

**c**. γH2AX staining in circadian rhythm synchronized DIvA cells following DSB induction (2h) at 28h and 40h after dexamethasone (Dex) treatment. A representative experiment is shown.

**d.** γH2AX foci intensity, measured by high content microscopy, in U2OS cells after etoposide-induced TC-DSBs, in G1, S and G2 phases at different time points after dexamethasone (Dex) treatment. The heatmap representing PER2 cDNA levels (PER2 exp level) all along circadian rhythm synchronization is shown on the right. Data are normalized to the average. Mean and SEM (n=3 biological replicates; >7000 nuclei acquired per condition) are shown. *P*, paired t-test (two-sided) was computed using high PER time points (28h, 30h, 32h) compared to low PER time points (40h, 42h, 44h).

**e**. RT-qPCR performed before (−DSB) and after (+DSB) DSB induction in circadian rhythm synchronized DIvA cells at 28h and 40h after dexamethasone (Dex) treatment, or in asynchronous DIvA cells (No Dex). cDNA levels of five genes carrying TC-DSBs (either in their gene body or promoter) were measured by qPCR and normalized to GAPDH and TBP and to the untreated sample (−DSB). Mean and SEM are shown for n=4 biological replicates. *P*, paired t-test (two-sided).

**f**. RAD51 foci intensity, measured by high content microscopy, in circadian rhythm synchronized DIvA cells after DSB induction. The heatmap representing PER2 cDNA levels at different time points after dexamethasone treatment is indicated on the right. Data were normalized to the average. Mean and SEM are shown for n=3 biological replicates. *P*, paired t-test (two-sided) was computed using high PER time points (28h, 30h, 32h) compared to low PER time points (40h, 42h, 44h).

**g**. Model. A two-arm control of HR repair during the 24h circadian cycle. Maximum PER1/2 expression is observed before noon in diurnal primates, while CRY1 expression peaks at sunset. The absence of CRY1 combined with the presence of PER complex ensures that TC-DSBs are resected and further repositioned to the NE to complete HR. At nightfall, resection drops at its minimum, and the absence of the PER complex further precludes TC-DSB tethering at the NE and RAD51 assembly, resulting in minimum HR during the night.

### Extended Data Tables

#### Extended Data Table 1: Oligonucleotides and AsiSI-DSB sites

| SIRNAS USED IN THIS STUDY | | | |
|---|---|---|---|
| **Gene Symbol** | **Gene ID** | **Gene Accession** | **Pool Catalog Number** |
| ***TC-DSBR-focused SMARTpool siRNA library (siGENOME from Dharmacon)*** | | | |
| CTRL | | Non-targeting Control #2 | D-001206-13 |
| SETX | 23064 | NM_015046 | M-021420-01 |
| AHCTF1 | 25909 | NM_015446 | M-013961-01 |
| ALB | 213 | NM_000477 | M-010989-01 |
| ALDH18A1 | 5832 | NM_001017423 | M-006785-01 |
| ALDOA | 226 | NM_000034 | M-010376-01 |
| ALYREF | 10189 | NM_005782 | M-012078-01 |
| ATAD3A | 55210 | NM_018188 | M-008191-01 |
| ATP5A1 | 498 | NM_004046 | M-017064-01 |
| ATP5C1 | 509 | NM_005174 | M-012705-01 |
| BAG2 | 9532 | NM_004282 | M-011961-01 |
| BAZ1B | 9031 | NM_032408 | M-006901-02 |
| BCLAF1 | 9774 | NM_001077441 | M-020734-01 |
| CCAR1 | 55749 | NM_018237 | M-013828-00 |
| CFH | 3075 | NM_001014975 | M-007920-02 |
| DDX17 | 10521 | NM_001098505 | M-013450-01 |
| DDX21 | 9188 | NM_004728 | M-011919-01 |
| DDX47 | 51202 | NM_016355 | M-016299-00 |
| DDX5 | 1655 | NM_004396 | M-003774-01 |
| DEK | 7913 | NM_003472 | M-003881-02 |
| DHRS2 | 10202 | NM_005794 | M-009581-01 |
| DHX15 | 1665 | NM_001358 | M-011250-01 |
| DHX9 | 1660 | NM_001357 | M-009950-01 |
| ERAL1 | 26284 | NM_005702 | M-012709-01 |
| ERH | 2079 | NM_004450 | M-012670-01 |
| FANCI | 55215 | NM_018193 | M-022320-01 |
| FASN | 2194 | NM_004104 | M-003954-04 |
| FBL | 2091 | NM_001436 | M-011269-00 |
| GAPDH | 2597 | NM_002046 | M-004253-02 |
| GLYR1 | 84656 | NM_032569 | M-008332-01 |
| HMGA1 | 3159 | NM_145903 | M-004597-02 |
| HNRNPA0 | 10949 | NM_006805 | M-012314-01 |
| HNRNPA1 | 3178 | NM_002136 | M-008221-02 |
| HNRNPA2B1 | 3181 | NM_002137 | M-011690-01 |
| HNRNPA3 | 220988 | NM_194247 | M-019347-01 |
| HNRNPD | 3184 | NM_001003810 | M-004079-01 |
| HNRNPDL | 9987 | NM_031372 | M-012070-01 |
| HNRNPH2 | 3188 | NM_001032393 | M-013245-01 |
| HNRNPR | 10236 | NM_001102397 | M-010905-02 |

| | | | |
|---|---|---|---|
| HNRNPUL2 | 221092 | NM_001079559 | M-032789-01 |
| HRNR | 388697 | NM_001009931 | M-027450-01 |
| ILF3 | 3609 | NM_153464 | M-012442-02 |
| JUP | 3728 | NM_002230 | M-011708-00 |
| KHDRBS1 | 10657 | NM_006559 | M-020019-00 |
| KIF11 | 3832 | NM_004523 | M-003317-01 |
| KRT1 | 3848 | NM_006121 | M-012865-02 |
| KRT2 | 3849 | NM_000423 | M-011066-00 |
| KRT3 | 3850 | NM_057088 | M-012902-01 |
| KRT5 | 3852 | NM_000424 | M-011067-01 |
| KRT6B | 3854 | NM_005555 | M-012117-02 |
| KRT75 | 9119 | NM_004693 | M-019835-01 |
| KRT9 | 3857 | NM_000226 | M-011068-01 |
| LBR | 3930 | NM_002296 | M-021505-00 |
| LMNA | 4000 | NM_005572 | M-004978-01 |
| LMNB2 | 84823 | NM_032737 | M-005290-00 |
| LRRC59 | 55379 | NM_018509 | M-010669-00 |
| MAIP1 / C2orf47 | 79568 | NM_024520 | M-014355-00 |
| MGST3 | 4259 | NM_004528 | M-008382-00 |
| MRPS31 | 10240 | NM_005830 | M-012111-00 |
| MSH6 | 2956 | NM_000179 | M-019287-01 |
| MTA2 | 9219 | NM_004739 | M-008482-00 |
| MTCH1 | 23787 | NM_014341 | M-007370-01 |
| NDUFA5 | 4698 | NM_005000 | M-012000-00 |
| NDUFS8 | 4728 | NM_002496 | M-019600-00 |
| NDUFV2 | 4729 | NM_021074 | M-012589-01 |
| NOLC1 | 9221 | NM_004741 | M-019843-01 |
| NONO | 4841 | NM_007363 | M-007756-00 |
| NOP56 | 10528 | NM_006392 | M-019143-01 |
| PARP1 | 142 | NM_001618 | M-006656-01 |
| PGAM5 | 192111 | NM_138575 | M-015919-02 |
| PHB | 5245 | NM_002634 | M-010530-00 |
| PHB2 | 11331 | NM_007273 | M-018703-00 |
| PHGDH | 26227 | NM_006623 | M-009518-01 |
| PHIP | 55023 | NM_017934 | M-019291-00 |
| PIGT | 51604 | NM_015937 | M-003691-01 |
| PLEC | 5339 | NM_201378 | M-003945-03 |
| PPHLN1 | 51535 | NM_001143787 | L-021306-02-0020 |
| PPP1R10 | 5514 | NM_002714 | M-011358-00 |
| PRKDC | 5591 | NM_006904 | M-005030-01 |
| PRPF31 | 26121 | NM_015629 | M-020525-01 |
| PRRC2C | 23215 | NM_015172 | M-014078-02 |
| PSMC2 | 5701 | NM_002803 | M-008180-00 |
| PSMD1 | 5707 | NM_002807 | M-011363-00 |
| PSMD2 | 5708 | NM_002808 | M-017212-01 |
| QARS | 5859 | NM_005051 | M-008910-00 |
| RBBP4 | 5928 | NM_005610 | M-012137-00 |
| RBM22 | 55696 | NM_018047 | M-021186-00 |
| RBM25 | 58517 | NM_021239 | M-021976-01 |

| | | | |
|---|---|---|---|
| RBM3 | 5935 | NM_006743 | M-018969-01 |
| RBM39 | 9584 | NM_004902 | M-011965-00 |
| RFC1 | 5981 | NM_002913 | M-009290-00 |
| RFC4 | 5984 | NM_002916 | M-008691-01 |
| RHBDD2 | 57414 | NM_001040456 | M-020778-01 |
| RPL23 | 9349 | NM_000978 | M-013634-02 |
| RPL28 | 6158 | NM_000991 | M-011145-00 |
| RPL29 | 6159 | NM_000992 | M-012979-00 |
| RPS4X | 6191 | NM_001007 | M-011138-01 |
| RSF1 | 51773 | NM_016578 | M-020374-00 |
| RTCB | 51493 | NM_014306 | M-017647-01 |
| RUVBL1 | 8607 | NM_003707 | M-008977-01 |
| RUVBL2 | 10856 | NM_006666 | M-012299-00 |
| S100A8 | 6279 | NM_002964 | M-011770-02 |
| SEC61A1 | 29927 | NM_013336 | M-021503-01 |
| SF3B3 | 23450 | NM_012426 | M-020085-01 |
| SFPQ | 6421 | NM_005066 | M-006455-02 |
| SLTM | 79811 | NM_001013843 | M-014434-01 |
| SMARCD2 | 6603 | NM_003077 | M-011393-02 |
| SNRPD2 | 6633 | NM_004597 | M-013617-00 |
| SPIN1 | 10927 | NM_006717 | L-019983-02-0020 |
| SPIN2B | 474343 | NM_001006681 | L-034890-02-0020 |
| SPIN3 | 169981 | NM_001010862 | L-024685-00-0020 |
| SPOUT / C9orf114 | 51490 | NM_016390 | M-010900-00 |
| SPTAN1 | 6709 | NM_003127 | M-009933-01 |
| SRP14 | 6727 | NM_003134 | M-017767-01 |
| SSR4 | 6748 | NM_006280 | M-012264-01 |
| SUB1 | 10923 | NM_006713 | M-009815-02 |
| SUN2 | 25777 | NM_015374 | M-009959-00 |
| SUPT16H | 11198 | NM_007192 | M-009517-00 |
| THRAP3 | 9967 | NM_005119 | M-019907-01 |
| TIMM23B | 653252 | XM_003118952 | M-039881-02 |
| TMTC3 | 160418 | NM_181783 | M-018618-01 |
| TOP1 | 7150 | NM_003286 | M-005278-00 |
| TOP2A | 7153 | NM_001067 | M-004239-02 |
| U2AF2 | 11338 | NM_001012478 | M-012380-01 |
| U2SURP | 23350 | NM_001080415 | M-023607-01 |
| UQCRC2 | 7385 | NM_003366 | M-008334-02 |
| UTP15 | 84135 | NM_032175 | M-014796-01 |
| VCP | 7415 | NM_007126 | M-008727-01 |
| WDR43 | 23160 | NM_015131 | M-022651-01 |
| WTAP | 9589 | NM_152857 | M-017323-02 |
| ZBTB20 | 26137 | NM_015642 | M-020529-01 |
| ZBTB24 | 9841 | NM_001164313 | M-006915-01 |

***Other SMARTpool siRNAs used (siGENOME from Dharmacon)***

| | | | |
|---|---|---|---|
| CTRL | | Non-targeting Control #2 | D-001206-13 |
| SETX | 23064 | NM_015046 | M-021420-01 |
| PER1 | 5187 | NM_002616 | M-011350-00 |

| | | | |
|---|---|---|---|
| PER2 | 8864 | NM_022817 | M-012977-01 |
| ARNTL / BMAL1 | 406 | NM_001030272 | M-010261-00 |
| CRY2 | 1408 | NM_021117.5 | M-014151-01 |
| CLOCK | 9575 | NM_001267843 | M-008212-00 |
| NUP153 | 9972 | NM_001278209 | M-005283-00 |
| RNF4 | 6047 | NM_001185009.3 | M-006557-03 |

***Other siRNAs used (from Eurogentec)***

| Gene Symbol | Gene ID | Sense Strand Sequence |
|---|---|---|
| Ctrl1 | | CAUGUCAUGUGUCACAUCU |
| SUN1 | 23353 | GGACGUGCAAGUCAGAGAA |
| SUN2 | 25777 | CGAGCCTATTCAGACGTTTCA |
| RBBP8/CtIP | 5932 | GCUAAAACAGGAACGAAUC |

**AsiSI-DSB SITES USED FOR qPCR IN THIS STUDY**

| Name | DSB repair | DIvA AsiSI site | Position (hg19) |
|---|---|---|---|
| **DSB1** | HR | SITE657 | chr22:38864101 |
| **DSB2** | HR | SITE379 | chr17:57184296 |
| **DSB3** | HR | SITE600 | chr20:30946312 |
| **DSB4** | HR | SITE526 | chr1: 89458596 |
| **DSB5** | NHEJ | SITE72 | chr11: 24518475 |
| **DSB6** | NHEJ | SITE622 | chr21: 33245518 |
| **DSB7** | HR | SITE343 | chr17:5390220 |

**qPCR PRIMERS USED IN THIS STUDY**

| **Name** (position/AsiSI) | Forward primer sequence | Reverse primer sequence | Panels |
|---|---|---|---|
| ***Primers for ChIP-qPCR*** | | | |
| **Ctrl** genomic locus | AGCACATGGGATTTTGCAGG | TTCCCTCCTTTGTGTCACCA | 2e, all to normalize |
| **DSB1** (76-230bp) | CCGCCAGAAAGTTTCCTAGA | CTCACCCTTGCAGCACTTG | 2e,3c-d,3j, 4c-d,6c-d, 6f,E5b-c, E6f,E6l |
| **DSB1** (736-903bp) | GGGTATGGAGCTGCCTCTAA | GACAAAGATGGCTGGAGGAG | 2b,6b |
| **DSB2** (242-329bp) | ATCTCTTCAGGTCCAGCGCC | CCACCGTTCGCCCTACTTCT | 3c-d,4c-d, 6c-d,6f, E5b,E6l |
| **DSB2** (1552-1660bp) | TCCCAAGATCGCTTTGAACTAC | CCAGAAATCTGAATTGCTGCC | 2b,6b |
| **DSB3** (685-899bp) | CCTAGCTGAGGTCGGTGCTA | GAAGAGTGAGGAGGGGGAGT | 3c-d,3j,4c-d, 6c-d,6f,E5b |
| **DSB3** (1266-1420bp) | GTCAGTATGGCCCCAGAGTC | ACGGCTGATGGACTTAGACG | 6b |
| **DSB4** (100-309bp) | GATTGGCTATGGGTGTGGAC | CATCCTTGCAAACCAGTCCT | 2e,3c-d,4c-d, 6c-d,6f, E5b,E6f |
| **DSB5** (124-321bp) | TGAAGATACTCCTCGCTCCCAG | GCTGAATTTCATGCTGCTGCC | 3c-d,3j,4c, 6c-d,6f, E5b,E6f |
| **DSB6** (771-967bp) | GGCACCTCAACAGGTAGCAT | GCCTCTCTTCGATGCTTTTG | 3j,6f,E5b-c |

| | | | |
|---|---|---|---|
| **LAD#1** | AGCACATGGGATTTTGCAGG | TTCCCTCCTTTGTGTCACCA | E4f, E5a |
| **LAD#2** | AGAAACTCACAAACTCGCTGGA | GGAAATGATGGTGTAGGGTGC | E4f, E5a |
| ***PER1* Promoter** | TCTCCCTCTCTCCTCCCTTCC | CACTAAGGTCAGGGCTGTGTCAC | 2e |
| ***BTG2* Promoter** | CAGGGTAACGCTGTCTTGTGG | CTCAGGAGGCTGGAGAGGAAG | 2e |
| ***Primers for RT-qPCR*** | | | |
| ***GAPDH*** | GACAGTCAGCCGCATCTTCT | GTTAAAAGCAGCCCTGGTGAC | 4a-b,6h, E8a,E8e |
| ***TBP*** | TTCGGAGAGTTCTGGGATTG | GAAAATCAGTGCCGTGGTTC | 4a-b,6h, E8a,E8e |
| ***18S*** | GGCGGCTTTGGTGACTCTA | GCCTTCCTTGGATGTGGTAG | 4a-b,6h, E8a,E8e |
| ***PER1*** | CAGGATGTGGGAGTCTTCTATGG | CTTGGTCACATACGGGGTTAGG | E2a |
| ***PER2*** | AAGCCACGAGAATGAAATCCG | CCGCCCTTTCATCCACATC | E2a,E8a |
| ***ARNTL / BMAL1*** | ACCGCAAACGGAAAGGCAG | TATCCCGACGCCGCTTTTC | E8a |
| ***ASXL1*** | CAGGAAAGAAGCCAGCACAG | CCCTATTCCCCAACCCCATT | 4a-b |
| ***KLF7*** | ATGACAGGAGGTGGTTGGAG | TGCTCTTTCCTGGAAGTCGT | 4a-b,E8e |
| ***MIS12*** | CAGGCCGTTGAACAGGTTAT | TCAGCTGCAAAAACAGTTGC | 4a-b,E8e |
| ***RBMXL1*** | GTGCTCCACTTACACGAGGG | TGCCAACCCTGTCACAACTT | 4a-b,E8e |
| ***SRSF6*** | CCTGATTGGCCTTGGCTTTT | ACTGCTGTATCCACCTCCAC | 4a-b,E8e |
| ***TRIM37*** | CTCCCCGAGCCTTGATACAT | GCAGTCCTTCCAGATGACCT | 4a-b,E8e |
| ***RNF19B*** | CATCAAGCCATGCCCACGAT | GAATGTACAGCCAGAGGGGC | 6h |
| ***PLK3*** | GCCTGCCGCCGGTTT | GTCTGACGTCGGTAGCCCG | 6h |
| ***PPM1D*** | CTTGTGAATCGAGCATTGGG | AGAACATGGGGAAGGAGTCA | 6h |
| ***SLC9A1*** | TGTTCCTCAGGATTTTGCGG | ATGAAGCAGGCCATCGAGC | 6h |
| ***LPHN2*** | CGATTTGAAGCAACGTGGGA | TGATACTGGTTGGGGAAGGG | 6h |
| ***UTP18*** | TCCTACTGTTGCTCGGATCTC | CGTGGCTAAAACTTCTTCCCC | 6h |
| ***Primers for Resection assays*** | | | |
| **DSB1** (132-362bp) | ACCATGAACGTGTTCCGAAT | GAGCTCCGCAAAGTTTCAAG | E6i-j |
| **DSB7** (1014-1228bp) | TTCCCCTCTCGACAGTTTCC | AGAAGGGAGTGCTGCTTCAT | E6i-j |
| ***Primers for Translocation analyses*** | | | |
| ***t(MIS12;TRIM37)*** | GACTGGCATAAGCGTCTTCG | TCTGAAGTCTGCGCTTTCCA | 1b,2c,5e-h, E1d,E1h,E2i |
| ***t(LINC00271;LYRM2)*** | GGAAGCCGCCCAGAATAAGA | TCCATCTGTCCCTATCCCCAA | E1h,E2i |
| **Norm1** (chr1) | AGCACATGGGATTTTGCAGG | TTCCCTCCTTTGTGTCACCA | 1b,2c,5e-h, E1d,E1h,E2i |

| | | | |
|---|---|---|---|
| **Norm17** (chr17) | ACAGTGGGAGACAGAAGAGC | CTCCATCATCGCACCCTTTG | 1b,2c,5e-h, E1d,E1h,E2i |
| ***Primers for Cloning experiments*** | | | |
| **AsiSI-F** | GATTTGAGGATGTATAAAAGCGGATCCTACCCATACGATGTTCCAGATTACGCTAGC | | |
| **AsiSI-R** | GCAGAGAGAAGTTCGTGGCACGCGTCAACATCACCTGGTCGATAAGGTGGC | | |
| **mAID-F** | CTCACTCCCGGGTCCAAGGAGAAGAGTGCTTGTCCTAAAG | | |
| **mAID-R** | CTCTCGAATTCCACGCGTCCTAGGTGGATCCGCTTTTATACATCCTCAAATCGATTTTCCTC | | |
| **P2A-F** | CGCGTGCCACGAACTTCTCTCTGCTAAAGCAAGCTGGCGATGTGGAAGAAAATCCCGGGCCTG | | |
| **P2A-R** | AATTCAGGCCCGGGATTTTCTTCCACATCGCCAGCTTGCTTTAGCAGAGAGAAGTTCGTGGCA | | |
| **Puro-F** | GTGGAAGAAAATCCCGGGCCTGAATTCATGACAGAGTACAAGCCAACAGTGAGAC | | |
| **Puro-R** | AATTTTGTAATCCAGAGGTTGATTATCATATGCTAGGCACCTGGCTTTCTGGTCATGCAC | | |
| **T2A-F** | CCGGAGGGCAGAGGAAGTCTGCTAACATGCGGTGACGTCGAGGAGAATCCTGGACCCGGGTCACTCA | | |
| **T2A-R** | CGCGTGAGTGACCCGGGTCCAGGATTCTCCTCGACGTCACCGCATGTTAGCAGACTTCCTCTGCCCT | | |
| **TIR1-F** | CGATCACGAGACTAGCCTCGAGGTTTCCTAGGCGTACGCGCCACCATGACG | | |
| **TIR1-R** | CTGCCCTCCGGAGATCCCGTAGTTTCGAGGTCTTCCTCCGAGATGAGTTTCTG | | |
| **F1_fwd** | TAGCATATGATAATCAACCTCTGG | | |
| **F1_rev** | TAGCATATGATAATCAACCTCTGG | | |
| **F2_fwd** | GCCTCACTGATTAAGCATTGGTAACTGTCAGACCAAGTTTACTC | | |
| **F2_rev** | AGAACAAGAGCCCTGAACCGA | | |
| **F3_fwd** | TCCCTCGGTTCAGGGCTCTTGTTCTTATCAGCTGCGAAGGCTTC | | |
| **F3_rev** | GAATTCAGGCCCGGGATTTTC | | |
| **BSD_fwd** | GGAAGAAAATCCCGGGCCTGAATTCATGGCCAAGCCTTTGTCTC | | |
| **BSD_rev** | TCCAGAGGTTGATTATCATATGCTAGCCCTCCCACACATAACC | | |

**Extended Data Table 2: primary Antibodies used in this study**

| Target Protein | Host | Reference | Application and quantity |
|---|---|---|---|
| 53BP1 | Mouse | BD biosciences 612522 | PLA (1:1000) |
| BMAL1 | Rabbit | Abcam ab3350 | ChIP (2 µg) WB (1:500) |
| CLOCK | Rabbit | Abcam ab3517 | WB (1:200) |
| CRY1 | Rabbit | Bethyl A302-614A | WB (1:1000) |
| CRY2 | Rabbit | Proteintech 13997-1-AP | WB (1:500) |
| DDX5 | Rabbit | Bethyl A300-523A | ChIP (5 µg) |
| DDX5 | Mouse | Sigma 05-850 | WB (1:5000) |
| DDX17 | Rabbit | Proteintech 19910-1-AP | ChIP (5 µg) WB (1:1000) |
| GAPDH | Mouse | Sigma MAB374 | WB (1:10 000) |
| γH2AX | Rabbit | Abcam ab81299 | ChIP (1 µg) |
| γH2AX | Mouse | Sigma JBW301 | IF (1:1000) PLA (1:2500) |
| LAMIN B1 | Rabbit | Abcam ab16048 | ChIP (5 µg) PLA (1:4000) |
| MYOSIN | Rabbit | Sigma M3567 | WB (1:2000) |
| NONO | Rabbit | Proteintech 11058-1-AP | WB (1:1000) |
| NUP153 | Rabbit | Abcam ab84872 | ChIP (0.5 µg) WB (1:500) |
| PCNA | Mouse | Santa Cruz sc-56 | WB (1:1000) |
| PER2 | Rabbit | Proteintech 20359-1-AP | ChIP (5 µg) WB (1:500) |
| PER2 | Rabbit | Abcam ab179813 | WB (1:1000) |
| RAD51 | Rabbit | Santa Cruz sc8349 | ChIP (3 µg) IF (1:200) |
| RNF4 | Goat | Bio-Techne AF7964 | WB (1:1000) |
| RPA32/RPA2 | Rabbit | Abcam ab10359 | ChIP (2 µg) |
| SUN1 | Rabbit | Sigma HPA008346 | ChIP (5 µg) IF (1:500) |
| SUN1 | Rabbit | Abcam ab124770 | WB (1:1000) |
| SUN2 | Rabbit | Abcam ab124916 | ChIP (5 µg) IF (1:1000) WB (1:1000) |
| α-Tubulin | Mouse | Sigma T6199 | WB (1:10 000) |
