## Supplemental Movie Legends for "Circadian PERIOD proteins regulate TC-DSB repair through anchoring to the nuclear envelope"

1. *MCD, Centre de Biologie Intégrative (CBI), CNRS, Université de Toulouse, UT3, 31062 Toulouse, France*
2. *BigA, NGS data analysis facility, Centre de Biologie Intégrative (CBI), CNRS, Université de Toulouse, UT3, 31062 Toulouse, France*
3. *Institut de Pharmacologie et de Biologie Structurale (IPBS), Université de Toulouse, CNRS, Université Toulouse III - Paul Sabatier (UT3), Toulouse, France*
4. *Equipe Labellisée Ligue Contre le Cancer 2018, Toulouse, France*
5. *LITC Core Facility, Centre de Biologie Intégrative (CBI), CNRS, Université de Toulouse, UT3, 31062 Toulouse, France*
6. *Lead author*

**Running title:** Circadian regulation of TC-DSB repair

### Supplementary Video Legends 1-4

#### Supplementary Video Legends

**Supplementary Movie 1: Close contact between a DSB and the nuclear envelope.**

Live imaging showing close contact between a 53BP1-GFP focus (white) and mcherry-Lamin B1 (purple) at the nuclear envelope during DSB induction in DIvA cells. Single Z slice from 3D Time-series super-resolved RIM. Movie of 39min with 40sec intervals starting ~5h30 after DSB induction. The scale bar (1µm) and the time scale are indicated.

**Supplementary Movie 2: Close contact between DSBs and the nuclear envelope.**

Live imaging showing close contact between 53BP1-GFP foci (white) and mcherry-Lamin B1 filament (purple) during DSB induction in DIvA cells. 3D view (VTK) from 3D Time-series super-resolved RIM. Movie of 34min with 40sec intervals starting ~4h after DSB induction. The scale (from 0 to 7 µm) on the X- and Y-axis and the time scale are indicated.

**Supplementary Movie 3: Close contact between DSBs and the nuclear envelope.**

Live imaging showing close contact between 53BP1-GFP foci (white) and mcherry-Lamin B1 filament (purple) during DSB induction in DIvA cells. 3D view (VTK) from 3D Time-series super-resolved RIM. Movie of 24min with 40sec intervals starting ~4h after DSB induction. The scale (from 2 to 8 µm) on the X- and Y-axis and the time scale are indicated.

**Supplementary Movie 4: Close contact between a DSB and the nuclear envelope.**

Live imaging showing close contact between a 53BP1-GFP focus (white) and mcherry-Lamin B1 (purple) at the nuclear envelope during DSB induction in DIvA cells. Single Z slice from 3D Time-series super-resolved RIM. Movie of 39min with 40sec intervals starting ~5h30 after DSB induction. The scale bar (1µm) and the time scale are indicated.
